# Neuronal activity triggers widespread changes in RNA stability

**DOI:** 10.1101/2025.09.05.674581

**Authors:** Erin E. Duffy, Inés L. Patop, Shelly Kalaora, Elena G. Assad, Shon A. Koren, Lisa Traunmüller, Sebastian Krüttner, Naeem S. Pajarillo, Benjamin Finander, Zachary Barsdale, Mia M. Macias, Min Yi Feng, Joao A. Paulo, Eric C. Griffith, Brian T. Kalish, Steven P. Gygi, L. Stirling Churchman, Michael E. Greenberg

**Affiliations:** Department of Neurobiology, Harvard Medical School, Boston, MA, USA; Department of Genetics, Blavatnik Institute, Harvard Medical School, Boston, MA, USA; Department of Cell Biology, Harvard Medical School, Boston, MA, USA; Program in Neuroscience, Harvard Medical School, Boston, MA, USA; Biology of Adversity Project, Broad Institute of MIT and Harvard, Cambridge, MA, USA; Department of Molecular Genetics, University of Toronto, Toronto, ON M5G 1A8, Canada; Division of Newborn Medicine, Boston Children’s Hospital, Boston, MA, USA; F.M. Kirby Neurobiology Center, Boston Children’s Hospital, Boston, MA USA

## Abstract

Neuronal activity shapes brain development and refines synaptic connectivity in part through dynamic changes in gene expression. While activity-regulated transcriptional programs have been extensively characterized, the holistic effects of neuronal activity on the full RNA life cycle remain relatively unexplored. Here, we show that neuronal activity influences multiple stages of RNA metabolism *in vitro* and *in vivo*. Among these, RNA stability emerges as a previously underappreciated regulator of gene expression, exerting a stronger influence than transcription on total RNA levels for ∼15% of activity-dependent genes. We go on to profile 3′UTR mRNA motifs that are sufficient to modulate activity-dependent mRNA stability and employ machine learning to identify the neuronal-specific RNA-binding protein HuD as a key regulator of activity-dependent mRNA stabilization. We demonstrate that HuD shapes activity-dependent mRNA abundance of hundreds of transcripts in both soma and distal neuronal processes and that neuronal activity drives the reorganization of HuD-interacting proteins, thereby stabilizing HuD-bound mRNAs and directing them into translationally active granules. Finally, we find that many variants associated with autism spectrum disorder (ASD) and other neurodevelopmental disorders disrupt or promote aberrant activity-dependent changes in mRNA stability. These findings reveal mRNA stability as a widespread mechanism of stimulus-responsive gene regulation in neurons with direct implications for the understanding of neurodevelopmental disorders.

## INTRODUCTION

In all stages of life, neurons must convert transient synaptic activity into sustained changes in connectivity. For example, during early postnatal life and into adulthood, sensory experience drives the refinement of topographic sensory maps.^1–4^ Additionally, neuronal activity facilitates the synaptic plasticity that underlies learning, memory, and behavior.^5,6^ These changes in neuronal connectivity are accomplished in part through the dynamic regulation of gene expression.^7^ While it is well appreciated that each step of the RNA life cycle, including RNA transcription, processing (e.g., splicing, polyadenylation, nuclear export, and localization), degradation, and translation, contributes to a dynamic regulatory network that controls RNA and protein abundance,^8,9^ our understanding of the coordinated effect of acute external stimuli on this regulatory network remains incomplete. This coordination is particularly critical in neurons, which face the unique challenge of modulating gene expression across elaborate polarized structures that receive distinct types of synaptic input (e.g. somatic inhibitory synapses versus dendritic excitatory synapses).^10,11^ Historically, neuronal activity-dependent changes in gene expression have been studied in the context of signaling to the nucleus to promote new gene transcription,^7^ as well as localized translation within dendrites and axons.^12^ In contrast, the contribution of RNA stability regulation to activity-dependent responses remains largely unexplored.

Messenger RNA half-lives vary widely within cells, ranging from as short as five minutes to as long as two weeks, which significantly influences corresponding protein abundance, and disruptions in mRNA stability can lead to substantial alterations in gene expression.^13^ In mammalian cells, mRNA stability regulation is mediated via a network of factors, including RNA-binding proteins (RBPs), microRNAs (miRNAs), and chemical modifications such as N6-methyladenosine (m^6^A).^14^ RBPs recognize specific RNA sequence and/or structural motifs and can either stabilize or destabilize mRNAs through various mechanisms, including modulation of RNA secondary structure, protection from or recruitment of RNA decay factors, and sequestration into membraneless cytoplasmic compartments such as P-bodies and stress granules.^15–18^ Furthermore, many RBPs exhibit tissue-specific or activity-dependent expression patterns, suggesting specialized roles in different cellular contexts.^19^ The combinatorial action of RBPs, miRNAs, and RNA modifications creates a complex regulatory network that fine-tunes mRNA stability in response to different cellular conditions. The ability of RBPs to control RNA stability in specific cellular regions may allow for rapid and localized gene regulation in response to external cues. However, despite the potential importance of RNA stability regulation for stimulus-responsive gene expression, the specific mechanisms by which neuronal activity or other external stimuli influence RNA turnover dynamics in neurons, and the scope of this regulation across the neuronal transcriptome, remain poorly understood.

Here, we characterized how neuronal activity affects each stage of the mRNA life cycle and found that activity-dependent changes in mRNA stability play a central role in shaping gene expression in both primary neuronal cultures and in the mouse hippocampus. We observed activity-dependent changes in RNA stability, both stabilization and destabilization, for thousands of transcripts, and these changes were tightly linked to protein abundance. Notably, for roughly 10% of activity-dependent genes, changes in RNA stability had a greater impact than transcription on total mRNA levels. *In vivo* massively parallel reporter assays and machine learning revealed the neuronal RBP HuD as a key regulator of activity-stabilized mRNAs in both soma and distal neuronal processes. We found that HuD interacts with an activity-sensitive ensemble of RBPs, including proteins involved in RNA granule formation, suggesting a mechanism of selective mRNA stabilization via subcellular relocalization. Activity-dependent transcripts and HuD-bound mRNAs are enriched for ASD risk genes, and 3′UTR variants linked to neurodevelopmental disorders disrupt both basal and activity-dependent RNA stability, highlighting the importance of this pathway for neuronal function and development. Together, these findings highlight RNA stability as a widespread and consequential mechanism by which neurons orchestrate activity-dependent gene expression programs to regulate neuronal development and plasticity.

## RESULTS

### Measuring activity-dependent RNA dynamics

To measure activity-dependent changes in the RNA life cycle transcriptome-wide, we leveraged an RNA metabolic labeling strategy that allow us to measure RNA transcription, intron half-life, and mRNA half-life using the non-canonical nucleoside 4-thiouridine (s^4^U) (**Fig. 1A left panel**).^20–24^ Cultured primary mouse cortical neurons were treated at day in vitro 7 (DIV7) with an elevated level of extracellular potassium chloride (KCl) in a time course from 15 min to 8 h. Elevated KCl reduces the potassium gradient, leading to membrane depolarization, opening of L-type voltage-gated calcium channels, and a subsequent influx of calcium that triggers the activation of activity-dependent gene programs.^25–28^ The robust response generated by KCl treatment has served as a powerful tool to identify activity-dependent gene programs that are also activated *in vivo* in response to sensory stimuli.^29–32^ Newly transcribed RNAs were labeled with s^4^U for 15 min at the end of KCl stimulation to achieve a snapshot of the nascent transcriptome, and both total RNA and biochemically enriched s^4^U-RNA fractions from each time point of membrane depolarization were analyzed via high-throughput sequencing. To test whether activity-dependent changes in total RNA levels lead to meaningful changes in protein abundance, we also carried out ribosome profiling to quantitatively measure active translation in response to neuronal activity and mass spectrometry-based whole-cell quantitative proteomics with Tandem mass tag (TMT) to capture resulting changes in relative protein abundance (**Fig. 1A center and right panel, Fig. S1, S2**).

**Figure 1:**
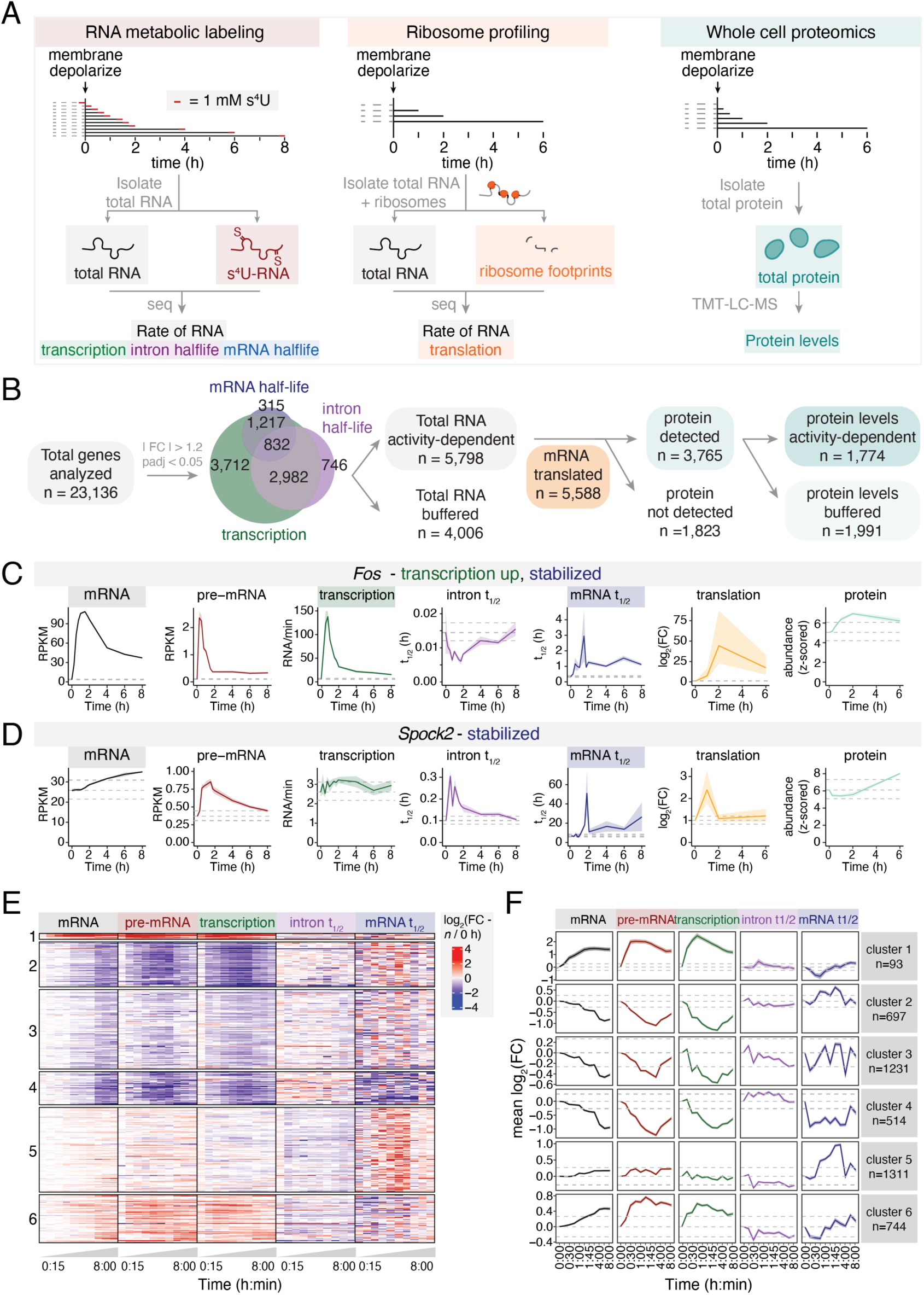
Measuring activity-dependent RNA dynamics. (A) Schematic of s^4^U RNA metabolic labeling, ribosome profiling, and proteomics in membrane-depolarized primary hippocampal neurons. Metabolic labeling: *n*=4 independent biological replicates (0, 15, 30, 45 min; 1, 1.5, 1.75, 2, 4, 6, 8 h membrane depolarization). Ribosome profiling: *n*=4 independent biological replicates (0, 1, 2, 6 h membrane depolarization). Proteomics: *n*=3 independent biological replicates (0, 15, 30 min; 1, 2, 6 h membrane depolarization). (B) Flow chart of activity-dependent genes regulated at each stage of the RNA life cycle. (C&D) Line plots of synthesis, processing, degradation, and translation, as well as total RNA, pre-mRNA, and protein abundance, for select activity-stabilized genes. Dark line = mean, shading = ± SEM, dashed lines = mean in unstimulated neurons ±1.2-fold change. (E) Heatmap of log_2_(fold change (FC)) in rates for all activity-regulated genes at different times of membrane depolarization. (F) Metaplots of mean fold change

Rates of RNA transcription, intron half-life (i.e. the rate of intron excision from pre-mRNA) and mRNA half-life were calculated for each gene using our time series of paired total-RNA and s^4^U-RNA sequencing data via the computational pipeline INSPEcT.^33^ To validate that the model yields expected rates of RNA turnover, we first assessed mRNA half-lives in unstimulated neurons. As expected, fast-turnover RNAs (top 10%, *n*=975) were enriched for transcripts that encode transcription factors (Results from gProfiler2 analysis: DNA-templated transcription, *P* < 10^−5^), while slow-turnover RNAs (bottom 10%, *n*=975) were enriched for transcripts that encode synaptic proteins (Results from gProfiler2 analysis: synaptic vesicle cycle, *P* < 10^−8^, chemical synaptic transmission, *P* < 10^−8^) (**Fig. S3A**). Together, these observations suggest that our approach accurately captures physiologically relevant mRNA half-lives in neurons, providing a robust foundation for measuring stimulus-induced changes in RNA dynamics.

### Widespread activity-dependent modulation of RNA stability is a key determinant of RNA output

We next examined how membrane depolarization affects the neuronal RNA life cycle. Across the transcriptome, we identified 8,743 genes with altered transcription rates, 4,723 genes with activity-dependent changes in intron half-life, and 2,527 genes with changes in mRNA half-life (**Fig. 1B**, **Table 1**). In total, 5,798 genes displayed significant activity-dependent changes in steady-state RNA levels (Fig. 1B, S3B). These transcripts often showed coordinated regulation across multiple stages of the RNA life cycle, with buffering between synthesis and decay in some cases masking underlying dynamic changes (e.g., increased transcription offset by increased degradation; **Fig. 1B, S3A**). At the protein level, we detected 3,765 proteins, of which 1,774 (47%) showed significant activity-dependent changes in expression (**Fig. S3B**). These results indicate that neuronal activity induces widespread remodeling of protein abundance alongside coordinated regulation of RNA transcription, intron half-life, and mRNA half-life, with additional layers of control (e.g., translation, protein turnover, and localization) shaping the final proteomic response.

By way of example, our metabolic labeling measurements confirmed a robust transcriptional induction of *Fos*, a well-established early response gene (ERG),^34,35^ and also demonstrate a >3-fold increase in *Fos* mRNA half-life following neuronal activity that maintains elevated *Fos* mRNA levels even after its transcription returns to near baseline levels (**Fig. 1C**). By contrast, *Spock2*, a gene that encodes a proteoglycan that regulates the neuronal extracellular matrix during CNS development,^36^ showed increased mRNA and protein abundance through an increased half-life without a significant change in *Spock2* mRNA transcription (**Fig. 1D**). Activity-dependent mRNA downregulation occurred via similarly diverse regulatory strategies. *Calb2,* a gene that encodes the protein Calretinin, a buffer of intracellular calcium levels that influences synaptic plasticity and neuronal excitability,^37^ showed activity-dependent RNA destabilization coupled with transcriptional downregulation **(Fig. S3C).** By contrast, *Cacna1h*, a gene that encodes a low-voltage T-type calcium channel that controls intrinsic neuronal excitability during development,^38^ showed a decreased RNA half-life following stimulation, without a significant change in *Cacna1h* mRNA transcription, resulting in decreased mRNA and protein abundance after 4 h of activity **(Fig. S3D).** Together, these examples illustrate that neurons shape activity-regulated gene expression through both transcriptional and post-transcriptional mechanisms, and our analysis offers a transcriptome-wide resource detailing how each step of the RNA life cycle is modulated by neuronal activity.

To assess the relative contribution of RNA stability to activity-dependent changes in mRNA and protein expression, we first clustered transcripts based on their rates of transcription, mRNA half-life, and translation, as well as changes in the total level of RNA and protein (**Fig. 1E, F, S4A, B**). This analysis revealed different combinations of transcriptional and post-transcriptional control driving activity-dependent changes in mRNA and protein expression. We observed gene groups where coordinated changes in transcription and mRNA stability either increase or reduce mRNA abundance (clusters 1 and 4, respectively; **Fig. 1E, F**). For instance, *Fos* showed both transcriptional induction and mRNA stabilization in response to neuronal activity (cluster 1, **Fig. 1C**). In contrast, other groups of genes showed opposing changes in synthesis and degradation rates that resulted in a buffering effect on the level of RNA (clusters 2 and 6, **Fig. 1E, F**). Notably, we also identified groups of genes where changes in RNA stability occur independently of a change in transcription, such as cluster 5 (**Fig. 1E, F**) which includes activity-stabilized genes such as *Spock2* and is enriched for genes that function at synapses (**Fig. S4C, Table 2**). These findings highlight that RNA stability is a widespread mechanism for activity-dependent gene regulation that can operate independently of changes in transcription, thus expanding the regulatory toolkit available to neurons for fine-tuning gene expression responses possibly within specific neuronal compartments.

We next asked whether changes in transcription, intron half-life, and mRNA half-life are required for activity-dependent changes in total mRNA abundance. For each rate, we compared models in which the rate was either fixed or allowed to vary dynamically after membrane depolarization (**Fig. 2A**). Genes were classified as dynamically regulated when a dynamic model provided a better fit than a constant-rate model (Akaike information criteria (AIC) dynamic model < constant model and Root Mean Square Error (RMSE) dynamic - constant model < −0.5). This analysis revealed that, among activity-regulated genes, 73% (n=3,401) depended on transcription, 53% (n=2,677) on mRNA half-life, and 2% (n=86) on intron half-life. (**Fig. 2B, S5A**). These categories were not mutually exclusive, as many transcripts depended on multiple regulatory modes (**Fig. S5B**). Indeed, by quantifying the relative contribution of each rate, we observed a gradient of dependencies (**Fig. 2C, S5B**): the abundance of some transcripts was almost entirely explained by transcriptional changes (e.g. *Bdnf*), others were driven predominantly by changes in RNA stability (e.g. *Cnnm2*), and still others required contributions from both transcription and stability to account for their activity-dependent expression patterns (e.g., *Gad1*) (**Fig. 2D**). Notably, for 659 transcripts (14% of activity-regulated transcripts), changes in RNA stability were the primary driver of activity-dependent mRNA abundance in neurons (**Fig. 2C**). Thus, regulated RNA stability emerges as not only a widespread feature of the neuronal activity response, but also a key determinant of total RNA levels, acting independently or in concert with transcriptional regulation.

**Figure 2:**
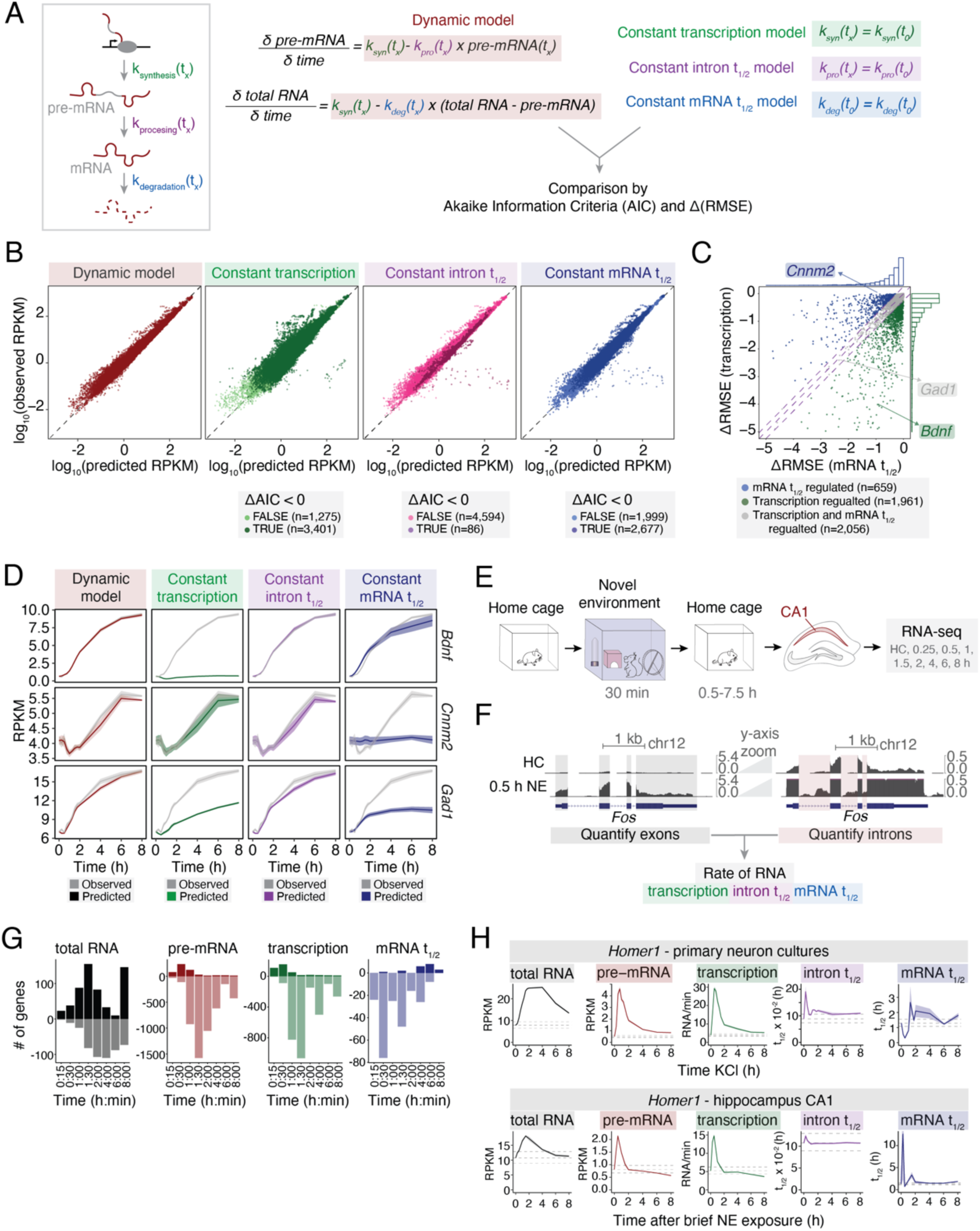
Widespread activity-dependent modulation of RNA stability is a key determinant of RNA and protein output. (A) Description of modeling strategy with a dynamic versus constant change in RNA transcription, intron half-life (t_1/2_), or mRNA t_1/2_. (B) Correlation plot between total RNA RPKM for measured RNA versus RNA levels predicted by a dynamic model (gray) or constant transcription (green) intron t_1/2_ (purple) or mRNA t_1/2_ (blue) models. Points are colored based on Akaike information criteria (AIC). ΔAIC > 0 (light green, purple, or blue, respectively) or < 0 (dark green, purple, or blue, respectively). (C) Scatterplot of Root Mean Square Error for predicted versus observed RNA (ΔRMSE) in constant transcription versus constant degradation models. More negative ΔRMSE indicates that the dynamic regulation model fits the total RNA levels better when compared with the fixed transcription or degradation model. Points are colored based on total RNA levels that are more dependent on dynamic transcription (green), mRNA t_1/2_ (blue) or both transcription and mRNA t_1/2_ (gray). (D) Modeling of genes defined by model in (A) to be dependent on activity-regulated transcription (*Bdnf*), mRNA t_1/2_ (*Cnnm2*) or both transcription and mRNA t_1/2_ (*Gad1*) for total RNA abundance.(E) Schematic of novel environment paradigm (adapted from Traunmüller et al.^32^). *n*=4-5 independent biological replicates per time point. (F) Tracks of *Fos* exon and intron expression in CA1 in homecage (HC) versus 30 min novel environment exposure (NE). Half-lives are calculated using exon vs intron abundance over time. (G) Bar plot of # genes with similar regulation between primary cultured neurons and CA1 hippocampus *in vivo*. For total RNA and pre-mRNA, upregulated and downregulated genes are plotted as positive and negative values on the y-axis, respectively. For transcription and mRNA t_1/2_ rates, genes with increased and decreased rates are plotted as positive and negative values on the y-axis, respectively. Note that there were no genes with overlapping intron t_1/2_ rates between *in vitro* and *in vivo* measurements. (H) Example rates for *Homer1* in primary neuronal cultures and CA1 hippocampus *in vivo*. *Homer1* is both transcriptionally induced and activity-stabilized after a rapid and transient destabilization. Dark line = mean, shading = ± SEM, dashed lines = mean in unstimulated neurons ± 1.2-FC.

To test whether physiologically relevant stimuli elicit similar activity-dependent changes in RNA turnover *in vivo*, we leveraged a previously generated RNA-seq dataset from the CA1 region of the mouse hippocampus following brief exposure to a novel environment,^32^ which activates hippocampal neurons in a naturalistic context and stimulates gene programs supporting learning, memory, and synaptic plasticity (**Fig. 2E**). We analyzed this dataset using a computational method that imputes kinetic rates from total RNA-seq data without requiring nascent transcript information (INSPEcT-)^39^ (**Fig. 2F**, **Table 1**). Because intronic reads are often lower in abundance in total RNA-seq data and some genes lack introns, we limited our analysis to genes where the kinetic rates correlated well *in vitro* between the computational analysis using metabolic labeling implementation and using only nascent RNA quantification from intronic reads in our primary neuron data (**Fig. S5B**). We then compared these genes at matched time points between cultured neurons (membrane depolarization) and *in vivo* in the CA1 region of the hippocampus (novel environment) and found overlapping regulation for total RNA (718 genes), pre-mRNA (3,186 genes), transcription (2,222 genes), and mRNA half-life (226 genes) across the two datasets (**Fig. 2G**). For example, *Homer1*, which encodes a synaptic scaffolding protein that binds group 1 metabotropic glutamate receptors,^40,41^ showed concordant regulation across both paradigms. This included transcriptional induction accompanied by rapid mRNA destabilization within the first hour, followed by sustained stabilization from 2-8 h post-activity (**Fig. 2H**). The presence of shared regulatory targets strongly suggests that activity-dependent RNA stability changes are physiologically relevant and occur *in vivo* in response to behaviorally salient experiences.

### Modular 3’UTR RNA sequences regulate activity-dependent RNA stability *in vivo*

Motifs that regulate RNA stability are commonly found within 3’UTRs,^42^ and recent studies suggest that discrete RNA elements within 3’UTRs are able to influence multiple aspects of RNA fate including RNA turnover and transport.^43,44^ However, the extent to which 3’UTR elements contribute to activity-dependent mRNA stability in neurons remained unknown. To address this gap, we first confirmed the ability of the 3’UTR of an example neuronal activity-stabilized gene (*Nfkbiz*) to confer strong activity-dependent regulation to a constitutive luciferase reporter gene and mapped this effect to a discrete 165 nt stability element (**Fig. S6A-B**). Having validated that a discrete 3’UTR element can drive activity-dependent mRNA stabilization, we then scaled this approach using a massively parallel reporter assay (MPRA) to identify the general principles governing the bidirectional stabilization of mRNAs in response to neuronal activity (**Fig. 3A**, **Table 3**).

**Figure 3:**
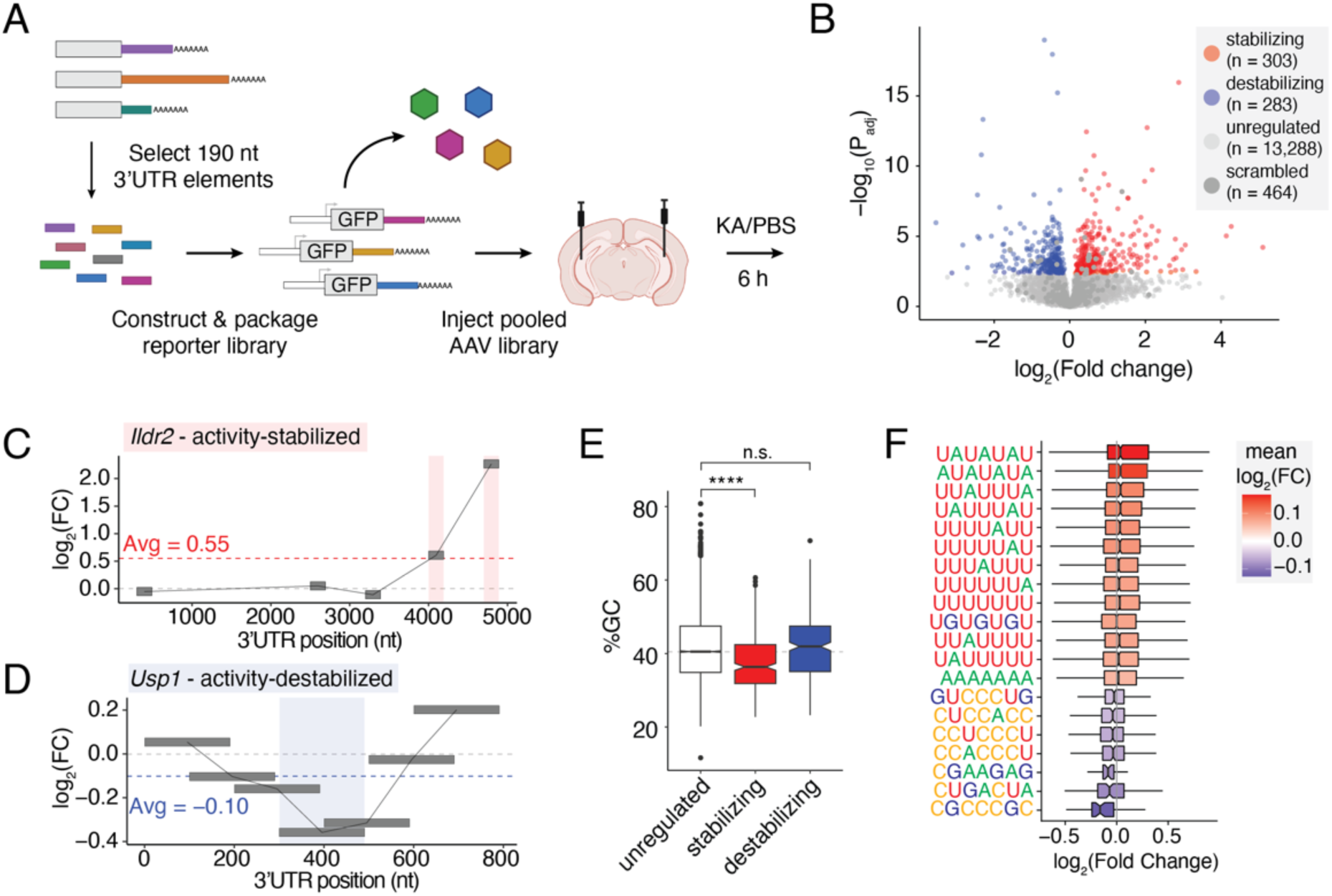
Modular 3’UTR RNA sequences are sufficient to drive activity-dependent changes in RNA stability *in vivo*. (A) Schematic of massively parallel reporter assay strategy. *n*=5-6 mice per condition (6 h KA or PBS). (B) Volcano plots of −log10(*P*adj) versus log2(FC) in reporter RNA abundance between 6 h KA and PBS. Red = activity-stabilizing elements (Padj < 0.1, log_2_(FC) > 0) blue = activity-destabilizing elements (P_adj_ < 0.1, log_2_(FC) < 0), light gray = P_adj_ > 0.1, dark gray = scrambled controls. (C&D) Log_2_(FC) in MPRA motif stability across the 3’UTR of (C) *Ildr2* (activity-stabilized gene) and (D) *Usp1* (activity-destabilized gene). X-axis shows relative position within the endogenous 3’UTR. Each box represents a 190-nucleotide motif, with position indicating its location within the 3’UTR and color/y-axis position reflecting log_2_(FC) by MPRA. Black outlines indicate statistically significant changes (Padj < 0.1). Lines connect consecutive motifs, where gaps indicate motifs excluded from the assay. Dotted horizontal lines show the average log_2_(FC) across all tested motifs for each gene (red, C; blue, D). (E) Box plot of %GC content of activity-stabilizing and -destabilizing elements. Kolmogorov-Smirnov Test (KS Test) with Dunn’s test for multiple hypotheses. **** = P_adj_ < 2.2e-16. (F) Box and whisker plot of log_2_(FC) in reporter expression for each MPRA library member containing the indicated 7-mer. All displayed 7-mers are enriched in either activity-stabilizing or -destabilizing elements relative to non-regulated sequences.

We tiled the 3’UTRs of activity-stabilized and -destabilized genes to generate 13,000 RNA elements (190 nt each), along with 1,500 elements from the 3’UTRs of activity-insensitive genes and 500 GC-matched scrambled controls derived from activity-regulated 3’UTRs. We cloned these 15,000 elements into GFP reporters driven by the non-inducible EF1ɑ promoter, and packaged the MPRA library into AAV for stereotaxic injection into mouse hippocampi. Six hours after KA-induced seizures or control PBS treatment, we microdissected the CA1 region and used high-throughput sequencing to assess differential MPRA reporter RNA expression, with PBS and KA samples separating well by PCA analysis (**Fig. S6C-E**). Critically, this approach distinguishes between constitutive regulatory elements which regulate mRNA stability in a manner that is independent of neuronal activity and 3’UTR elements that dynamically modulate mRNA stability only upon stimulation.

We first assayed the effects of 3’UTR elements on steady-state RNA stability by comparing total RNA abundance to DNA payload for each member of the reporter library (**Fig. S6F**). Consistent with previous results in other cell types,^45,46^ we found that steady-state stabilizing motifs in neurons tended to be more GC-rich, whereas destabilizing motifs tended to be more AU-rich (**Fig. S6G, H, I**). We observed a significant positive trend between endogenous transcript half-life and the average stability score of its constituent motifs in the MPRA, consistent with these elements exerting additive effects on RNA stability in neurons (**Fig. S6J**). We next assayed the effects of 3’UTR elements on activity-dependent RNA stability by comparing normalized total RNA abundance between KA- and PBS-treated animals, which revealed 303 activity-stabilizing and 283 activity-destabilizing elements (**Fig 3B, S6K, L**). For example, a 3’UTR motif within the activity-stabilized gene *Ildr2*, which encodes a transmembrane protein localized to the endoplasmic reticulum and cell junctions, is sufficient to drive activity-dependent reporter mRNA stabilization (**Fig. 3C**). In contrast, a 3’UTR motif within the activity-destabilized gene *Usp1*, which encodes a deubiquitinating enzyme involved in protein degradation, is sufficient to drive activity-dependent reporter mRNA destabilization (**Fig. 3D**).

In contrast to steady-state measurements, activity-stabilizing elements tended to be more AU-rich than unregulated or activity-destabilizing elements (**Fig. 3E**), with activity-stabilizing motifs strongly enriched for 7 nt k-mers containing AU-rich sequences (**Fig. 3F, S6M**). This finding reveals a surprising regulatory shift: while AU-rich sequences typically destabilize mRNAs under basal conditions,^46,47^ these same elements can become stabilizing motifs following neuronal activity. Conversely, activity-destabilizing elements showed enrichment for the DRACH motif (D=A, G or U; H=A, C or U), the canonical site of m^6^A RNA modification,^48,49^ suggesting that distinct regulatory factors mediate activity-dependent stabilization and destabilization. Overall, these results demonstrate that individual, modular 3’UTR elements are sufficient to drive bidirectional activity-dependent changes in RNA stability *in vivo* and suggest that multiple elements work cooperatively to regulate the stability of mRNAs.

### Profiling trans-acting factors that mediate activity-dependent RNA stability

To investigate whether trans-acting factors regulate endogenous transcripts that harbor activity-sensitive 3’UTR elements, we examined candidate RBPs with known or predicted affinity for k-mers enriched in our MPRA. We focused on two cytoplasmic Hu family proteins, HuD and HuR, that have been previously shown to bind sequences similar to our activity-stabilizing k-mers.^50^ We also included CELF4, a member of the CELF family of RBPs that binds UGUGUG motifs^51^ and the most highly expressed CELF family member in our neuronal cultures. Finally, we selected FUS, which does not have a strict consensus RNA motif but shows broad RNA-binding activity in a length- and structure-dependent manner and is implicated in neurodegenerative diseases such as frontotemporal dementia (FTD) and amyotrophic lateral sclerosis (ALS).^52^

We identified the RNA targets of CELF4, FUS, HuD and HuR by RNA immunoprecipitation followed by high-throughput sequencing (RIP-seq) from primary neuronal cultures that had been membrane depolarized for 1, 2, or 6 h, as well as an unstimulated control (**Fig. 4A, S7A-C, Table 4**). Most RNA targets were shared between HuD, HuR, and CELF4, consistent with previous findings that these RBPs bind similar RNA sequence motifs.^45,50^ In contrast, >50% of FUS RNA targets were unique to FUS (**Fig. 4B**). We next examined how RBP binding dynamics change in response to neuronal activity by comparing RIP-seq enrichment to total RNA abundance at various times after membrane depolarization. For each RBP, we grouped RBP-bound mRNAs into quartiles based on their degree of activity-dependent stabilization and compared the fold change in RBP binding between the most stabilized (Q1) and most destabilized (Q4) quartiles. FUS exhibited a transient but pronounced increase in binding to destabilized RNAs at 1 h post-stimulation that resolved by 6 h, suggesting a role in early destabilization events (**Fig. 4C, S7E**). In contrast, CELF4, HuR, and HuD showed preferential binding to activity-stabilized targets. CELF4 binding largely mirrored overall RNA abundance but displayed significantly greater enrichment on stabilized targets at 6 h post-stimulation (**Fig. 4C, S7D**). Most notably, HuR and HuD showed robust and sustained increases in binding at both 1 and 6 h, implicating these RBPs in activity-dependent RNA stabilization (**Fig. 4C, S7F, G**).

**Figure 4:**
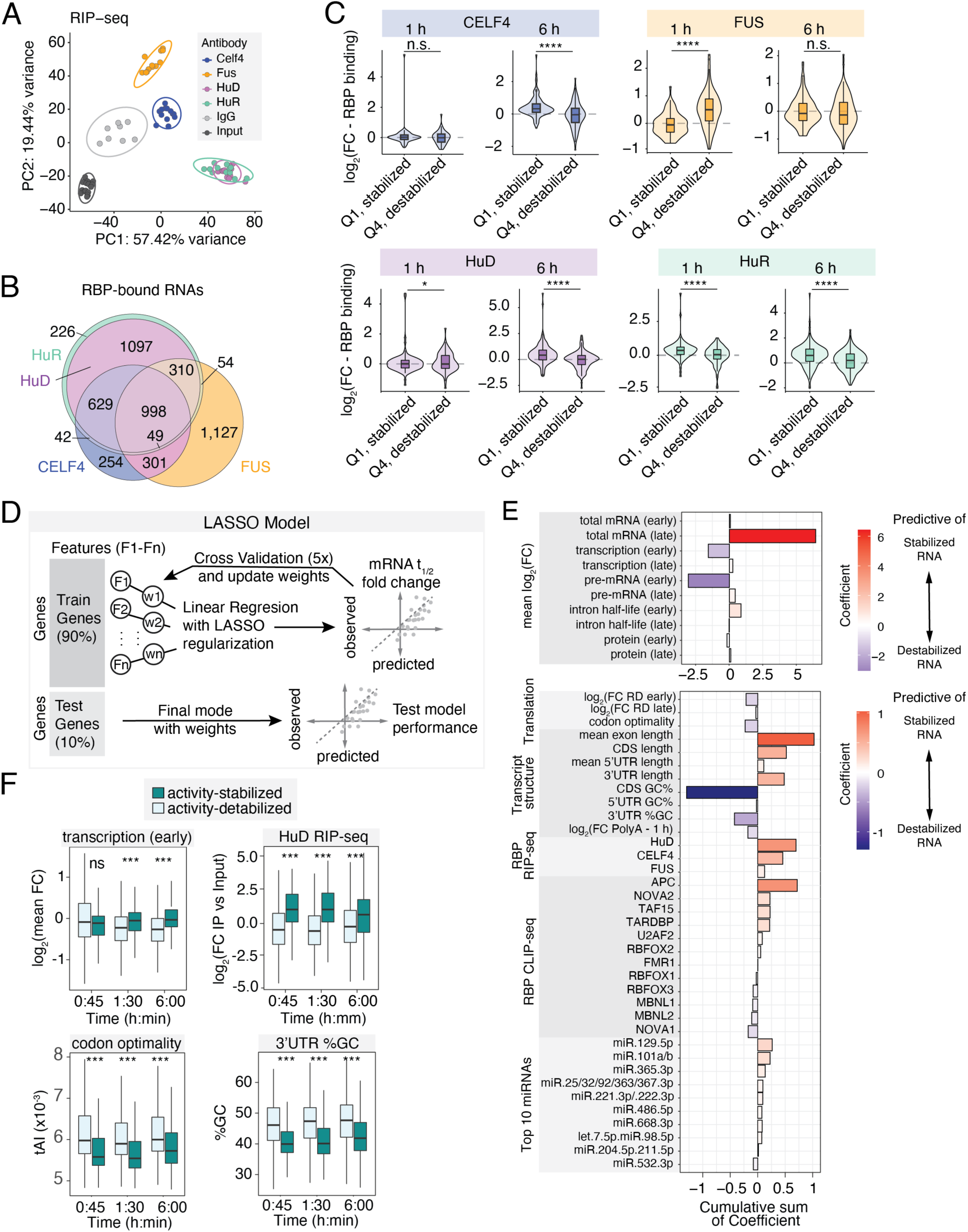
Profiling trans-acting factors that mediate activity-dependent RNA stability. (A) PCA plot of the top 2000 most variable genes in RIP-seq for HuD, HuR, CELF4 and FUS versus Input and IgG control. *n*=3-5 independent biological replicates per RBP and time point. (B) Venn diagram of overlap between RNA targets of each RBP defined by RIP-seq as >2-fold enriched over Input, excluding genes that were also enriched in IgG compared to Input. (C) Violin plots of log_2_(FC) in RBP binding at 1 and 6 h KCl, separated by RBP targets that are the most activity-stabilized (Q1) versus most activity-destabilized (Q4) at the matched time point. * = P < 0.05, **** = P < 0.0001 by KS test. (D) Schematic of machine learning model. Modeling was repeated for each time point of membrane depolarization (15 min - 8 h). (E) Bar plot of the cumulative sum of LASSO predictive Coefficients across all time points for features that have significant predictive power features determined by machine learning. (F) Boxplots showing representative features associated with cytoplasmic RNA stability, separated by activity-stabilized and activity-destabilized transcripts at three representative time points (0:45, 1:30, 6:00). Features include early transcriptional response (log₂ fold change < 2 h), HuD binding (RIP-seq), codon optimality (tRNA adaptation index, tAI), and 3′UTR %GC. Boxplots are shown as median ± IQR (whiskers = 1.5*IQR). Statistical significance was assessed with the Kruskal-Wallis test with BH correction; P_adj_ < 0.001 (***).

To determine whether changes in RBP binding were predictive of RNA stability changes and to identify the molecular determinants of activity-dependent RNA stabilization, we performed a systematic feature analysis of the stabilized RNAs using machine learning. We trained a LASSO regression model to predict the log_2_ fold change in RNA half-life at each time following neuronal activity (15 min to 8 h). We tested an extensive set of RNA features encompassing both cis-regulatory elements (sequence composition, UTR lengths, predicted secondary structures) and trans-regulatory factors, including published binding data for additional RBPs as well as our own RIP-seq data, and predicted miRNA binding sites for all miRNAs expressed in neurons. We also incorporated dynamic transcription rates and intron half-lives from metabolic labeling, along with ribosome density from ribosome profiling and protein abundance from TMT-based quantitative proteomic analysis (**Fig. 4D**). Because LASSO modeling can inconsistently underweight features that are highly correlated with others, we first assessed correlation between features and then retained a single representative feature from each correlated group to predict cytoplasmic RNA stability changes in response to membrane depolarization (**Fig. S8A, Table 5 and Methods**).

Our classifier achieved strong predictive performance with an average coefficient of determination (R^2^) > 0.3 (**Fig. S8B**) and average root mean squared error (RMSE) < 0.8 for all time points (**Fig. S8C**). Among all features, RNA-binding proteins (RBPs) such as HuD, CELF4, and APC emerged as strong predictors in the LASSO model, but HuD also showed the most pronounced enrichment for activity-stabilized transcripts genome-wide (**Fig. 4E, F, S8D**).

Additional predictive features reflected diverse regulatory signatures. Higher GC content in both coding sequences (CDS) and 3′ UTRs was more associated with activity-destabilized genes, while increased ribosome density and codon optimality were only modestly predictive (**Fig. 4E, F, S8D**). Mean exon length also emerged as predictive of activity-stabilized transcripts, although the underlying mechanism remains unclear. Notably, an increase in total RNA abundance at later time points (>2 h) was predictive of stabilization, but this is expected because stabilized RNAs accumulate over time and thus provide limited mechanistic insight (**Fig. 4E, S8D**). miRNA binding sites were generally less predictive than RBP binding, although miR-129-5p, which is known to contribute to synaptic plasticity,^53^ showed the strongest predictive power (**Fig. 4E, S8D**). In contrast, HuD binding stood out as both highly predictive and strongly enriched among stabilized RNAs, highlighting its central role in activity-dependent RNA stabilization. Together, these findings illustrate that activity-dependent RNA stability is determined by a complex interplay between sequence-intrinsic properties and protein binding, with HuD playing a prominent role in neuronal activity-dependent RNA stabilization.

### HuD regulates activity-dependent RNA stability

HuD is a neuronal-enriched RBP that has been implicated in diverse aspects of RNA metabolism, including mRNA stability, alternative splicing, and polyadenylation site selection, and is best studied for its role in promoting neurite outgrowth.^54^ However, despite decades of study, its primary function remains unclear. In particular, whether HuD plays a central role in shaping RNA stability at steady-state and in response to external stimuli, and whether these effects are similar across the transcriptome, remains unknown. To directly test whether HuD is required globally for activity-dependent RNA stabilization, we depleted HuD in primary neurons using shRNA (>80% knockdown of *Elavl4* mRNA and protein; **Fig. 5A, B, S9A**) and measured total RNA abundance and stability with s^4^U metabolic labeling (**Fig. 5A, S9B-F, Table 6**).^55,56^

**Figure 5:**
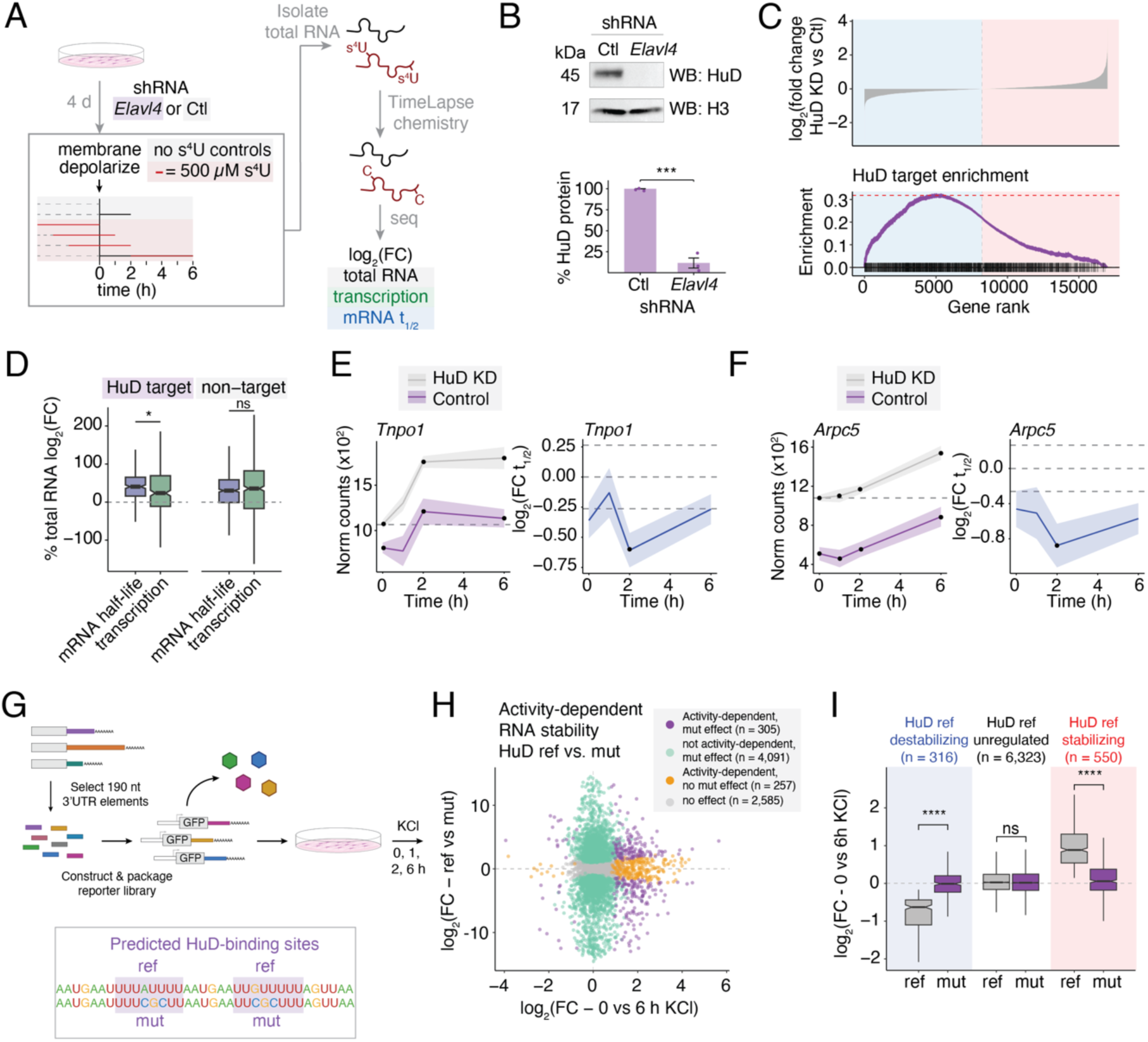
HuD regulates activity-dependent RNA stability]. (A) Schematic of experimental setup to knock down HuD in primary neuronal cultures using shRNAs against *Elavl4,* the gene that encodes HuD, versus a non-targeting shRNA control, followed by membrane depolarization and metabolic labeling to calculate log_2_(FC) in total RNA abundance, transcription, and mRNA half-life in HuD knockdown (KD) versus control. *n*=3 independent biological replicates per condition. (B) Representative western blot of HuD protein following shRNA KD in primary neuronal cultures versus a non-targeting control. Experiment was repeated three times with similar results. Unprocessed blots are provided in the source data. (C) Top: log_2_(FC) in mRNA abundance between HuD KD and control neurons, ordered from most downregulated (blue) to most upregulated (red) upon loss of HuD. Bottom: gene set enrichment analysis (GSEA) of HuD-bound mRNAs, where each black tick mark represents an HuD-bound mRNA and purple indicates the random walk enrichment score. Enrichment = 0.3319505, P < 1e-4, 10,000 permutations. (D) Box and whisker plot of the relative contribution of transcription and mRNA half-life to total RNA fold changes in HuD KD versus control at steady-state. Boxplots are shown as median ± IQR (whiskers = 1.5*IQR). P_adj_ = 0.0349 (*); P_adj_ = 0.258 (ns); KS test with BH multiple hypothesis correction. (E&F) Line plot of normalized RNA counts in HuD KD versus control (left) and log_2_(FC mRNA t_1/2_) between HuD KD and control (right) for *Tnpo1* (E) and *Arpc5* (F), two activity-stabilized mRNAs. Line = mean, shading = ± SEM. Black points indicate time points with a significant difference in expression between HuD KD and control (left, DESeq2 P_adj_ < 0.05; right, EZbakR P_adj_ < 0.1). (G) Schematic of mutational MPRA reporter library containing 3’UTR elements from HuD-bound mRNAs with two neighboring predicted HuD-binding sites (ref) versus mutation of both HuD-binding sites to CGC (mut) to disrupt HuD binding sites in primary cultured neurons. *n*=6 independent biological replicates. (H) Volcano plot of log_2_(FC) in RNA expression between ref and mut sequence (y-axis) and between 0 and 6 h KCl (x-axis). (I) Box plot of fold change in reporter RNA expression (0 vs 6 h KCl) between ref and mut sequences, separated by whether the ref sequence promotes activity-dependent reporter stabilization, destabilization, or has no effect on stability (unregulated). **** P_adj_ < 1e-16, ns P_adj_ < 0.05, KS test with BH correction.

While 5,222 genes were misregulated upon HuD knockdown, likely reflecting both direct and indirect effects of HuD loss, HuD-bound mRNAs were significantly more likely to be sensitive to HuD knockdown than mRNAs that do not bind HuD (Chi-squared test, p=9.37e-06). Furthermore, they were significantly more likely to be downregulated upon HuD knockdown (GSEA, P<1e-4, 10,000 permutations; **Fig. 5C, S10D**), consistent with the results of our machine learning predictions that HuD functions primarily as an mRNA-stabilizing factor. Thus, we focused on these HuD-sensitive RNAs, which we defined as HuD-bound mRNAs whose expression was perturbed upon HuD depletion (1,068 transcripts, **Fig. S10A**). Notably, HuD-sensitive RNAs contained significantly more predicted HuD-binding sites than unaffected mRNAs whose expression was unaffected by HuD disruption, suggesting that genes with more HuD-binding sites are more susceptible to perturbation upon HuD depletion (**Fig. S10B, C**). Many of the HuD-sensitive mRNAs are activity-regulated, and their expression trajectories were perturbed upon HuD knockdown (**Fig. S10A**). For example, the nuclear import factor *Tnpo1* is an activity-stabilized transcript that fails to be fully induced in HuD knockdown neurons following membrane depolarization due to reduced transcript stability (**Fig. 5E, F**). By contrast, *Arpc5*, a component of the actin-related protein 2/3 complex, is destabilized both at steady state and in response to activity in HuD-depleted neurons, highlighting that HuD loss can impact both basal and activity-dependent stability regulation (**Fig. 5E, F**).

Across all genes, changes in RNA abundance upon HuD depletion reflected contributions from both transcription and RNA stability, consistent with compensatory feedback loops. Importantly, for HuD-bound RNAs, changes in total RNA abundance were primarily driven by RNA stability, with less substantial contributions from transcription changes whereas for non-targets, changes in abundance were due equally to alterations in transcription and stability (**Fig. 5D**). We then stimulated the neurons and found that HuD depleted neurons did not mount the expected transcriptional response (**Fig. S10E, F**), consistent with changes in intrinsic excitability caused by prolonged HuD depletion, which has been previously suggested.^57^ Together, these results indicate that HuD-sensitive RNAs are uniquely dependent on RNA stability for both basal and activity-dependent regulation, highlighting HuD’s central role in shaping stimulus-responsive gene expression.

While global HuD knockdown can likely alter RNA stability through both direct and indirect mechanisms, we sought to determine which effects required direct HuD binding. To this end, we mutated HuD-binding sites within the 3’UTRs of 806 mRNAs using MPRAs and assessed the effect on reporter gene expression (**Fig. 5G**, **Table 7**). We scanned the 3’UTRs of mRNAs identified as HuD targets by our RIP-seq data for predicted HuD-binding sites, focusing on regions containing two predicted HuD-binding sites within 10 bp of one another, a configuration that has previously been shown to confer high-affinity HuD binding.^58,59^ We included 7,500 predicted HuD-binding sites in our MPRA library, along with 3 bp substitutions in each predicted HuD-binding site that were designed to disrupt HuD binding. This library was packaged into AAV, used to infect primary cultured neurons, and assayed for reporter RNA stability at baseline and following membrane depolarization by comparing RNA abundance to DNA payload. We found that 55% of the tested reference 3′UTRs significantly altered reporter RNA stability (**Fig. 5H**). Notably, HuD binding sites were sufficient to drive activity-dependent changes in RNA stability, with 550 motifs having an activity-stabilizing effect and 316 motifs having an activity-destabilizing effect. Mutating these HuD sites significantly blunted activity-dependent reporter RNA stability (**Fig. 5H, I**), confirming that sequence-specific HuD binding is required for the observed effects. Together, these comprehensive loss- and gain-of-function experiments establish HuD as a multifaceted regulator of both constitutive and activity-dependent mRNA stability in neurons, thereby expanding its previously recognized roles in neuronal gene regulation.

### HuD mediates local activity-dependent gene expression in soma and neurites

Neurites, the dendrites and axons that extend from the neuronal cell bodies (soma), are sites of RNA localization and translation critical for synaptic function. Given that these processes can span hundreds of microns and form synaptic networks far from the nucleus, neurons must regulate RNA stability and translation locally to respond rapidly to activity-dependent cues. As HuD protein is localized to soma and neurites in both primary neurons (**Fig. 6A**) and *in vivo,*^60^ we hypothesized that HuD regulates RNA stability in specific neuronal compartments. Local regulation of RNA stability in dendrites, axons, and at synapses would allow neurons to fine-tune the persistence of specific transcripts at sites of activity, thereby complementing local translation and transport mechanisms. Such spatially restricted stabilization would enable sustained protein production at active synapses without the need for new transcription, providing a rapid and localized means of reinforcing synaptic changes underlying plasticity.

**Figure 6:**
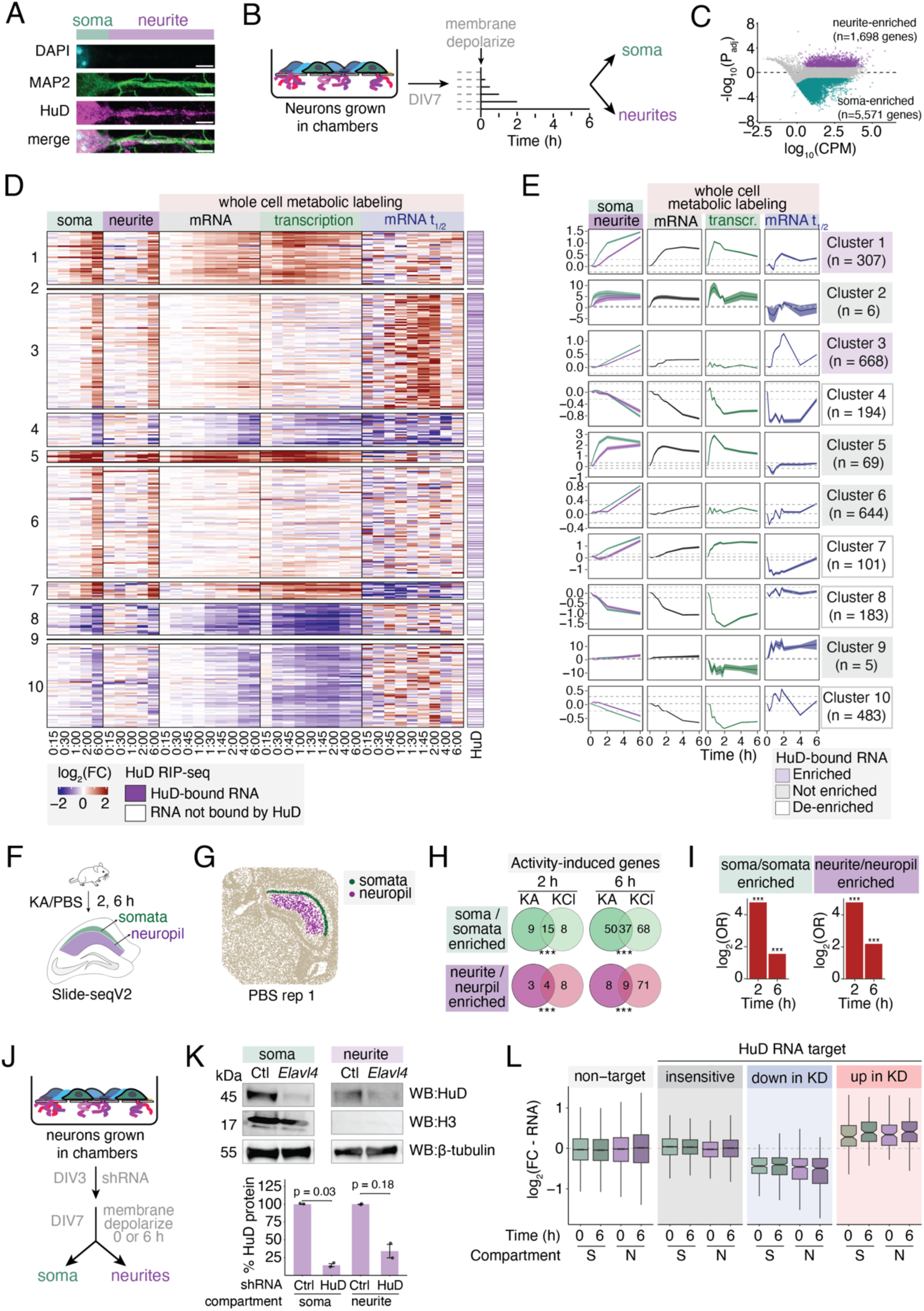
HuD mediates local activity-dependent gene expression in soma and neurites. (A) Representative maximum intensity project image of soma (green) and straightened neurites (purple) of primary neurons. Blue = DAPI, purple = MAP2, Cyan = HuD. Scale bar, 10 μm. (B) Schematic of RNA-seq strategy from membrane-depolarized primary neurons cultured in separation chambers. (C) MA plot of log_10_(counts per million reads (CPM)) versus log_2_(FC) between soma and neurite transcriptomes. Genes were defined as enriched in soma vs neurite compartments based on DESeq2 P_adj_ < 0.05 and log_2_(FC) > 1 (neurite-enriched) or < - 1 (soma-enriched). (D) Heatmap of log_2_(FC) in gene expression in soma and neurites following membrane depolarization, as well as log_2_(FC) in total mRNA, transcription, and mRNA t_1/2_ calculated in Fig. 1 for matched genes. Left annotation = HuD-bound mRNA as determined by RIP-seq. (E) Metaplots showing activity-dependent changes in mRNA abundance in soma and neurites, mRNA abundance in whole cells in primary neurons, as well as transcription rates and mRNA half-life as measured by metabolic labeling for clusters defined in (D). Dark line = mean log_2_(FC) of all genes in the cluster, shading = ±SEM, dashed lines = mean in PBS-injected animals (top) or unstimulated neurons (middle, bottom) ± 1.2-FC. Cluster headers are colored based on whether transcripts within a given cluster are enriched (purple), not enriched (gray), or de-enriched (white) for HuD-bound RNAs (Fisher’s Exact Test, P_adj_<0.05). (F) Schematic of Slide-seqV2 in the mouse hippocampus. *n*=2-3 mice per time point, including male and female animals. (G) Spatial plot of beads defined as CA1 soma (green) and neuropil (purple). (H) Venn diagram of overlap between activity-regulated genes in soma and neurites between KCl-depolarized neurons in separation chambers and KA-stimulated CA1 hippocampus. (I) Log-odds ratio test of overlap between the two datasets in (H). P_adj_ < 0.001 (***), Chi-squared overlap test. (J) Schematic of shRNA-mediated HuD knockdown in separation chambers. *n*=3 independent biological replicates. (K) Representative western blot (top) and quantification (bottom) of HuD protein following shRNA knockdown of *Elavl4* versus a non-targeting control in soma and neurites. Loading controls = H3 (soma only) and β-tubulin (soma and neurites). P-values determined by two-tailed t-test. (L) Boxplot of log_2_(FC) in RNA abundance in soma (S) vs neurites (N) for HuD-bound mRNAs and non-targets at 0 and 6 h post-KCl, separated by HuD-insensitive, upregulated and downregulated genes as defined in Fig. S10A.

While local translation of thousands of mRNAs occurs in dendrites and axons,^12^ whether the modulation of RNA stability similarly occurs in these compartments has not been previously addressed.

To investigate this possibility, we cultured primary neurons and then used a well-established protocol to separate soma and neuronal projections (neurites)^61–63^ followed by RNA-seq to assess mRNA expression at various times after membrane depolarization (**Fig. 6B**, **Table 8**). We confirmed that soma and neurite fractions were well-separated (**Fig. 6C, S11A, B**) and that the neurite fraction was enriched for transcripts known to encode proteins that localize to the pre- and post-synaptic specialization, suggesting that our fractionation protocol is able to capture both dendrites and axons as well as synapses (**Fig. S11C**).

We next clustered transcripts based on activity-dependent gene expression changes in soma and neurite fractions, together with whole-cell transcription rates and mRNA half-lives determined by RNA metabolic labeling (**Fig. 6D, E**). These analyses revealed diverse kinetics of gene expression changes across compartments. In some clusters, changes in RNA stability in the soma preceded changes in neurites and were accompanied by activity-dependent transcriptional responses. In contrast, clusters showing temporally coordinated changes in both soma and neurites tended to be enriched for activity-stabilized or -destabilized RNAs, suggesting that regulation of RNA stability occurs in both compartments with similar kinetics (**Fig. 6E**). Strikingly, two clusters enriched for activity-stabilized RNAs were also enriched for HuD-bound transcripts, which were coordinately upregulated in both soma and neurites (Clusters 1 and 3, **Fig. 6D, E, S11D**). Transcripts within these clusters encode both transcription factors and proteins that support local synaptic plasticity (**Fig. S11E**). These results suggest that HuD preferentially associates with activity-stabilized RNAs that are upregulated across subcellular compartments in neurons.

To determine whether these compartment-specific gene expression programs identified in cultured neurons also occur *in vivo*, we applied Slide-seqV2^64^ to mouse hippocampus after seizure induction with kainic acid (KA) or PBS control (**Fig. 6F, G, S12, Table 8**). KA induced robust transcriptional programs in hippocampal subregions, consistent with prior reports.^30,65,66^ Differential expression analysis revealed hundreds of transcripts enriched in somatic or neuropil layers of CA1, and these significantly overlapped with compartment-specific transcripts identified in separation chambers (FDR < 1e-4; **Fig. S13A, B**). Importantly, we also observed significant overlaps between activity-regulated genes in the two systems overall and between soma- and neurite-specific activity-regulated genes (FDR < 0.001; **Fig. 6H, I, S13C, D**). Thus, both *in vitro* and *in vivo* datasets capture conserved compartment-specific and activity-dependent gene expression programs.

Building on these findings, we tested the hypothesis that HuD can regulate RNA stability locally within neurites. We performed HuD knockdown in primary neurons and assessed mRNA levels in both soma and neurites following membrane depolarization (**Fig. 6J, K**, **Table 8**). We found that upon HuD knockdown, the expression of HuD-sensitive mRNAs decreases in both the soma and the neurites following membrane depolarization, while the level of mRNAs that are upregulated upon HuD loss increased in both soma and neurites (**Fig. 6L**). While our half-life measurements were performed in whole cells, the consistent directionality and magnitude of expression changes across soma and neurites following HuD depletion strongly support a model in which HuD regulates –––mRNA stability locally within neuronal processes that form synapses.

### Neuronal activity reorganizes the HuD protein interactome

We next sought to determine the mechanisms by which HuD function is modulated by neuronal activity. As HuD protein levels remained unchanged following membrane depolarization (**Fig. S14A, B**), and no significant activity-dependent changes at detectable HuD phosphorylation sites were observed (**Fig. S14C-F**), we hypothesized that neuronal activity alters HuD function by reshaping its protein-protein interactions. To test this idea, we performed RNase-treated immunoprecipitation followed by quantitative mass spectrometry (IP-TMT-MS) in primary neuronal cultures expressing FLAG-HA-tagged HuD (HuD-FH) or a GFP control (**Fig. 7A**). We identified 40 “core interactors” that were significantly enriched for binding to HuD versus GFP at all time points (0, 1, 6 h), along with 20 proteins that preferentially bound to HuD at baseline but not upon stimulation and 19 that interacted with HuD specifically following membrane depolarization, indicating a broad reorganization of the HuD interactome in response to membrane depolarization of neurons (**Fig. 7B, S14G, H, Table 9**).

**Figure 7:**
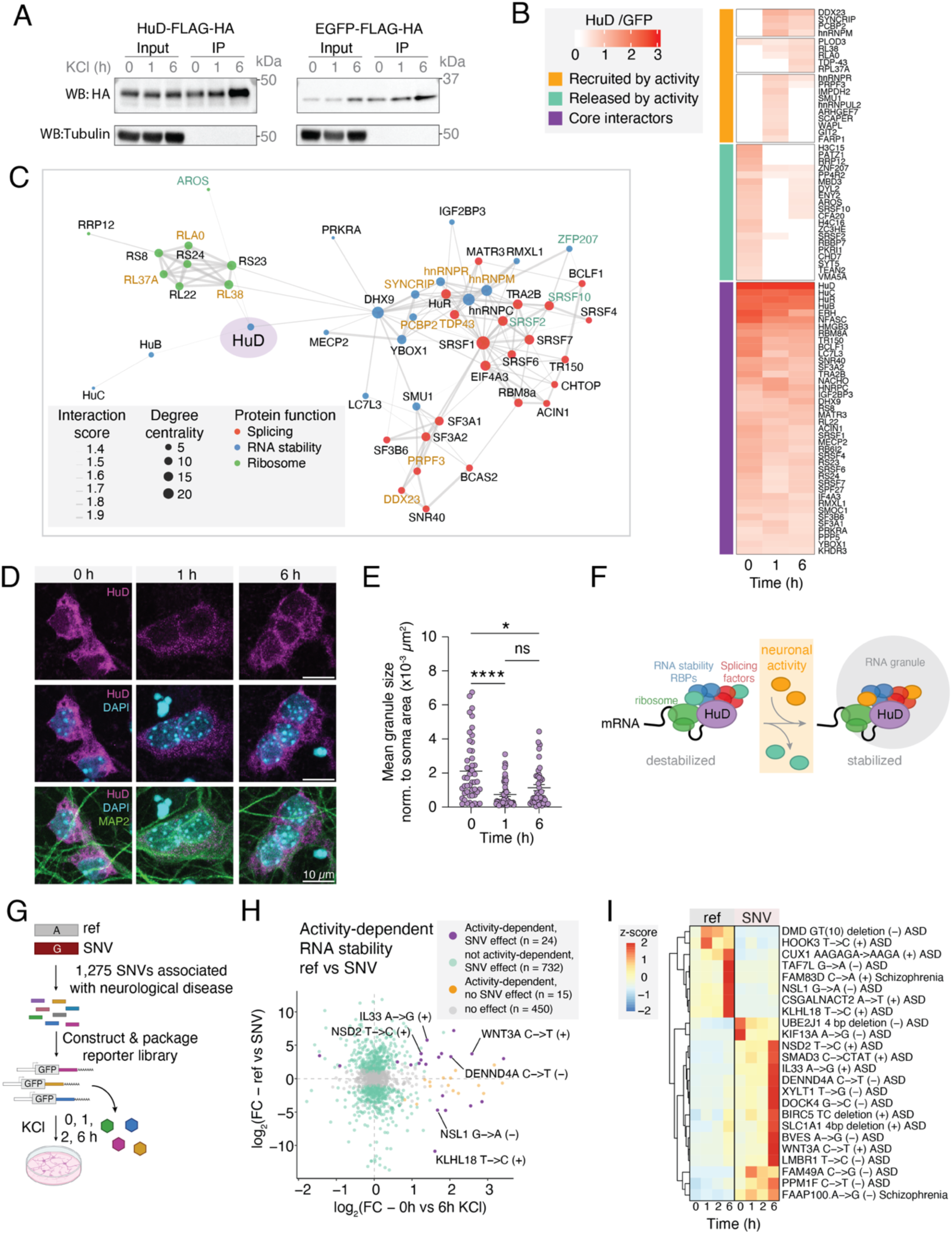
Neuronal activity reorganizes the HuD protein interactome and disease-associated variants disrupt activity-dependent RNA stability. (A) Representative western blot of HuD-FLAG-HA and EGFP-FLAG-HA immunoprecipitation from primary neurons depolarized with KCl. Experiment was repeated three times with similar results. Unprocessed blots are provided in the source data. (B) Heatmap of HuD interactors by IP-TMT-MS, separated into core interactors and activity-dependent interactors. (C) STRING map of proteins that interact with HuD, colored by main function. Names of proteins that are recruited to HuD following neuronal activity are colored in orange, while proteins that are released by activity are colored in green. (D) Representative image of HuD in unstimulated versus membrane-depolarized neurons. Blue = DAPI, green = MAP2, magenta = HuD. Scale bar = 10 µm. (E) Quantification of mean HuD granule size after normalization to soma size for each cell, nonparametric Kruskal-Wallis comparison with Dunn’s multiple comparison correction, P_adj_ <0.0001 (****), P_adj_ = 0.0213 (*). N = 42-49 cells, *n*=2 biological replicates. (F) Model for how HuD contributes to activity-dependent RNA stabilization. (G) Schematic of MPRA to assess the effects of neurological disorder-associated SNVs on RNA stability. *n*=6 independent biological replicates. (H) Scatterplot of log_2_(FC) in RNA abundance (normalized to DNA payload) in ref vs SNV, versus log_2_(FC) in RNA abundance with activity. Teal = SNVs that affect steady-state RNA stability, orange = activity-dependent RNA elements with no effect of SNV on stability, purple = SNVs that affect activity-dependent RNA stability, gray = no change in RNA stability with SNV or activity. (I) Heatmap of depth- and payload-normalized RNA counts (z-score) for ref and SNV motifs within motifs in which SNVs affect activity-dependent RNA stability.

HuD-associated proteins fall into three main classes: ribosomal proteins, RBPs that have previously been linked to mRNA stability (e.g. Hu proteins, IGF2BP3, PCBP2), and known splicing factors, many of which also influence RNA stability in the cytoplasm (e.g. SRSF proteins ^67^, MATR3 ^68^, and hnRNPC ^69^ (**Fig. 7C, S15A**). AlphaFold-multimer was used to predict structural interfaces between HuD and its co-immunoprecipitated partners,^70^ providing additional insight into how activity influences HuD-protein interactions (**Fig. S15B**). Across 30 predicted HuD-partner pairs, 10 displayed high-confidence interaction residues within 8 Å. Notably, two of these proteins, PCBP2 and MATR3, showed activity-dependent phosphorylation changes at their predicted interaction interfaces with HuD; PCBP2 was dephosphorylated and MATR3 phosphorylated upon membrane depolarization (**Fig. S15C–H**). Such activity-dependent modifications at HuD interfaces may alter its interaction landscape, potentially influencing how HuD is redistributed into RNA granules.

Notably, several activity-induced HuD interactors, including TDP-43, SYNCRIP, and PCBP2, are also components of neuronal RNA granules,^71–75^ suggesting an activity-dependent redistribution of HuD to RNA granules. Consistent with this idea, immunostaining revealed that membrane depolarization acutely shifts HuD from a diffuse cytoplasmic pattern to smaller, punctate granules within 1 h of activity (**Fig. 7D, E, S15I, J**), and these granules were also observed in neurites (**Fig. 6A**). Although RNA granules are often associated with translational repression, many neuronal granules support ongoing or regulated translation.^76,77^ To determine whether HuD-bound RNAs undergo translational changes following activity, we analyzed ribosome occupancy using ribosome profiling and RNA-seq. HuD-bound mRNAs exhibited a small but significant reduction in ribosome density compared to RNAs not bound by HuD following stimulation (**Fig. S15K**). This modest decrease argues against widespread ribosome stalling or translational silencing, and instead suggests that HuD-bound RNAs remain largely engaged with ribosomes even after redistribution into granules. Together, these findings support a model in which neuronal activity modulates HuD-containing RNPs into compact RNA granules that stabilize mRNAs while maintaining their translational competence.

### Disease-associated variants disrupt activity-dependent RNA stability

Having established HuD as a key regulator of activity-dependent mRNA stability, we next asked whether the disruption of this post-transcriptional control contributes to neurological disease. Notably, HuD-bound mRNAs are significantly enriched for ASD-associated genes, with 325 of 1,238 SFARI-annotated genes bound by HuD (Fisher’s exact test, *P* < 2.2e-16), suggesting that impaired regulation of RNA stability may contribute to pathogenesis. Moreover, genetic variants within 3′UTR sequences that regulate activity-dependent RNA stability represent a previously unexplored mechanism by which noncoding variation may alter neuronal gene expression. We therefore investigated whether genetic variants associated with neurological disorders that lie within 3’UTRs alter mRNA stability.

To this end, we examined the effect of 1,275 single-nucleotide variants (SNVs) and insertions/deletions (indels) previously associated with neurological disorders (ASD, schizophrenia, ADHD, MDD, ID) by GWAS^77,78^ on mRNA stability using a MPRA in primary mouse neurons (**Fig. 7G**, **Table 10**). Each human reference or variant 3′UTR sequence (with flanking context) was cloned downstream of GFP driven by the constitutive EF1α promoter, and the MPRA library was packaged into AAV for neuronal delivery. Following infection, neurons were stimulated by membrane depolarization (1, 2, and 6 h), and reporter RNA abundance was quantified by high-throughput sequencing to assess RNA stability at baseline and after stimulation. We identified 732 variants (57%) that altered steady-state RNA stability and 24 that significantly affected activity-dependent stability (**Fig. 7H**). For example, *Klhl18* mRNA is bound by HuD and stabilized in response to neuronal activity, and an ASD-associated T-to-C SNV in the human *KLHL18* 3′UTR disrupts a predicted HuD site and abolishes activity-dependent reporter RNA stabilization (**Fig. 7I**). Conversely, *Nsd2* mRNA is not activity-regulated in mouse neurons, but a T-to-C SNV in the human *NSD2* 3′UTR disrupts a predicted HuD site and facilitates activity-dependent reporter RNA stabilization (**Fig. 7I**). These findings demonstrate that variants associated with neurological disorders can rewire RNA stability landscapes under both basal conditions and in response to neuronal activity, by modifying sequence features that are critical for mRNA stability.

Together, these results reveal that 3′UTRs act as dynamic regulatory hubs integrating neuronal activity, RNA-binding proteins, and sequence variation to fine-tune RNA lifespan. Because dendritic and axonal mRNAs often rely on local stability to sustain protein synthesis at synapses, even subtle perturbations to these mechanisms may have lasting effects on circuit development and plasticity. The enrichment of HuD-bound and stability-regulated RNAs among ASD-associated genes suggests that disrupted 3′UTR-mediated RNA control could represent a convergent molecular vulnerability in neurodevelopmental disorders. By linking disease-associated variants to activity-dependent defects in RNA stability, our findings position the 3′UTR as a critical interface between neuronal signaling and genetic risk for brain disorders.

## DISCUSSION

Gene expression is regulated through coordinated changes between RNA synthesis, processing, degradation, and translation. While individual steps in the RNA life cycle have been studied in isolation, how these processes are integrated to control gene expression in response to external stimuli remains a fundamental open question. In neurons, activity-regulated gene programs have traditionally been studied in the context of transcription and local translation, where these processes have been shown to be important to facilitate synaptic plasticity, learning, and memory. While a handful of genes have been shown to undergo neuronal activity-dependent changes in RNA stability (e.g. *Adcyap1,*^79^ *Fos* and *Bdnf*^80^) the extent to which acute modulation of cytoplasmic RNA turnover globally contributes to activity-dependent gene responses is not well understood. Our findings establish that activity-dependent modulation of RNA stability is a widespread mechanism that can amplify transcriptional responses, buffer against excessive RNA accumulation, or operate independently of transcription to shape gene expression dynamics locally within neuronal processes.

Using MPRAs, we demonstrate that discrete sequence elements within the 3’UTRs are sufficient to drive bidirectional, activity-dependent changes in RNA stability *in vivo*. Unexpectedly, we find that AU-rich sequences, classically known to destabilize mRNAs under basal conditions, were found to act as stabilizing elements following neuronal depolarization. This switch underscores the context-dependent nature of RNA stability regulation and shows that single sequence motifs can exert opposing effects depending on cell state. These findings are particularly notable given that many stability-regulated mRNAs are enriched in neurites, where their dynamics are consistent with local activity-dependent post-transcriptional control. Because neurons experience spatially restricted inputs rather than uniform stimulation across the plasma membrane, local RNA stabilization may allow for compartment-specific increases in RNA and protein abundance without eliciting a global response, thereby contributing to localized synaptic plasticity.

We further identify HuD as a central regulator of both basal and activity-dependent RNA stability across a broad set of transcripts. While previous work suggested that HuD abundance increases 24–72 h after neuronal stimulation,^60,81^ we observed no significant change in HuD protein levels within 6 h of depolarization. This indicates that acute HuD-mediated changes in RNA stability are more likely governed by alterations in protein–protein interactions than by HuD abundance. Supporting this view, neuronal activity triggered a dynamic remodeling of HuD ribonucleoprotein complexes, including recruitment of the RNA granule–associated proteins TDP-43, SYNCRIP, and PCBP2. We propose that activity-dependent recruitment of these proteins promotes relocalization of HuD-bound transcripts into granules that protect RNAs from degradation while preserving translational competence.

Several additional neuronal RBPs that regulate mRNA translation have been implicated in activity-dependent processes, including FMRP (Fragile X Mental Retardation Protein), which regulates dendritic mRNA translation and is critical for synaptic plasticity,^82^ and CPEB (Cytoplasmic Polyadenylation Element Binding protein), which controls activity-dependent polyadenylation and translation of plasticity-related mRNAs.^83^ Notably, neither FMRP nor CPEB was detected as a HuD interactor in our IP-MS experiments, suggesting that HuD-containing granules represent a distinct regulatory environment. Whereas FMRP- and CPEB-containing granules primarily modulate translation, HuD granules may preferentially stabilize mRNAs. Such division of labor could provide an efficient mechanism for transiently coupling stimulus-dependent mRNA stabilization and translation to mediate aspects of neural plasticity.

Consistent with this model, neural disruption of HuD function in the mouse has been shown to impair activity-dependent processes such as learning and memory.^57,84^ Moreover, aberrant HuD activity has been linked to neurodegenerative disorders, including frontotemporal dementia and Alzheimer’s disease.^85^ Together, these observations highlight the importance of HuD-mediated RNA stability control in neuronal physiology and suggest that further elucidation of the mechanisms mediating HuD-regulated mRNA stability may uncover new therapeutic targets for neurological disease.

Taken together, our findings establish RNA stability as a widespread mode of activity-dependent gene regulation in neurons, working synergistically with transcriptional mechanisms to provide precise temporal and spatial control of gene expression. The identification of disease-associated variants that disrupt RNA stability regulation has important implications for understanding neurodevelopmental disorders. Although the link between individual 3’UTR variants and neurological disease remains uncertain, the finding that over half of tested variants disrupt basal RNA stability, and two dozen alter activity-dependent stability, suggests that dysregulation of this mechanism may broadly contribute to neurodevelopmental disorders. These results point to RNA stability as a potentially underappreciated layer of gene regulation relevant to neurological and psychiatric disease. The identification of HuD as a regulator of this process, operating through activity-dependent recruitment into RNA granules, may also provide new insight into the molecular mechanisms that potentially underlie synaptic plasticity.

## Resource Availability

Data reprocessed from Traunmüller et al.^32^ were accessed from the Gene Expression Omnibus database under accession number GSE283483.

## Acknowledgements

The Greenberg Laboratory is supported by the Allen Discovery Center Program, a Paul G. Allen Frontiers Group advised program of the Paul G. Allen Family Foundation and the Tang-Yang Autism Center at Harvard Medical School. E.E.D. was a Damon Runyon-National Mah Jongg League, Inc. Breakthrough Scientist supported by the Damon Runyon Cancer Research Foundation and was supported by the Warren Alpert Distinguished Scholar Award and a Mahoney Postdoctoral Fellowship. I.P. was supported by a Parkinson Foundation Postdoctoral Fellowship. S.Kalaora was supported by the Helen Hay Whitney Foundation (Merck fellow) and a Gene Lay Institute Postdoctoral Fellowship. L.T. was supported by an EMBO Postdoctoral Fellowship, Long-term Human Frontiers Science Program Fellowship and the William Randolph Hearst Fund. S.Krüttner was supported by a Y. Eva Tan Fellowship of the Tan Yang Autism Center at Harvard University. S.A.K. was supported by the NSF GRFP DGE 2140743. M.Y.F. was supported by an Ontario Graduate Scholarship. L.S.C. was supported by funding from NIA R21 AG091645 and NHGRI R01 HG007173. M.E.G. was supported by funding from NINDS R01 NS115965. The funders had no role in study design, data collection and analysis, or the decision to publish, or preparation of the manuscript. We thank members of the Greenberg and Churchman laboratories for helpful discussions on the manuscript. We thank Stefano de Pretis and Mattia Furlan for helpful discussions about the INSPEcT pipeline. We thank Isaac Vock and Matthew Simon for helpful discussions about EZbakR implementation. We thank the Harvard Medical School Neurobiology Department for consultation and instrument availability that supported this work.

## Author Contributions

E.E.D. and I.L.P. performed all RNA-seq assays. S.Kalaora, I.L.P., J.A.P. and S.P.G. performed mass spectrometry based proteomics. E.E.D. and I.L.P. performed all bioinformatic analysis. I.L.P. designed and implemented machine learning modeling. L.T. and S.Krüttner performed stereotaxic injections and hippocampal microdissection. B.T.K. and M.Y.F. performed spatial transcriptomics. E.G.A. performed RNA immunoprecipitation. E.G.A. and M.M.M., performed western blotting and protein immunoprecipitation. N.P., B.F., and E.E.D. performed reporter assays. Z.B. prepared viruses. S.A.K. and E.G.A. performed immunofluorescence, and S.A.K. performed image analysis. E.E.D., I.L.P., S.Kalaora, J.A.P., and S.P.G. performed immunoprecipitation followed by proteomics. M.M.M. performed Alphafold predictions. E.E.D. and M.E.G. conceptualized the study. E.E.D., I.L.P., E.C.G., L.S.C. and M.E.G. drafted the manuscript, with input from all co-authors.

## SUPPLEMENTARY FIGURES

**Supplementary Figure 1:** RNA metabolic labeling quality metrics, Related to Figure 1.

**Supplementary Figure 2:** Ribo-seq and whole-cell proteomics quality metrics, Related to Figure 1.

**Supplementary Figure 3**: Measuring activity-dependent RNA dynamics, Related to Figure

**Supplementary Figure 4:** Activity-dependent modulation of RNA stability is widespread and impacts protein abundance, Related to Figure 1.

**Supplementary Figure 5:** Activity-dependent modulation of RNA stability is widespread and a key determinant of RNA and protein output, Related to Figure 2.

**Supplementary Figure 6:** Modular 3’UTR RNA sequences are sufficient to drive activity-dependent changes in RNA stability *in vivo,* Related to Figure 3.

**Supplementary Figure 7:** RIP-seq QC, Related to Figure 4.

**Supplementary Figure 8:** LASSO machine learning QC, Related to Figure 4.

**Supplementary Figure 9:** HuD regulates activity-dependent RNA stability, Related to Figure 5.

**Supplementary Figure 10:** HuD regulates activity-dependent RNA stability, Related to Figure 5.

**Supplementary Figure 11:** Separation chamber RNA-seq QC, Related to Figure 6.

**Supplementary Figure 12:** Slide-seqV2 QC, Related to Figure 6.

**Supplementary Figure 13:** Overlap between Slide-seqV2 and separation chamber RNA-seq, Related to Figure 6.

**Supplementary Figure 14:** Activity-dependent HuD protein-protein interactions, Related to Figure 7.

**Supplementary Figure 15:** Activity-dependent HuD protein-protein interactions, Related to Figure 7.

## METHODS

### EXPERIMENTAL MODEL

#### Mouse models

Animals were handled according to protocols approved by the Harvard University Standing Committee on Animal Care and were in accordance with federal guidelines. Wild-type C57/BL6 (JAX 000664) were housed in a 12 h light/dark cycle with ad libitum access to food. Temperatures range from 67 to 74°F and humidity between 35 and 65%. For experiments in which seizures were induced, P28-P32 C57BL/6 mice were injected intraperitoneally (IP) with kainic acid (KA, 15-20 mg/kg, Sigma Aldrich K0250) or PBS and sacrificed 2 or 6 h after injection. All experiments were sex-balanced, with both male and female mice included in each experimental condition.

#### Mouse neuron culture

Embryonic hippocampi and/or cortices from C57BL/6 mice (JAX 000664) were dissected at age E16.5 and then dissociated with papain (Sigma Aldrich 10108014001). Tissue was pooled from embryos in the same litter to generate mixed sex cultures. Papain digestion was quenched by the addition of ovomucoid trypsin inhibitor (Worthington LS003086), and cells were gently triturated through a P1000 pipet tip and passed through a 40 µm cell strainer. Neurons were quantified using trypan blue stain on a Countess II Automated Cell Counter (Thermo Fisher) and plated onto cell culture dishes pre-coated overnight with poly-D-lysine (20 µg/mL, Sigma Aldrich P7405) and laminin (4 µg/mL, Life Technologies 23017015). Neurons were cultured in Neurobasal medium (Life Technologies 21103049) containing B27 supplement (2%, Life Technologies 17504044), penicillin-streptomycin (50 U/mL each, Life Technologies 15140122), and glutaMAX (1 mM, Life Technologies 35050061). Neurons were incubated at 37°C with a CO--_2_ concentration of 5%. At 3-4 days *in vitro* (DIV), 50% of the culture media was exchanged for fresh media. To prevent spurious activity, neurons were silenced at 6DIV overnight by the addition of 1 µM TTX (Abcam ab120055) and 100 µM AP5 (Fisher Scientific 010610). Neurons were harvested at 7DIV. In all experiments, independent replicates were generated by preparing primary cultures from pools of embryos dissected from different dams.

### METHOD DETAILS

#### RNA pulse labeling with s^4^U

Hippocampal neurons were plated as described above in 6-well dishes at 1×10^6^ neurons/well and cultured for 7DIV. At 7DIV, silenced neurons were depolarized for 15, 30, 45, 60, 75, 90, 120, 240 (4 h), 360 (6 h), or 480 (8 h) by addition of depolarization buffer (Final concentrations in media: 55 mM KCl, 660 µM CaCl_2_, 330 mM MgCl_2_, 3.33 mM HEPES pH 7.4) or a mock depolarization buffer in which 55 mM KCl is replaced with equimolar NaCl to control for changes in osmolarity with the addition of buffer to media. During the last 15 min before cell harvest, s^4^U (1 mM final concentration, Sigma Aldrich T4509) was added directly to neurons in culture media and mixed gently. After 15 min RNA metabolic labeling with s^4^U, neurons were washed twice with 1X PBS to remove dead cells and scraped immediately into Trizol (Life Technologies 15596026).

The enrichment of s^4^U-RNA was performed using MTS-biotin chemistry as previously described.^22,87^ Cells lysed in Trizol were chloroform extracted once, and nucleic acids were precipitated using isopropanol. DNA was removed using Turbo DNase (Life Technologies AM1907), and RNA was purified using Agencourt RNAClean XP beads (Fisher Scientific NC0068576). One volume of RNAClean beads was added to each sample, incubated at room temperature for 8 min, and captured on a magnetic rack. Beads were washed twice with 80% ethanol, and RNA was eluted from dried beads using RNase-free water. Total RNA fractions were prepared by reserving 1% of input RNA before MTS-biotin enrichment. To biotinylate s^4^U-RNA, samples (15-20 µg total RNA) were incubated with 2 μg MTS-biotin in biotinylation buffer (10 mM HEPES pH 7.4, 1 mM EDTA, 20% dimethylformamide) for 30 min in the dark. Excess biotin was removed via chloroform extraction using Phase-Lock Gel Tubes. RNA was precipitated with a 1:10 volume of 3 M NaOAc and an equal volume of isopropanol and centrifuged at 20,000 × g for 20 min. The pellet was washed with an equal volume of 75% ethanol. Purified RNA was dissolved in 50 μL RNase-free water. Biotinylated RNA was separated from non-labeled RNA using glycogen-blocked Dynabeads Streptavidin C1 Beads (Invitrogen). Beads (10 μL) were added to each sample and incubated for 15 min at room temperature, then washed three times with high salt wash buffer (100 μL each, 100 mM Tris-HCl [pH 7.4], 10mM EDTA, 1 M NaCl, and 0.1% Tween-20). In order to improve the stringency of the washes, an additional 3 washes with buffer TE (10 mM Tris pH 7.4, 1 mM EDTA) at 55°C were added to the protocol. s^4^U-RNA was eluted from Dynabeads with 25 μL freshly prepared elution buffer (10 mM DTT, 100 mM NaCl, 10 mM Tris pH 7.4, 1 mM EDTA) and incubated for 15 min, followed by a second elution with an additional 25 μL elution buffer. Both elutions were pooled and purified using RNAClean beads as above.

Libraries were prepared from input and s^4^U-enriched RNA samples using the SMARTer Stranded Total RNA-seq Kit – Pico Input Mammalian V2 (Takara Bio 634413) according to the manufacturer’s instructions. Input and s^4^U-enriched samples were multiplexed with Illumina barcodes and sequenced using paired-end 2×75-nt cycles on an Illumina NextSeq 500 instrument to a depth of at least 20M reads per sample.

#### Ribosome profiling

Cortical neurons were plated as described above in 10 cm dishes at 1×10^7^ neurons/well and cultured for 7DIV. At 7DIV, silenced neurons were depolarized for 1, 2, or 6 h by addition of depolarization buffer or mock buffer. Neurons were washed once with 1X PBS supplemented with 100 µg/mL cycloheximide and scraped into 1xPBS supplemented with 100 µg/mL cycloheximide. Cell pellets were snap-frozen in liquid nitrogen, followed by storage at −80°C. Frozen cell pellets were thawed on ice. Each sample was dounced 15 times in 400 µL of ice-cold lysis buffer: 20 mM Tris pH 7.4, 150 mM NaCl, 5 mM MgCl2, 1 mM DTT and 100 µg/mL of cycloheximide (Sigma-Aldrich). The lysate was further sheared using a 26-gauge syringe. The lysate was clarified by centrifugation at 20,000 x g for min at 4 °C, then subjected to RNase I digestion (0.5 U of RNaseI per µg of RNA) at room temperature for 15 minutes with gentle agitation. To isolate ribosome-protected fragments, the RNase-digested lysate was transferred to Ultra-Clear 11 × 34 mm centrifuge tubes (Beckman Coulter) and underlaid with 0.9 mL of sucrose cushion. Samples were centrifuged in a TLS-55 rotor at 51,000 rpm for 2 h at 4 °C. The supernatant was discarded, and the pellet was resuspended in 300 µL of TRIzol. Ribosome-protected fragments were purified from TRIzol using the Zymo Direct-zol kit. RNA was precipitated by adding precipitation buffer (15 µg GlycoBlue (ThermoFisher), 100 mM sodium acetate pH 5.5, 50% isopropanol in nuclease-free water) to a final volume of 300 µL. The mixture was incubated overnight at −20°C. Samples were centrifuged for 30 min at 20,000*g* at 4 °C. The supernatant was discarded, and the RNA pellet was resuspended in 5 µL of 10 mM Tris pH 8. Next, 5 µL of 2x denaturing sample loading buffer (10 mM EDTA and 300 µg/mL bromophenol blue in formamide) was added to each sample, and samples were denatured at 80°C for 90 s. Ribosome-protected fragments, along with control oligos^88^ and 12 µL of NEB miRNA marker, were run on a 15% polyacrylamide gel at 200 V for 65 min. The gel was stained with SYBR Gold in 1x TBE, and gel fragments between 17 and 34 nt were excised and placed in a microfuge tube with 400 µL of gel extraction buffer (300 mM sodium acetate, 1 mM EDTA, 0.25% SDS). Samples were frozen on dry ice for 30 min and then thawed overnight with gentle agitation.

After overnight gel extraction, 400 µL of eluate was transferred to a new microfuge tube. RNA was precipitated by adding 1.5 µL GlycoBlue and 500 µL isopropanol. After overnight incubation at −20°C, the sample was centrifuged at 20,000*g* for 30 min at 4 °C. The supernatant was discarded, and precipitated RNA was resuspended in 4 µL of 10 mM Tris pH 8. Samples were then dephosphorylated using T4 PNK (4 µL RNA in 10 mM Tris pH 8, 0.5 µL T4 PNK enzyme, 0.5 µL T4 PNK buffer and 0.5 µL Superasin) at 37°C for 1 h. Samples were then purified with RNAclean SPRI beads: 50 µL of sample in RNase-free water was added to 90 µL RNAclean beads plus 270 µL isopropanol. After washing with 85% ethanol, dried beads were resuspended in 7 µL of nuclease-free water. The supernatant was collected, and sequencing libraries were prepared using the Clontech smRNA library prep kit (Takara) according to the manufacturer’s instructions. Libraries were sequenced on an Illumina NovaSeq S2 with single-end 1× 50-nt reads.

#### Direct RNA sequencing

Neurons were plated as described above in 6-well dishes at 1×10^6^ neurons/well and cultured for 7DIV. At 7DIV, silenced neurons were depolarized for 1 h by the addition of depolarization buffer as above, as well as a mock depolarization control. Cells were harvested in ice cold PBS and cell pellets were flash frozen and stored at −80 °C. RNA was extracted with Trizol following manufacturer instructions (Thermofisher, 15596026). RNA Poly(A)+ RNA was enriched using the Dynabeads mRNA purification kit (ThermoFisher, 61006) following manufacturer’s instructions. Direct RNA library preparation was performed using the SQK-RNA004 kits (Oxford Nanopore Technologies) with 500–700 ng of poly(A)+ RNA according to manufacturer’s instructions with the following exceptions: the RNA Calibration Strand (RCS) was omitted and replaced with 0.5 µL water, and the ligation of the reverse transcription adapter was performed for 15 min. Libraries were sequenced for up to 72 h with PromethION 2 Solo device (Oxford Nanopore Technologies).

#### Whole-cell proteomics and phosphoproteomics

Neurons were plated as described above in 6-well dishes at 1×10^6^ neurons/well and cultured for 7DIV. At 7DIV, silenced neurons were depolarized for 15, 30, 60, 120, or 360 (6 h) min by the addition of depolarization buffer as above, as well as a mock depolarization control. Cells were washed twice with ice cold PBS, harvested in ice cold PBS, and cell pellets were flash frozen and stored at −80 °C. Cell pellets were syringe-lysed in 8 M urea and 200 mM EPPS (pH 8.5) with protease and phosphatase inhibitors. BCA assay was performed to determine protein concentration of each sample. Samples were reduced in 5 mM TCEP, alkylated with 10 mM iodoacetamide, and quenched with 15 mM DTT. Proteins were precipitated using SP3 beads, and then digested on the beads at a 1:100 protease-to-peptide ratio with Lys-C overnight at room temperature, followed by Trypsin digestion for 6 h at 37°C. Following digestion, supernatant was collected to a different tube, and acetonitrile was added to the eluted peptides for a final volume of 30%. Peptides were labeled with the tandem mass tag (TMT) pro18-plex reagent for 1 h. The reaction was quenched with a final concentration of 0.5% hydroxylamine. The samples were mixed at a 1:1 ratio across all channels, based on initial ratio check analysis. Combined sample was then cleaned on a SepPak cartridge.

High-Select Fe-NTA Phosphopeptide Enrichment Kit (Thermo Fisher) was used to enrich the phosphorylated peptides (phosphopeptides) according to the manufacturer’s protocol. The eluent from the phosphopeptide enrichment was desalted and analyzed by liquid chromatography-tandem mass spectrometry (LC-MS/MS). Flow-through and washes from phosphopeptide enrichment were combined, dried, and fractionated with basic pH reversed-phase (BPRP) high-performance liquid chromatography (HPLC). We used an Agilent 1260 pump equipped with a degasser and a single wavelength detector (set at 220 nm). Peptides were subjected to a 50-min linear gradient from 8 to 40% acetonitrile in 10 mM ammonium bicarbonate pH 8 at a flow rate of 0.25 mL/min over an Agilent 300Extend C18 column (3.5 μm particles, 4.6 mm ID and 250 mm in length). The peptide mixture was fractionated into a total of 96 fractions, which were consolidated into 24 fractions. Twelve fractions were desalted and analyzed by liquid chromatography-tandem mass spectrometry (LC-MS/MS).

Mass spectrometry data were collected using an Orbitrap Exploris480 mass spectrometer (Thermo Fisher Scientific, San Jose, CA) coupled nLC-1200 liquid chromatograph. Peptides were separated on a 100 μm inner diameter microcapillary column packed with ∼35cm of Accucore C18 resin (2.6 μm, 150 Å, Thermo Fisher Scientific). For each analysis, we loaded ∼2 μg onto the column. Peptides were separated using a gradient of 5 to 29% acetonitrile in 0.125% formic acid with a flow rate of 400nL/min. The scan sequence began with an Orbitrap MS1 spectrum with the following parameters: resolution 60K, scan range 350-1350, automatic gain control (AGC) target was set to “standard,” maximum injection time was set to “auto,” and centroid spectrum data type. We used a cycle time of 1 s for MS2 analysis. Phosphoproteomics data were acquired across a 150 minutes gradient, collision energy of 36%, with compensation voltages (CV) of −30, −50, and −70 V for the first injection and of −40, −60, and −80 V for the second injection. Proteomics data were acquired for 12 of the 24 fractions across a 90 min gradient, with compensation voltages (CV) of −30, −50, and −70 V. In both analyses, we used collision energy of 36%, 45K resolution with TurboTMT activated an automatic gain control (AGC) setting of 250% and a maximum injection time of 150 ms. Dynamic exclusion was set to automatic.

#### Luciferase assays

The renilla luciferase promoter (HSV-TK) was PCR-amplified from the pGL4.74 plasmid (Promega) using primers that incorporated 5’ and 3’ *HindIII* restriction enzyme cut sites. The empty (minimal 3’UTR) construct was cloned using the Firefly luciferase reporter plasmid pGL4.11 (Promega), which was linearized using HindIII digestion followed by Gibson assembly with the HSV-TK amplified DNA sequence using HiFi Gibson Assembly Master Mix (New England Biolabs). The full 3’UTR of *Nfkbiz* (1-1355 nt), as well as a truncated *Nfkbiz* 3’UTR missing the first 165 nt (166-1355 nt), or the first 165 nt of the *Nfkbiz* 3’UTR (1-165 nt), were amplified from mouse genomic DNA using PCR primers that incorporated 5’ and 3’ *XbaI* restriction enzyme cut sites. These DNA amplicons were cloned into the HSV-TK:pGL4.11 construct described above using Gibson Assembly.

Firefly luciferase reporter constructs (*Fluc,* 500 ng plasmid) were co-transfected with the internal control pGL4.74 Renilla luciferase expression construct (*Rluc,* 10 ng plasmid) into mouse primary cultures (described above) at 5DIV using Lipofectamine 2000 (Life Technologies) following the manufacturer’s instructions. Silenced cultures were depolarized on 7DIV as above for 2 or 6 h before washing once with cold 1× PBS and collecting cells in lysis buffer. Protein lysates were analyzed using Promega reagents according to Assay System instructions on a BioTek synergy 4 microplate reader, Gen5 1.11. Firefly luciferase activity readings were normalized for each experimental replicate using the Renilla luciferase activity reading from the same sample: *Fluc*/*Rluc*. Two normalized *Fluc* values (experimental replicates) were averaged for each biological replicate value. The average normalized *Fluc* value of each condition treated with KCl was divided by the average normalized *Fluc* value of that same condition left untreated to obtain the fold-induction of the *Fluc* reporter for each condition in each biological replicate. Statistics were performed using a one-way ANOVA followed by Tukey’s post-hoc test for multiple comparisons.

#### Massively Parallel Reporter Assays

The EF1a:GFP reporter plasmid was generated by first PCR amplifying EGFP from OD1056 using primers that incorporated *XbaI* (5’) and *EcoRI* (3’) restriction enzyme cut sites. EGFP was cloned into pAAV-Ef1a-fDIO hChR2(H134R)-EYFP (Addgene #55639) digested with *XbaI* and *EcoRI* using Gibson assembly as above, resulting in the base plasmid pAAV-Ef1a-EGFP-WPRE.

In Fig. 3A, sequences were selected for library generation by tiling the 3’UTRs of transcripts expressed in mouse neurons using a sliding window of 190 nt with a 100 nt step size. These 190 nt motifs were then filtered and sorted into seven classes: Class 1 - Conserved elements (PhyloP ≥ 0.5) from activity-stabilized or -destabilized 3′UTRs (*n*=11,522); Class 2: Fast-evolving elements (PhyloP ≤ −0.5) from activity-regulated 3′UTRs (*n*=244); Class 3: Non-conserved elements (−0.5 < PhyloP < 0.5) from activity-regulated 3′UTRs (*n*=1,234); Class 4: Conserved elements from genes not regulated by activity (*n*=500); Class 5: Non-conserved elements from non-regulated genes (*n*=500); Class 6: Conserved elements from transcription-only genes (*n*=500); Class 7: GC-matched scrambled sequences from class 1 (*n*=500). A pool of the corresponding oligos flanked by adapter sequences and a randomly generated 8-nt barcode (GTACAAGTAAGAATTC-motif-barcode-GAATTCGATATCAAGC) was synthesized (Twist Bioscience or Agilent Technologies). Fragments were PCR-amplified using Q5 High-Fidelity DNA Polymerase (NEB) and cloned via Gibson assembly into the pAAV-Ef1a-EGFP-WPRE plasmid digested with *EcoRI* and *AfeI* to remove the WPRE element prior to assembly as this element has been previously shown to undergo mixed polyA tailing that could affect reporter RNA stability in cells ^89^. The resulting library was transformed into *E. coli* NEB Stable competent cells (NEB). To assess complexity and uniformity of the plasmid pool, we performed high-throughput sequencing on the cloned library.

In Fig. 5G, sequences were selected for library generation by tiling the 3’UTRs of HuD-bound mRNAs defined by RIP-seq using a sliding window of 190 nt with a 100 nt step size. These 190 nt motifs were scanned for predicted HuD-binding sites using RBPmap ^90^ and filtered for motifs that contained two predicted HuD-binding sites with ≤10 bp in between the sites. Reference sequences, as well as variant sequences in which the central 3 bp of each predicted HuD-binding site were mutated to CGC which was previously shown to disrupt HuD binding *in vitro* ^58^, were included as separate motifs in the MPRA library, as well as 250 HuD-bound mRNAs with no predicted HuD motif and 250 motifs from RNAs that were not HuD-bound mRNAs. Oligos were synthesized and cloned into pAAV-Ef1a-EGFP-WPRE as above.

In Fig. 7G, variants were selected from the Varicarta database^77^ and the NHGRI GWAS Catalog v1.0.2^78^ accessed on 4/22/2024 and filtered for 50 traits broadly related to neurodevelopmental, neuropsychiatric, mood, and brain-related disorders, such as attention deficit hyperactivity disorder (ADHD), autism spectrum disorder (ASD), schizophrenia, bipolar disorder, unipolar depression, anxiety, obsessive-compulsive disorder, mood disorders, and brain cancers (e.g., glioma, glioblastoma), as well as symptom- or behavior-based measurements such as memory performance and depressive or ADHD symptom scores. Variants were further filtered based on overlap with at least one annotated 3’UTR. Both the reference and variant sequence, as well as the surrounding 190 nt sequence context, were included as separate motifs in the MPRA library. Oligos were synthesized and cloned into pAAV-Ef1a-EGFP-WPRE as above.

#### AAV preparation

For the MPRA library shown in Fig. 4A, AAV2/9 vectors expressing the library of constructs described above were produced using triple transient transfection in HEK293T cells following established protocols.^91^ Briefly, 293T cells were seeded at 90% confluence in 150 mm dishes 24 h prior to transfection. Cells were transfected using polyethylenimine (PEI, 1 mg/mL) with a 4:2:1 ratio of helper plasmids: pAAV2/9 (Addgene # 112865), pHelper (Agilent 240071), and plasmid library per plate at a final PEI:DNA ratio of 3.5:1. Media was exchanged 12-24 h post-transfection with DMEM containing 5% FBS, GlutaMAX, and penicillin-streptomycin. Viral supernatant was harvested on day 4 and day 6 post-transfection. Cells were lysed using salt-active nuclease (SAN) buffer containing 0.5% Tween-20 and incubated at 37°C for 1 h. Viral particles were precipitated from the supernatant using polyethylene glycol 8000 (PEG, 1:5 v/v) and incubated on ice for 2 h, followed by centrifugation at 4,000g for 30 min at 4°C. The PEG pellet was resuspended in SAN buffer and combined with cell lysate for continued digestion at 37°C for 30 min. Viral particles were purified using iodixanol (OptiPrep) density gradient ultracentrifugation in a 70Ti rotor at 58,400 rpm (350,000g) for 2 h 25 min at 18°C. The 40% iodixanol fraction containing viral particles was collected and buffer-exchanged three times using Amicon Ultra-15 centrifugal filters (100 kDa MWCO) with high-salt formulation buffer (PBS with 0.001% Pluronic F-68 and 180 mM NaCl). Final viral preparations were concentrated to 200-500 μL, aliquoted in 10 μL volumes, and stored at −80°C. Viral titers were determined using both traditional EGFP-specific primer qPCR and the Takara AAV titration kit, with DNase I treatment followed by proteinase K digestion to release viral genomes prior to quantification. For the MPRA libraries shown in Fig. 5G and 7G, AAV2/9 vectors were produced by the Boston Children’s Hospital Viral Core.

#### Stereotaxic injection

All stereotaxic surgeries were performed in accordance with protocols approved by the Harvard University Standing Committee on Animal Care and federal guidelines. P21 male and female mice were anesthetized with isoflurane inhalation and positioned on a stereotaxic frame (Kopf Model 1900 Alignment System). Animal body temperature was maintained at 37°C by a heat pad and animal health (as indicated by breathing rate) was monitored carefully during the entire procedure. First, fur on the scalp of the head was trimmed using an electronic shaver and sterilized with three alternating washes of betadine and 70% ethanol. Needles for viral delivery were pulled on a Sutter Model P-2000 and cut at the very tip to generate long needles with a thin opening of approximately 50µm. The following coordinates for virus injection were used: AP −2.4, ML ± 2, DV −1.6. Virus (1 µL, 1.4×10^9^ vg/µL) was slowly injected, and the pipette was left in place for 5 min to allow for viral spreading. Postoperative analgesic slow-release Buprenorphine was administered at 50 mg/kg at the end of the surgery. Mice were monitored for four days after stereotaxic surgery for signs of infection, mobility and pain.

For MPRA library preparation, mice were IP injected with KA or PBS as described above. Six h after IP injection, mice were deeply anesthetized with isoflurane, decapitated, and brains were placed in a stainless-steel brain matrix on ice to cut 1 mm sections of anterior to posterior hippocampus. Sections were transferred into ice-cold PBS, and Cornu Ammonis (CA)1 was separated under a fluorescence dissection microscope, removing pieces of CA1 tissue where AAV did not spread (as assessed by GFP reporter fluorescence). Tissue was frozen in liquid nitrogen and stored at −80°C for further processing and DNA/RNA extraction.

For histology, mice were sacrificed and dissected brains were fixed in 4% paraformaldehyde (PFA) in PBS for 48 h at 4°C. Brains were sectioned on a vibratome LeicaVT1000 at 80 µm in PBS. Free-floating hippocampal sections were mounted on superfrost glass slides and counterstained with Fluoromount-G plus DAPI. Images were acquired with an Olympus VS200 slide scanner microscope using a 10x objective at 1024×1024 pixel resolution.

#### MPRA library preparation

For MPRA libraries in Figs. 5G and 7G, neurons were plated as described above in 24-well plates at 2×10^5^ neurons/well and cultured for 7DIV. At 3DIV, media was changed as above and AAV was added to a multiplicity of infection (MOI) of 1×10^5^ particles/neuron. At 7DIV, silenced neurons were depolarized or treated with mock depolarization buffer for 1, 2, or 6 h as above, harvested in ice-cold 1xPBS, and RNA samples were resuspended in TRIzol while DNA samples were pelleted and flash-frozen in liquid nitrogen. All samples were stored at −80°C.

For all MPRA libraries, RNA samples in Trizol were chloroform extracted once, and RNA was isolated using the Qiagen MinElute Cleanup Kit. DNA was isolated using the Qiagen Allprep Kit. Total RNA (1 µg) was treated with ezDNase (SuperScript IV VILO Master Mix Kit, Thermo Fisher Scientific) and reverse transcribed in a 20 µL reaction containing SuperScript IV VILO master mix and 2 µM gene-specific primer (GTGACTGGAGTTCAGACGTGTGCTCTTCCGATCTCGATANNNNNNNNNNNNGCTTG where N indicates randomized UMI sequence) according to the manufacturer’s instructions. An initial PCR step was performed on cDNA (2 µL) or genomic DNA (100 ng) using Q5 Master Mix (NEB) and primers (cDNA: GTGACTGGAGTTCAGACGTGTGCTCTTCGGATCTCGATAA and ACACTCTTTCCCTACACGACGCTCTTCCGATCTGCATGGACGAGCTGTACAAG; gDNA: ACACTCTTTCCCTACACGACGCTCTTCCGATCTGCATGGACGAGCTGTACAAG and GTGACTGGAGTTCAGACCGTGTGCTCTTCCGATCTCGATAAGCTTGATATCGAATTC). Reactions were purified with 1.2× SPRI beads and eluted in 23 µL water. A barcoding PCR step was carried out with 5 µL of purified product using Clontech RNA-seq barcoding primers and DNA polymerase (Takara), yielding a 334 nt product, followed by cleanup with 1.2 volumes of SPRI beads (Agilent) according to the manufacturer’s instructions. A final amplification step was performed using 5 µL of template with Clontech RNA-seq PCR2 primers and DNA polymerase as above. Final products were purified using SPRI beads, eluted into 10 µL nuclease-free water, and quantified by Agilent Bioanalyzer, which showed a single peak at ∼334 nt. Libraries were sequenced using paired-end 2×150-nt cycles on a Novaseq 24B instrument to a depth of at least 20M reads per sample.

#### RNA immunoprecipitation

Neurons were plated as described above in 10 cm dishes at 1×10^7^ neurons/dish and cultured for 7DIV. At 7DIV, silenced neurons were depolarized or treated with mock depolarization buffer for 1, 2, or 6 h as above, harvested in ice-cold 1X PBS, flash frozen in liquid nitrogen, and stored at −80°C. Cells were lysed in 100 µL lysis buffer (20 mM Tris pH 7.5, 100 mM KCl, 5 mM MgCl_2_, and 0.5% NP-P40 (Life Technologies # 28324)) supplemented with 0.5X Halt Protease and Phosphatase Inhibitor (Life Technologies # 78442) and 0.4 U/µL RiboLock RNase Inhibitor (Thermo Scientific # EO0381), and incubated on ice for 5 min. Lysates were centrifuged at 14,000 rpm for 10 min at 4℃. Supernatant (100 µL) was collected, reserving 10% of lysate as input. Dynabeads Protein G beads (Life Technologies #10004D, 50 µL/sample) were washed twice with 500 µL ice-cold wash buffer (50 mM Tris pH 7.5, 150 mM NaCl, 1 mM MgCl_2_, 0.05% NP-P40) and incubated with 500 ng antibody (FUS, Santa Cruz Biotechnology SC-47711; HuR, Santa Cruz Biotechnology 5261; CELF4, Sigma Aldrich HPA037986; HuD, Life Technologies 24992-1-AP; and IgG, Santa Cruz Biotechnology 2025) in 100 µL of wash buffer and incubated for 30 min at 4°C with rotation. Beads were washed twice with 500 µL wash buffer and resuspended in 900 µL IP buffer (Wash buffer supplemented with 20 mM EDTA and 5 µL RiboLock RNase Inhibitor). 100 µL of lysate was added and beads were incubated for 3 h at 4℃ with rotation.

Beads were washed six times with 500 µL ice-cold wash buffer, then resuspended in 150 µL elution buffer (wash buffer supplemented with 1% SDS and 360 µg Proteinase K (Qiagen 19131). Beads were incubated at 55℃ for 30 min shaking at 300 rpm to digest the protein. The supernatant of each sample was then transferred to a MaXtract High Density tube (Qiagen # 129056) containing 250 µL of wash buffer and 400 µL of UltraPure Phenol:Chloroform:Isoamyl Alcohol (Life Technologies # 15593031) and was vortexed for 15 s before centrifugation at 12,000*g* for 10 min at room temperature to separate the aqueous phase. This step was repeated using 350 µL of the aqueous phase and 400 µL of Chloroform (Sigma Aldrich # C2432). RNA was ethanol precipitated as above and resuspended in a final volume of 10 µL of RNase-free water.

RNA quality and concentration were assessed by RNA BioAnalyzer (Agilent), and samples were excluded if the amount of RNA immunoprecipitated was less than the IgG control. Each batch of samples always included at least one IgG control sample. Libraries were prepared using the SMARTer Stranded Total RNA-seq Kit - Pico Input Mammalian V2 (Takara Bio 634413) according to the manufacturer’s instructions. Samples were multiplexed with Illumina barcodes and sequenced using paired-end 2×75-nt cycles on an Illumina NextSeq 500 instrument to a depth of at least 20M reads per sample.

#### shRNA cloning and lentiviral preparation

For shRNA treatments, 21 bp targeted sequences from TRC (Sigma Aldrich) were cloned into the FUW vector downstream of the U6 promoter using *PacI* restriction sites. The following sequences were used: Control shRNA (GCGCGATAGCGCTAATAATTT), *Elavl4* shRNA (GTGTTGCAAGTTTCCTTTAAA).

Lentiviral vectors were produced using Lenti-X 293T cells (Takara 632180) through calcium phosphate transfection under BSL-2 containment conditions. Lenti-X 293T cells (5-6 × 10⁶) were seeded on poly-ornithine-coated 10 cm dishes in MEF media (DMEM supplemented with 10% calf cosmic serum, 1% penicillin/streptomycin, 1% non-essential amino acids, and 1% sodium pyruvate) and grown to ∼80% confluence. For transfection, cells were co-transfected with 10 μg shRNA plasmid, 5 μg pMDLg/pRRE, 2.5 μg pRSV-rev, and 2.5 μg pMD2.G using calcium phosphate precipitation. Briefly, plasmid DNA was mixed with 2.5 M CaCl₂ in sterile water (500 μL total volume), combined with an equal volume of 2× BBS buffer (50 mM BES, 280 mM NaCl, 1.5 mM Na₂HPO₄, pH 6.95), and incubated for 30 min at room temperature before dropwise addition to cells. Media was replaced 16-20 h post-transfection with low-serum MEF media (5% calf cosmic serum). Transfection efficiency was monitored using a TetO-EGFP control plasmid, with >60% EGFP-positive cells considered acceptable for viral production. Viral supernatant was harvested 40-44 h post-transfection, filtered through 0.45 μm filters to remove cellular debris, and concentrated using polyethylene glycol precipitation (1:4 v/v ratio with 4× PEG concentrator solution containing 40% PEG-8000, 1.2 M NaCl, and 1× PBS). Following overnight incubation at 4°C, viral particles were pelleted by centrifugation at 1,500*g* for 45 min at 4°C. The pellet was resuspended in 200 μL DMEM, aliquoted, flash frozen in liquid nitrogen, and stored at −80°C until later use.

#### Separation chambers

Neurons were plated as described above in 6-well plates with Millicell cell culture inserts (PISP30R48 13 μm, Millipore) at 9×10^5^ neurons/well and cultured for 7DIV. At 7DIV, silenced neurons were depolarized for 15 min, 30 min, 1 h, 2 h, or 6 h by the addition of depolarization buffer as above, as well as a mock depolarization control. At time of collection wells were washed once with cold PBS. Samples were then collected using 250 µL of RIPA buffer (Sigma Aldrich) using a cell lifter (3010 Corning) first from the neurite side of the insert and then from the soma side. RNA was extracted with Trizol LS (CAT 10296010, Thermofisher) according to the manufacturer’s instructions.

#### Spatial transcriptomics

Mice were injected with kainic acid or PBS as described above and harvested 2 or 6 h after injection. Whole brain was removed and embedded in OCT, flash frozen on dry ice, and stored at −80°C. OCT-embedded brain tissue was cryo-sectioned at a thickness of 10 μm at −21°C using the CM1950 Leica cryostat and immediately mounted onto a 10×10 mm Slide-seq V2 chip. Libraries were prepared using a Slide-SeqV2 kit (Curio Biosciences) according to the manufacturer’s instructions, and libraries were sequenced on a Novaseq X flowcell 10B (paired-end, 2×150 cycles).

#### HuD shRNA knockdown and TimeLapse-seq

Neurons were plated as described above in 24-well dishes at 2×10^5^ neurons/well and cultured for 7DIV. At 3DIV, 50% of the culture media was exchanged for fresh media supplemented with 8 µL lentivirus. At 7DIV, silenced neurons were depolarized for 1, 2, or 6 h by the addition of depolarization buffer as above, as well as a mock depolarization control. s^4^U was added to a final concentration of 500 µM for 4 h before neurons were harvested, and additional s^4^U was added with depolarization buffer to maintain a concentration of 500 µM. Neurons were washed twice with 1X PBS to remove dead cells and scraped immediately into Trizol (Life Technologies 15596026). RNA isolation and TimeLapse-chemistry were performed as previously described.^55^ Briefly, cells lysed in Trizol were chloroform-extracted once, and nucleic acids were precipitated using isopropanol. DNA was removed using Turbo DNase (Life Technologies AM1907), and RNA was purified using Agencourt RNAClean XP beads (Fisher Scientific NC0068576). One volume of RNAClean beads were added to each sample, incubated at room temperature for 8 min, and captured on a magnetic rack. Beads were washed twice with 80% ethanol, and RNA was eluted from dried beads using RNase-free water.

Isolated total RNA was added to a mixture of 2,2,2-trifluoroethylamine (TFEA, 600 mM), EDTA (1 mM) and sodium acetate (pH 5.2, 100 mM) in water. A solution of NaIO4 (10 mM) was then added dropwise, and the reaction mixture was incubated for 1 h at 45°C. RNA was purified using RNAClean beads as above, resuspended in nuclease-free water, and reducing buffer was added to a final concentration of 10 mM DTT, 10 mM Tris pH 7.4, 1 mM EDTA, and 100 mM NaCl and incubated for 30 min at 37°C. RNA was purified using RNAClean beads as above, resuspended in nuclease-free water, and RNA concentration was assayed by Nanodrop. RNA-seq libraries were prepared from 20 ng of chemically treated RNA using the SMARTer Stranded Total RNA-seq Kit – Pico Input Mammalian V2 (Takara Bio 634413) according to the manufacturer’s instructions. Samples were multiplexed with Illumina barcodes and sequenced using paired-end 2×150-nt cycles on a Novaseq 24B instrument to a depth of at least 20M reads per sample.

#### HuD immunoprecipitation and mass spectrometry

HuD-FLAG-HA and EGFP-FLAG-HA dsDNA fragments were synthesized by Twist Bioscience and cloned into an FUW vector that was linearized using *EcoRI* and *NheI* restriction enzyme digestion followed by Gibson assembly using HiFi Gibson Assembly Master Mix (New England Biolabs). Lentiviral vectors were prepared as above from FUW-HuD-FLAG-HA and FUW-EGFP-FLAG-HA plasmids separately.

Neurons were plated as described above in 10-cm dishes at 1×10^7^ neurons/well and cultured for 7DIV. At 3DIV, 50% of the culture media was exchanged for fresh media supplemented with lentivirus (100 µL HuD-FLAG-HA and 25 µL EGFP-FLAG-HA). At 7DIV, silenced neurons were depolarized for 1 or 6 h by the addition of depolarization buffer as above, as well as a mock depolarization control, harvested in ice-cold 1X PBS, flash frozen in liquid nitrogen, and stored at −80°C. Two 15-cm plates were used per biological replicate and lysed in MCLB buffer (50 mM Tris pH 7.5, 150 mM NaCl, 0.5% NP-40) supplemented with protease inhibitors (Roche) and 1 mM DTT. Cell pellets were lysed by tumbling for 20 min at 4°C, followed by centrifugation at 16,100 × g for 20 min to clear the lysate. Cleared lysates were incubated with 80 μL of pre-washed anti-FLAG magnetic beads (Sigma) for 3 h with gentle rotation at 4°C. Following immunoprecipitation, beads were washed three times with 1 mL MCLB buffer, then three additional times with detergent-free buffer (50 mM Tris pH 7.5, 150 mM NaCl) to remove excess detergent prior to mass spectrometry analysis. Bound proteins were eluted twice with 20 μL elution buffer (1% formic acid) at room temperature for 10 min with gentle agitation. Eluates were combined and concentrated by speed vacuum evaporation, then split with 20% reserved for western blot analysis and 80% processed for mass spectrometry analysis.

Samples were resuspended in 200 mM EPPS (pH 8.5) and digested at a 1:100 protease-to-peptide ratio with Lys-C overnight at room temperature, followed by Trypsin digestion for 6 h at 37°C. Following digestion, acetonitrile was added to the eluted peptides for a final volume of 30%. Peptides were labeled with the tandem mass tag (TMT) pro18-plex reagent for 1 h. The reaction was quenched with a final concentration of 0.5% hydroxylamine. The samples were combined, desalted, and analyzed by LC-MS/MS.

Mass spectrometry data were collected using an Exploris 480 mass spectrometer (Thermo Fisher Scientific, San Jose, CA) coupled with a Proxeon 1200 Liquid Chromatograph (Thermo Fisher Scientific). Peptides were separated on a 100 μm inner diameter microcapillary column packed with ∼35 cm of Accucore C18 resin (2.6 μm, 150 Å, Thermo Fisher Scientific). We loaded ∼1 μg onto the column. Peptides were separated using a 90 minutes gradient of 5 to 30% acetonitrile in 0.125% formic acid with a flow rate of 320 nL/min. The scan sequence began with an Orbitrap MS1 spectrum with the following parameters: resolution 60,000, scan range 350−1350 Th, automatic gain control (AGC) target “standard”, maximum injection time “auto”, RF lens setting 50%, and centroid spectrum data type. We selected the top twenty precursors for MS2 analysis, which consisted of HCD high-energy collision dissociation with the following parameters: resolution 15,000, AGC was set at “standard”, maximum injection time “auto”, isolation window 0.7 Th, normalized collision energy (NCE) 28, and centroid spectrum data type. A TopSpeed setting of 1 s was used. In addition, unassigned and singly charged species were excluded from MS2 analysis and dynamic exclusion was set to 90 s.

#### Immunoblotting

Whole-cell or soma/neurite fractions from primary neurons were harvested in RIPA buffer (Sigma Aldrich # R0278) and snap-frozen in liquid nitrogen, followed by storage at −80°C. Protein lysates were thawed on ice and incubated with 250 U benzonase (Sigma Aldrich # E8263) for 30 min at room temperature. Neurite fractions from primary neurons were concentrated using Pierce Protein Concentrators (Life Technologies # 88513). Digested lysates were mixed with NuPAGE LDS sample buffer to 1x (Invitrogen NP0007), heated to 70°C for 10 min, and resolved on 4%-12% Bis-Tris gels and transferred to nitrocellulose. Membranes were blocked in 5% milk and incubated overnight in the following primary antibodies: HuD (1:1000, Life Technologies # 24992-1-AP), HuR (1:1000, Santa Cruz Biotechnology 5261), CELF4 (1:500, Sigma Aldrich HPA037986), FUS (1:1000, Santa Cruz Biotechnology SC-47711), HA (1:1000, CST #3724), Histone H3 (1:10,000), ꞵ-Tubulin (1:5000). Following washing, membranes were incubated with secondary antibodies conjugated to HRP (anti-Rabbit, 1:10,000, CST #7074; anti-Mouse, 1:1000, CST #7076S), developed with SuperSignal West Pico PLUS or Atto Ultimate Chemiluminescent Substrate (Life Technologies # 34578 and #A38556, respectively) and imaged with a BioRad ChemiDoc MP System.

#### Immunofluorescence

Neurons were plated as described above on glass coverslips at 8×10^4^ neurons/coverslip and cultured for 7DIV. At 7DIV, silenced neurons were depolarized for 1 or 6 h by the addition of depolarization buffer as above, as well as a mock depolarization control. Cells were washed once with ice-cold 1xPBS and fixed with 4% paraformaldehyde in 1x GDB buffer (1xPBS supplemented with 0.1% gelatin and 0.3% Triton X-100) for 30 min at room temperature and washed three times with ice-cold PBS. The cells were permeabilized and blocked for 1 h at room temperature using 5% donkey serum in PBST (1× PBS and 0.1% Triton X-100). Coverslips were incubated with anti-HuD rabbit primary antibody (1:1000, Life Technologies # 24992-1-AP) and anti-MAP2 chicken primary antibody (1:2000, Lifespan Biosciences, LS-C61805) overnight at 4 °C. Coverslips were washed 3 times in PBST at room temperature and then incubated with fluorescently labeled secondary antibody (1:2000, Alexa Fluor 555 anti-rabbit and Alexa Fluor 488 anti-chicken) for 1 h at room temperature. Coverslips were washed 3× in PBST at room temperature and mounted onto Superfrost glass slides using DAPI Fluoromount-G (Thermo Fisher Scientific). Images were visualized using a Leica Stellaris confocal microscope equipped with a 405 nm diode and white light laser tuned to 488, 555, and 647 nm with appropriate filter sets to minimize spectral crosstalk. Images of HuD puncta were taken using a Leica HC PL APO 100x/1.44 OIL objective at 1X optical zoom (voxel size: 0.1136 x 0.1136 x 0.4985 µm^3^) with 2x line averaging and a pinhole size of 0.8 AU at 580 nm in 16-bit depth. Z-stacks of approximately 7-15 slices were taken to include the bottom and top of cultures.

### QUANTIFICATION AND STATISTICAL ANALYSIS

#### RNA-seq analysis

Illumina adapters were trimmed with Cutadapt (version 1.14) and aligned using Hisat2^92^ (version 2.1.0) to the *Mus musculus* genome (GRCm39) and transcriptome (Ensembl). Alignments and analysis were performed on the Orchestra2 high performance computing cluster through Harvard Medical School. Aligned BAM files were sorted using Picard Tools (version 2.8.0); stranded bedGraphs were generated using STAR^93^ (version 2.7.0f); and reads were quantified over annotated exons and gene bodies using FeatureCounts ^94^ (Subread version 2.0.6). Lowly expressed genes were filtered using the filterByExpr function in edgeR (version 4.0.16). Intronic reads were calculated by subtracting reads over exons from reads over gene bodies. Differential expression analysis was performed using DESeq2.^95^ Significant genes were defined as P_adj-_ < 0.05 and abs(log_2_(fold change)) > log_2_(1.2). Depth-normalized counts were generated using DESeq2. K-means clustering was performed in R using the kmeans function, with the maximum number of iterations set to 10,000. Gene Ontology (GO) enrichment analysis was performed using gProfiler2 in R (version 0.2.3), with a custom background of expressed genes based on expression-filtered RNA-seq genes and false discovery rate (FDR) < 0.05. Heatmaps were generated using ComplexHeatmap^96^ (version 2.18.0). Synaptic genes were defined using the SynGO database.^97^

#### Calculating RNA synthesis, processing and degradation rates

Rates of RNA synthesis, processing and degradation were calculated from pulse s^4^U-seq data using INSPEcT^33^ (version 1.38.0) assuming that RNA degradation is possible during the s^4^U pulse (degDuringPulse=TRUE). To estimate variance around the RNA kinetic rates, we employed a leave-one-out jackknife approach. With four biological replicates per condition, we systematically re-ran the INSPEcT analysis excluding one replicate at a time, generating four jackknife estimates for each rate parameter. The variance for each RNA rate was then calculated from these jackknife estimates using standard jackknife variance estimation procedures. Differential RNA rate analysis was performed using z-score based statistical testing. For each gene and time point comparison (t vs. time 0), log_2_ fold changes were calculated for each jacknife replicate, and the mean difference across replicates was tested using z-scores. P-values were adjusted for multiple testing using the Benjamini-Hochberg procedure within each rate type and time comparison. Genes were classified as significantly changed if they met dual criteria: P_adj_ < 0.05 and absolute log_2_ fold change > log_2_(1.2).

#### Ribosome profiling

Sequencing adapters were removed using Cutadapt (version 1.14); trimmed FASTQ files were aligned to mm10 ribosomal RNA sequences using Bowtie2 (version 2.3.4.3); and unaligned reads were mapped to the *Mus musculus* genome (GRCm39) and transcriptome (Ensembl) using STAR (version 2.7.3a) with standard settings and the following modified parameters: – clip5pNbases 3, –seedSearchStartLmax 15, –outSJfilterOverhang-Min 30 8 8 8, –outFilterScoreMin 0, –outFilterScoreMinOverLread 0.66, –outFilterMatchNmin 0, –outFilterMatchNminOverLread 0.66, –outSAMtype BAM Unsorted. Aligned BAM files were filtered for only uniquely mapped reads and sorted using Picard Tools (version 2.8.0). The RibORF pipeline was run on each sample individually using standard param-eters. Due to template switching during library preparation, reads contained three untemplated bases at the 3′ end that were not included in the alignment but added to the length of each read. Therefore, reads 30–33 nt in length (corresponding to RNA fragments 27–30 nt) were analyzed for three-nucleotide periodicity within known protein-coding ORFs (RefSeq). For each sample, we selected only the read lengths for which at least 50% of the reads matched the primary ORF of known protein-coding genes in a meta-gene analysis. Read lengths were offset-corrected and resulting P-site reads were used to quantify Ribo-seq expression using FeatureCounts^94^ (Subread version 2.0.6). Activity-dependent changes in ribosome density were calculated by two--way differential expression analysis between Ribo-seq P-sites and total RNA-seq using deltaTE^98^ in R version 4.2.1. Read normalization and size factor estimation were performed on RNA-seq and Ribo-seq data simultane-ously; samples were corrected for batch effects.

#### Mass spectrometry data analysis

Mass spectrometry data were processed using a Comet-based pipeline. Spectra were converted to mzXML using a modified version of ReAdW.exe. Database search included all entries for mouse from UniProt, in addition to GFP sequence. This database was concatenated with one composed of all protein sequences in the reversed order. Searches were performed using a 50-ppm precursor ion tolerance. TMT tags on lysine residues, peptide N termini (+304.207 Da), and carbamidomethylation of cysteine residues (+57.021 Da) were set as static modifications, while oxidation of methionine residues (+15.995 Da) was set as a variable modification. For phosphopeptide analysis, phosphorylation (+79.966 Da) on serine, threonine, and tyrosine was included as variable modification. Peptide-spectrum matches (PSMs) were adjusted to FDR < 0.01. PSM filtering was performed using a linear discriminant analysis (LDA), as described previously, while considering the following parameters: XCorr, ΔCn, missed cleavages, peptide length, charge state, and precursor mass accuracy. For TMT-based reporter ion quantitation, we extracted the summed signal-to-noise (S:N) ratio for each TMT channel and found the closest matching centroid to the expected mass of the TMT reporter ion. For protein-level comparisons, PSMs from all samples were identified, quantified, and collapsed to a peptide FDR < 0.01 and then collapsed further to a final protein-level FDR < 0.01, which resulted in a final peptide level FDR <0.001. Moreover, protein assembly was guided by principles of parsimony to produce the smallest set of proteins necessary to account for all observed peptides. PSMs with poor quality, TMT reporter summed signal-to-noise of less than 100, or isolation purity lower than 50% were excluded from quantification.

##### Whole cell proteomics and phosphoproteomics

With the TMT quantification we evaluated changes in protein levels and phosphorylation. In summary, we extracted the relative protein levels across TMT channels and for each protein and time point, calculated the log_2_ fold-change (FC) relative to time zero was calculated for each replicate. A z-score was computed for the mean log_2_FC across replicates, divided by the standard deviation. Proteins were classified as significantly increased (UP) or decreased (DOWN) if the BH-adjusted P value (P_adj_) < 0.05 and the absolute log_2_FC > log_2_(1.2).

##### IP-MS enrichment analysis

The TMT signal from the IP experiment was normalized by total protein levels. Then we applied a replicate-based method, where a z-score analysis was implemented. For each protein, the log_2_FC was calculated for each individual biological replicate pair (HuD/GFP for interactors at each time of KCl depolarization). The mean log_2_FC across replicates was then divided by the standard deviation of those log_2_FC values to generate a z-score. A two-tailed P value was derived from the z-score and adjusted for multiple testing using BH correction. To account for differences in IP efficiency across conditions, we normalized the HuD-enriched proteins’ fold change by the fold change of HuD protein levels. Proteins were classified into functional categories based on their temporal enrichment patterns: 1) Core interactors: Significantly enriched at all time points (0, 1, and 6 h). 2) Activity-specific interactors: Enriched only at stimulated time points (1 h and/or 6 h) but not at baseline. 3) Time-specific interactors: Uniquely enriched at individual time points. Proteins with fewer than 5 unique peptides, immunoglobulin proteins, and mitochondrial proteins (identified using MitoCarta2.0 database) were excluded from downstream analyses to reduce non-specific binding artifacts.

#### Dynamics vs. constant RNA rate modeling

To evaluate whether RNA rates remain constant or change dynamically during stimulation, we implemented a simplified two-compartment ordinary differential equations (ODE) model:

#### Dynamic Rate Model

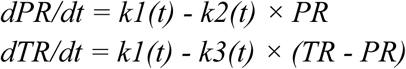

Where PR is pre-mRNA, TR is total RNA, k1 is synthesis rate, k2 processing rate, and k3 degradation rate.

#### Fixed synthesis, processing, or degradation rates

Same equations with fixed rates at 0 h KCl.

Rate functions for the dynamic model were created using linear interpolation between observed time points. Models were fit using the deSolve R package with the “lsoda” integration method to account for stiff ODE systems. To compare performance between models (dynamic versus constant) we used Akaike Information Criterion (AIC), with lower AIC indicating better model fit, and filtering for genes with ΔRMSE (dynamic - constant) < −0.5.

#### Novel environment RNA-seq analysis

Total RNA-seq data from mouse hippocampus CA1 following 30 min novel environment exposure was obtained from Traunmüller et al.^32^ The rate of RNA transcription, as well as intron half-life and mRNA half-life were calculated from RNA-seq timecourse data without metabolic labeling using INSPEcT- (version 1.38.0)^39^ (labeling_time=NULL, nascentExpressions=NULL). To estimate variance around the RNA kinetic rates, we employed a leave-one-out jackknife approach. With four biological replicates per condition, we systematically re-ran the INSPEcT analysis excluding one replicate at a time, generating four jackknife estimates for each rate parameter. The variance for each RNA rate was then calculated from these jackknife estimates using standard jackknife variance estimation procedures.

To enable direct comparison between *in vivo* data (INSPEcT-) and our in vitro metabolic labeling experiments, we restricted analysis to genes for which rate estimates were consistent between INSPEcT+ (s^4^U metabolic labeling) and INSPEcT- (label-free) models applied to the same *in vitro* dataset from primary mouse neurons depolarized with KCl. Specifically, we computed correlations for total RNA, pre-mRNA, transcription, intron half-life, and mRNA half-life across overlapping time points (0, 15, and 30 min; 1, 1.5, 2, 4, 6, and 8 h). Genes with correlation coefficients R² > 0.6 and significance p < 0.05 were retained for downstream analysis of the *in vivo* novel environment dataset.

Differential RNA rate analysis was performed using z-score based statistical testing. For each gene and time point comparison (t vs. time 0), log_2_ fold changes were calculated for each jacknife replicate, and the mean difference across replicates was tested using z-scores. P-values were adjusted for multiple testing using the Benjamini-Hochberg procedure within each rate type and time comparison. Genes were classified as significantly changed if they met dual criteria: P_adj_ < 0.05 and absolute log_2_(FC) > log_2_(1.2). Finally, overlap between *in vitro* and *in vivo* datasets was assessed for each RNA rate. A gene was considered overlapping if it showed a significant change in both datasets and the direction of log₂ fold change was consistent across conditions.

#### Separation chamber RNA-seq analysis

To evaluate transcriptomic differences between soma and neurites, we compared RNA counts in both compartments using DESeq2. To then calculate local changes in RNA levels we compared each subcellular compartment (soma or neurite) at each time versus basal conditions (time 0). RNA transcription rates and stability measurements from metabolic labeling experiments were integrated with subcellular expression data. Genes were filtered to include only those with significant activity-dependent changes in RNA kinetics (P_adj_ < 0.05 and absolute log_2_(FC) > log_2_(1.2)). Genes were required to pass expression filtering with at least one condition showing detectable expression. Additional k-means clustering was performed on the integrated dataset to identify coordinated regulation patterns across transcriptional and post-transcriptional mechanisms. Combined heatmaps were generated using the ComplexHeatmap package.

#### Slide-seqV2 analysis

FASTQ files generated from Slide-seqV2 libraries were processed using the Curio Seeker bioinformatics pipeline (v3.0.0; Curio Biosciences), implemented in Nextflow on a Slurm HPC cluster with Singularity containerization. Each sample included paired FASTQ files and the corresponding bead barcode whitelist for each tile. Barcodes were first extracted from read 1 to obtain bead and molecular barcodes. Barcodes were next matched against the whitelist, allowing Hamming distance ≤2. UMI deduplication was performed with UMI-tools (directional mode, Hamming distance ≤1) and reads were aligned to the *Mus musculus* reference genome (Ensembl GRCm38, provided by Curio Biosciences) using STAR, followed by gene quantification using FeatureCounts.

Raw count matrices and spatial coordinate files were imported using the spacexr R package (v2.0). For each sample, gene expression counts were combined with matched bead location coordinates to create SpatialRNA objects. The number of unique molecular identifiers (nUMI) per spatial location was calculated as the sum of counts across all genes for quality assessment. Cell type deconvolution was performed using the BICCN (Brain Initiative Cell Census Network) hippocampal formation single-cell RNA sequencing atlas as reference.^99^ Robust Cell Type Decomposition (RCTD) analysis was performed using the spacexr package with doublet mode enabled to identify locations containing mixtures of cell types. Gene expression matrices from SlideSeqV2 data were intersected with the reference dataset, retaining only genes present in both datasets. Cell type assignments were classified into four categories: singlet (single cell type), doublet_certain (high confidence mixture), doublet_uncertain (low confidence mixture), and reject (insufficient information). For downstream analysis, reject locations were excluded from further processing.

Individual sample results were integrated into Seurat objects using Seurat v5 framework (version 5.3.0). For each sample, count matrices were combined with RCTD metadata and spatial coordinates were incorporated using the SlideSeq class. SCTransform normalization was applied to account for technical variation and improve downstream analysis. Principal component analysis (PCA) was performed on variable features, with optimal dimensionality determined using the elbow method combined with two criteria: (1) principal components contributing >5% of standard deviation until 90% cumulative variance was reached, and (2) consecutive PC variance change <0.1%. UMAP dimensionality reduction and Louvain clustering (resolution = 0.5) were performed using the optimal number of principal components. Multiple samples were integrated using canonical correlation analysis (CCA) with SCTransform normalization. Sample-specific prefixes were added to cellular barcodes to maintain unique identifiers across the integrated dataset. For locations classified as doublets, cell type weights from RCTD doublet mode analysis were incorporated. A consensus cell type assignment was created by selecting the cell type with the highest weight for doublet locations, while retaining original assignments for singlet locations. This resulted in a unified “cellsubtype” annotation across all spatial locations.

Spatial plots were generated using the SpatialDimPlot function to visualize cell type distributions across tissue sections. Cell type occurrence frequencies and doublet co-occurrence patterns were analyzed and visualized using the spacexr package. All statistical analyses and visualizations were performed in R (version 4.0+) using the following packages: spacexr, Seurat, Matrix, ggplot2, dplyr, and biomaRt.

We evaluated the changes in gene expression by cell type at 2 and 6 h KA treatment using a pseudobulk approach. To this end we used the AggregateExpression function from Seurat grouping by cell type, treatment (KA or PBS), and replica to obtain the sum expression of genes by cell type. We then performed differential gene expression between each KA treatment (2 or 6 h) versus PBS using DESeq2. Differentially expressed genes were defined as P_adj_ < 0.05 and absolute log_2_(FC) > log_2_(1.5).

To define local regulation *in vivo*, we manually annotated somata and neuropil regions within hippocampal CA1. The soma layer was identified as the densely packed band of pyramidal cell bodies, while the neuropil was defined as the intervening space between CA1 and dentate gyrus somata, oriented perpendicular to the soma layer. Regions were delineated using CellChat in a custom RShiny application.^100^ Gene expression patterns of regional markers were visualized, and plotly-based lasso tools were used for manual outlining, enabling separation of soma and neuropil compartments based on both morphological and molecular criteria. Selected spatial coordinates were exported for downstream comparative analysis.

Glial contamination was minimized by removing genes specifically enriched in glial cells. Glia-enriched genes were identified using reference single-cell RNA-seq data from Saunders et al.^101^ comparing neuronal versus glial metacells from hippocampus with the criteria: P_adj_ < 0.05 and log_2_(FC) < −log_2_(1.5) (glial-enriched). We performed differential gene expression in the somata versus neuropil using pseudobulk across subcellular compartment and treatment (KA vs PBS) with DESeq2 with thresholds P_adj_ < 0.05 and absolute log_2_(FC) > log_2_(1.5).

We compared the genes subcellularly regulated *in vivo* and *in vitro* by overlapping these results with the results from the separation chambers. Venn diagrams were generated using the ggvenn package to visualize genes’ overlap. Fisher’s exact tests were performed to assess enrichment of differentially expressed genes between platforms.

#### Massively Parallel Reporter Assay analysis

Paired-end 150 bp reads were merged into single amplicons using Flash v1.2.11 (flags: -M 150, -O).^102^ Adapter sequences were trimmed using cutadapt (v1.14) with the following adapters: 5’ adapter: GTACAAGTAAGAATTC, 3’ adapter: GAATTCGATATCAAGC. Trimmed reads were aligned to the reference library using BWA-MEM (v0.7.8) with stringent parameters to prevent mismatches and indels (alignment penalties: -B 100, -O 100, -T 100). Only perfectly matched reads were retained for downstream analysis. Aligned reads were processed with samtools (v1.9) to filter unmapped reads.

Principal component analysis (PCA) was performed on the top 500 most variable MPRA motifs to assess sample clustering and detect potential outliers. One sample (KA-treated male mouse, replicate 1) was excluded from further analysis because KA treatment did not induce neuronal activity (**Fig. S6D**). Elements were retained for differential expression analysis only if they had at least one count in two or more RNA samples and were present in the corresponding DNA libraries. Steady-state RNA stability was analyzed using DESeq2 with the experimental design ∼ Sex + Type + KA, where *Type* distinguished RNA from DNA samples and *KA* indicated treatment condition. Size factors were estimated using the default DESeq2 method, and differential expression was assessed using the Wald test. Statistical significance was defined as an adjusted p-value < 0.1.

To account for DNA input variation, we performed all other MPRA analyses using MPRAnalyze.^103^ A negative binomial distribution was selected over a log-normal model based on Akaike Information Criterion (AIC) comparison and goodness-of-fit testing with the fitdistrplus package. Depth factors were estimated separately for DNA and RNA libraries using upper quartile normalization. The experimental design included an intercept-only model for DNA (∼ 1) and a full model for RNA (∼ Sex + KA). Treatment effects were evaluated using both likelihood ratio tests and coefficient testing. Random control sequences incorporated into the library were used to validate normalization and statistical modeling approaches.

To identify regulatory motifs, 3′UTR sequences were systematically tiled with overlapping 7-nucleotide windows at single-nucleotide resolution. For each unique 7-mer, mean log_2_(fold change) was calculated across all UTRs containing that sequence. Statistical significance was assessed using two-sided t-tests comparing the 7-mer–containing subset against the global distribution, with p-values adjusted using the Benjamini–Hochberg procedure. Significant 7-mers (adjusted p-value < 0.1) were clustered by hierarchical clustering based on Levenshtein distance. For steady-state analyses, significant 7-mers were grouped into 10 clusters using hierarchical clustering and represented as position weight matrices (PWMs) with the *Biostrings* package. Sequence logos for each cluster were generated using ggseqlogo. To link MPRA results with endogenous RNA behavior, log_2_(fold changes) across all library elements per transcript were averaged and correlated with RNA half-life measurements obtained from metabolic labeling experiments. Differences across groups were evaluated with Kruskal–Wallis tests followed by Dunn’s post hoc correction. All visualizations were generated using ggplot2. Unless otherwise noted, p-values were adjusted for multiple testing using the false discovery rate (FDR).

Counts from the MPRA libraries containing reference and SNV/HuD mutant 3′UTR sequences were merged across six biological replicates and time points, normalized for sequencing depth, and scaled to DNA input to control for library representation. A pseudocount of 1 was added to all elements prior to normalization, and elements lacking detectable expression were excluded. Normalized RNA/DNA ratios were analyzed using MPRAnalyze as above, with models including SNP status, KCl time point, and batch as design factors. Depth factors were estimated separately for DNA and RNA libraries, and SNP and activity-dependent effects were assessed by likelihood ratio testing with FDR correction. Reporter elements were categorized by whether they exhibited significant SNP effects, activity-dependent regulation, both, or neither.

#### RIP-seq analysis

To define RBP-bound transcripts, read counts from IP samples were compared to input mRNA-seq using DESeq2^95^ with the design ∼ Time + Type, where *Type* distinguished IP from input and *Time* accounted for KCl treatment duration. Transcripts were considered RBP targets if they showed at least two-fold enrichment in IP versus input with P_adj_ < 0.05. Any transcripts enriched in the IgG versus input comparison were removed to exclude nonspecific binding. To assess activity-dependent changes in RBP binding, separate DESeq2 models were fit for each RBP across time points of KCl stimulation using the design ∼ KCl. Pairwise contrasts (e.g. 1 vs 0 h KCl) were extracted from these models, and genes meeting significance thresholds (P_adj_ < 0.05) were classified as showing increased or decreased binding. For visualization, significant genes were intersected with the set of RBP targets defined above.

#### Direct RNA sequencing analysis and PolyA length quantification

Direct RNA sequencing reads were basecalled and aligned using **Dorado v0.7.0** with default settings, the optional parameter −-estimate-poly-a, and the alignment options −-mm2-opts “-k 14 -Y -ax splice -uf” against the mouse reference genome (mm39). Poly(A) tail lengths were extracted from individual reads using awk. To compare conditions (0 h vs. 1 h KCl treatment), we applied the Wilcoxon rank-sum test to evaluate differences in poly(A) length distributions by gene, followed by Benjamini–Hochberg correction to control the false discovery rate (FDR). Effect sizes were calculated as (i) the difference in mean poly(A) length (nt) between 1 and 0 h KCl and (ii) the fold change of mean poly(A) length between conditions. Genes were considered significant if they met the thresholds of FDR < 0.05, an absolute change in mean poly(A) length > 30 nt, and an absolute fold change > 1.1.

#### Machine learning

##### Feature engineer

We created an interpretable machine learning model using linear regression to predict the fold change of mRNA half-life at each time of membrane depolarization. Comprehensive genomic features were extracted from the mouse genome (mm39), filtering for protein-coding genes on standard chromosomes. For each gene, we calculated GC content, as well as number and length of introns, exons, coding sequences (CDS), and 5’ and 3’ untranslated regions (UTRs). We calculated tRNA Adaptation Index (tAI) using cortical neuron tRNA abundance data from previously published studies,^104^ incorporating relative adaptiveness values normalized by amino acid-specific maximum tRNA abundance. RBP binding sites were obtained from publicly available datasets organized in the CLIPdb database,^105^ filtering for brain/cortex experiments with confidence scores >3. Coordinates were lifted from mm10 to mm39 using the UCSC chain file. MicroRNA binding sites were extracted from TargetScan predictions, filtering for miRNAs expressed in developing dopaminergic neurons^106^ and filtering for targetscan score <80. Both RBP and miRNA binding site counts were normalized first by gene/UTR length to remove gene length bias. Our RNA immunoprecipitation (RIP-seq) data for neuronal RNA-binding proteins (FUS, HuD, HuR, CELF4) were integrated by defining targets as genes with an FDR < 0.05 and a fold change > 2 of IP over Input regressing out the effect of KCl treatment in binding and subtracting genes that were also enriched in the IgG IP over Input. The category of RBP interactors were then incorporated in the model as one-hot encoders. Poly(A) tail length differences were incorporated from nanopore sequencing analysis. To avoid multicollinearity we evaluated the multiple correlation of the features and decided to keep for downstream analysis the most relevant features for our purposes (identify features relevant for regulation of cytoplasmic RNA stability in response to KCl): for GC% features we kept only coding sequence and UTR quantifications (which correlate as expected with exonic and intronic GC content). For HuD and HuR RNA binding (R^2^ > 0.9), we decided to focus on HuD binding as HuD is neuronal enriched. Features spanning more than three orders of magnitude were log_10_-transformed.

##### Model training and testing

Genes were classified as activity regulated based on DESeq2 analysis and the model was ran for activity regulated genes alone. The preprocessed dataset was randomly split into a training set (90%) and a held-out test set (10%) using the train_test_split function from scikit-learn, with a fixed random state (random_state=20) to ensure reproducibility. Only the training set was used for hyperparameter tuning. All numerical features were scaled using the StandardScaler function from scikit-learn to ensure that the regularization penalty was applied equally across features. A LASSO linear regression model was implemented. The regularization strength (alpha) was optimized via grid search (GridSearchCV) with 10-fold cross-validation on the training data. The search space for alpha spanned eight orders of magnitude (from 0.00001 to 100). Cross validation coefficient of determination (R²) was used for optimization. The best-performing model from this search was selected for final evaluation.

#### HuD knockdown and TimeLapse-seq

TimeLapse-seq libraries from HuD knockdown and control neurons were processed from raw FASTQs with fastq2EZBakR (https://github.com/isaacvock/fastq2EZbakR) using default parameters. Samples were excluded if HuD knockdown efficiency was <50%, assessed by normalized HuD read counts relative to negative-control shRNA. After PCA analysis, non-targeting shRNA samples were analyzed together with no treatment shRNA samples. To identify HuD-sensitive genes, we used DESeq2 with a likelihood ratio test (LRT) across all time points, fitting the full model (∼ shRNA + Time) and testing against the reduced model (∼ Time). Genes with FDR (Benjamini–Hochberg) P_adj_ < 0.05 were considered HuD-sensitive. To obtain direction and time-specific effects, DESeq2 was run a second time by rerunning separate models at each time point (∼ shRNA), from which log2(fold change - HuD KD vs control) and Wald test statistics were extracted.

Enrichment of HuD RIP-seq targets among HuD-regulated genes was evaluated with the R package fgsea (Korotkevich, 2021, https://doi.org/10.1101/060012), ranking genes by log_2_(fold change) in HuD knockdown across all time points of KCl. To assess the potential for direct binding, we extracted the longest 3′UTR isoform per gene and scanned for predicted HuD recognition motifs using RBPmap.^90^

RNA half-lives were estimated using EZbakR.^56^ T-to-C mutation tables from fastq2EZBakR were used as input, with samples grouped by experimental condition (HuD knockdown vs control) and analyzed separately for each time of KCl treatment, assuming pseudo steady-state between conditions within the same time point. To correct for unequal levels of s^4^U incorporation between conditions, counts were normalized with NormalizeForDropout. Kinetic parameters of RNA degradation (*k_deg_*) and synthesis (*k_syn_*) were then estimated with EstimateKinetics, regularized across replicates using AverageAndRegularize, and compared between conditions using CompareParameters. Transcripts with P_adj_ < 0.1 and log_2_(FC k_deg_) > 0 were classified as destabilized, while those with log_2_(FC k_deg_) < 0 were classified as stabilized. Degradation rates were converted into mRNA half-lives (ln(2)/k_deg_) for display purposes.

#### Image quantification

Images were analyzed using ImageJ-Fiji (v1.5.4) first by background subtraction (rolling ball = 50), maximally Z-projecting the stack, and applying a 0.5 Gaussian filter. Manual somatic outlines were drawn and used to define cell edges for further analysis. DAPI masks were obtained through automatic Otsu thresholding with two rounds of dilation and hole filling to retain nuclear shape. HuD masks were obtained through a threshold of between 22,000-65,535 pixel intensity which was kept consistent through all images to lower background staining. Statistical analysis of image quantification was done in Graphpad Prism (v10) software after applying a 1% ROUT outlier test.

#### AlphaFold predictions

To investigate potential structural interfaces between HuD and its binding partners, we used AlphaFold3 to generate pairwise co-folded models of HuD with each interactor identified by IP–mass spectrometry. For each HuD–partner pair, five independent models were generated and both the CIF structural coordinates and the corresponding full-data JSON outputs were analyzed in R. Interaction interfaces were defined by parsing the predicted aligned error (PAE) matrices and contact probability scores for each model. Residues from HuD (chain A) and the interactor protein (chain B) were mapped to their chain-specific indices, and interaction domains (PAE <15 Å) were extracted and annotated with high-confidence contact residues (contact probability >0.5 and PAE <8 Å).

