## Supplementary_figures for "Neuronal activity triggers widespread changes in RNA stability"

### Supplementary Figure 1

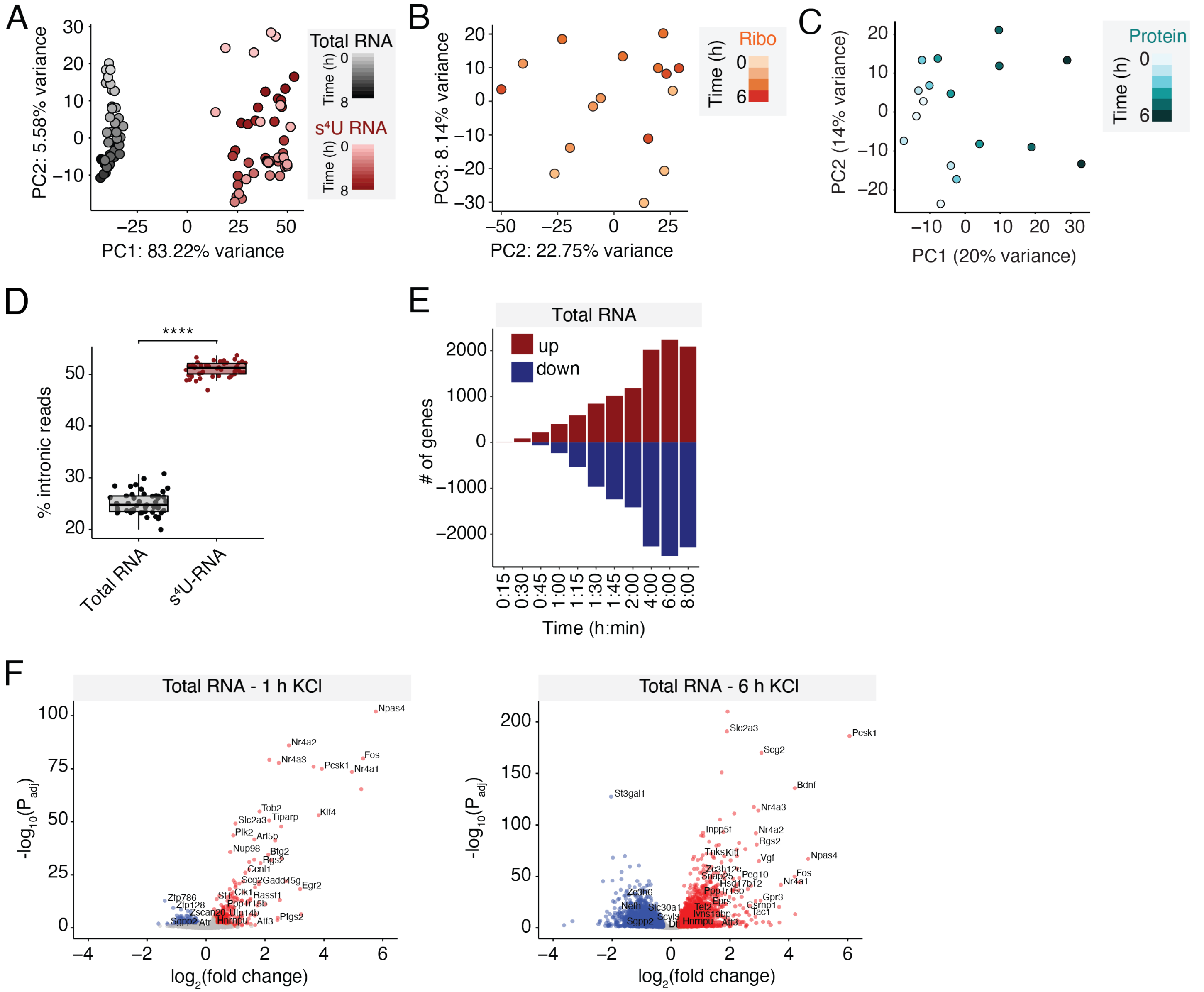


**Supplementary Figure 1: RNA metabolic labeling quality metrics, Related to Figure 1.**
(**A**) PCA analysis on total RNA and s$^4$U RNA for the top 2000 most variable genes. (**B**) PCA analysis on Ribo-seq RNA for the top 2000 most variable genes. (**C**) PCA analysis on whole-cell proteomics for the top 500 most variable proteins. (**D**) Boxplot of the average percentage of reads mapped to introns across all gene-mapped reads per sample for total RNA and s$^4$U-RNA samples. Newly transcribed RNAs are expected to be enriched for unspliced pre-mRNA relative to total RNA. **** $p < 2.2e\text{-}16$, exact two-sample Kolmogorov-Smirnov test. (**E**) Bar plot of significant (DESeq2 $P_{adj} < 0.05$, $|FC| > 1.2$) differentially expressed genes by total RNA-seq at various time points following membrane depolarization. Upregulated and downregulated genes are plotted as positive and negative values on the y-axis, respectively. (**F**) Volcano plots of $-\log_{10}(P\text{adj})$ versus $\log_2(\text{FC})$ in total RNA gene expression at 1 h (top) and 6 h (bottom) versus 0 h KCl.

Supplementary Figure 2

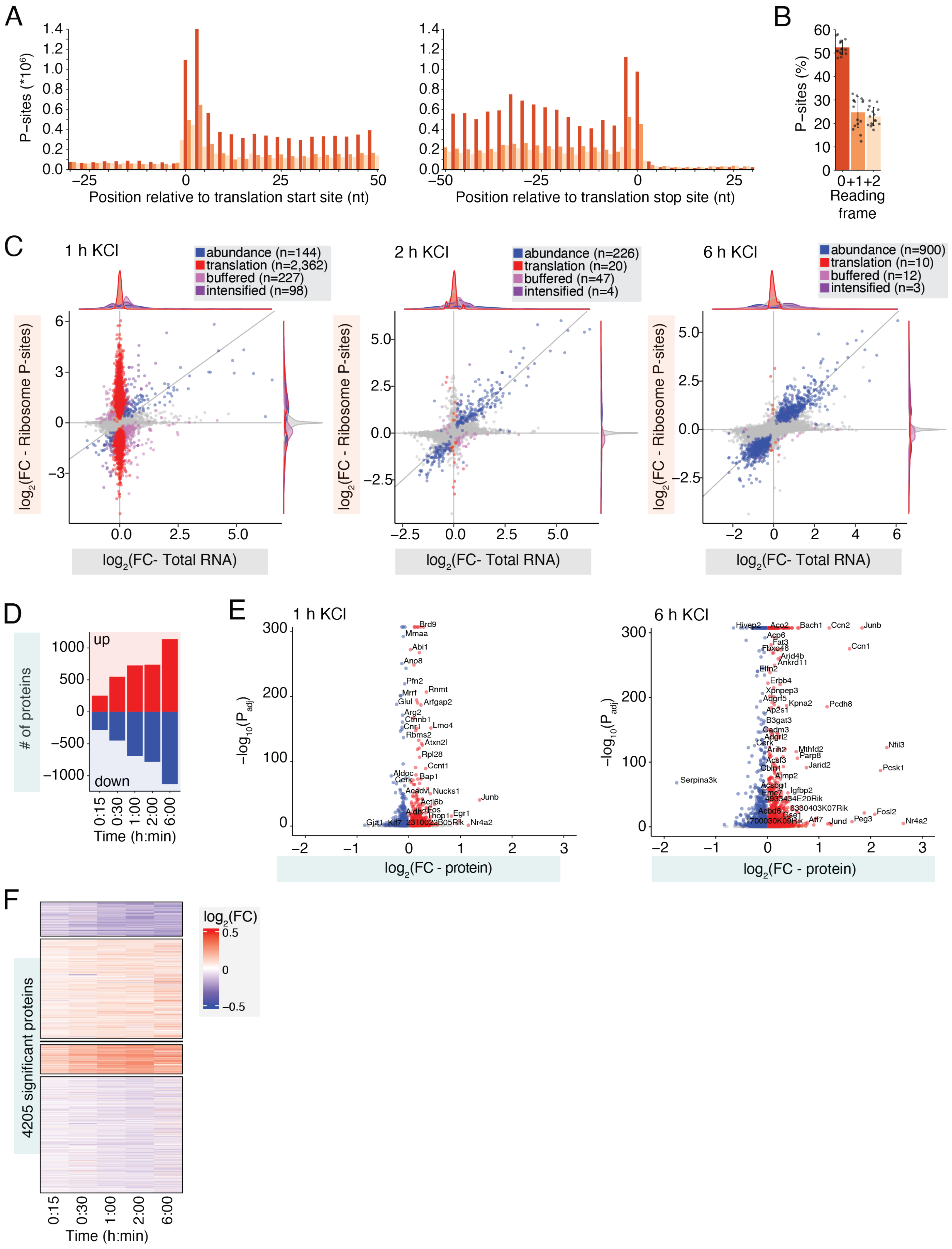

**Supplementary Figure 2: Ribo-seq and whole-cell proteomics quality metrics, Related to Figure 1.**

(**A**) Bar plot displaying P-sites derived from offset-corrected Ribo-seq reads in the first 50 nt of annotated ORFs relative to the translation start site (left) and in the last 50 nt of annotated ORFs relative to the translation stop site (right). (**B**) Bar plot of the percentage of footprints in each reading frame. Data are shown as mean ± s.d.; $n = 4$ biologically independent primary neuron cultures, 4 time points per biological replicate. (**C**) Scatter plot of $\log_2$(FC) in Ribo-seq (y-axis) versus RNA-seq (x-axis) between stimulated (1, 2, and 6 h KCl) and unstimulated primary neurons. Abundance-regulated genes (blue; change in RNA abundance with no change in ribosome density (RD)); translationally-regulated genes (red; change in RD with no change in transcription); buffered genes (light purple; change in RD that counterbalances the change in mRNA transcription); and intensified genes (dark purple; change in RD that amplifies the change in mRNA) are highlighted. (**D**) Bar plot of significant ($P_{adj} < 0.05$) differentially expressed proteins by TMT-MS proteomics at various time points following membrane depolarization. Upregulated and downregulated genes are plotted as positive and negative values on the y-axis, respectively. (**E**) Volcano plots of −log10(Padj) versus log2(FC in protein abundance) changes at 2 h (left) and 6 h (right) versus 0 h KCl. (**F**) Heatmap of $\log_2$(FC) in protein abundance for all proteins with a significant activity-dependent change in protein abundance in at least one time point relative to unstimulated neurons.

### Supplementary Figure 3

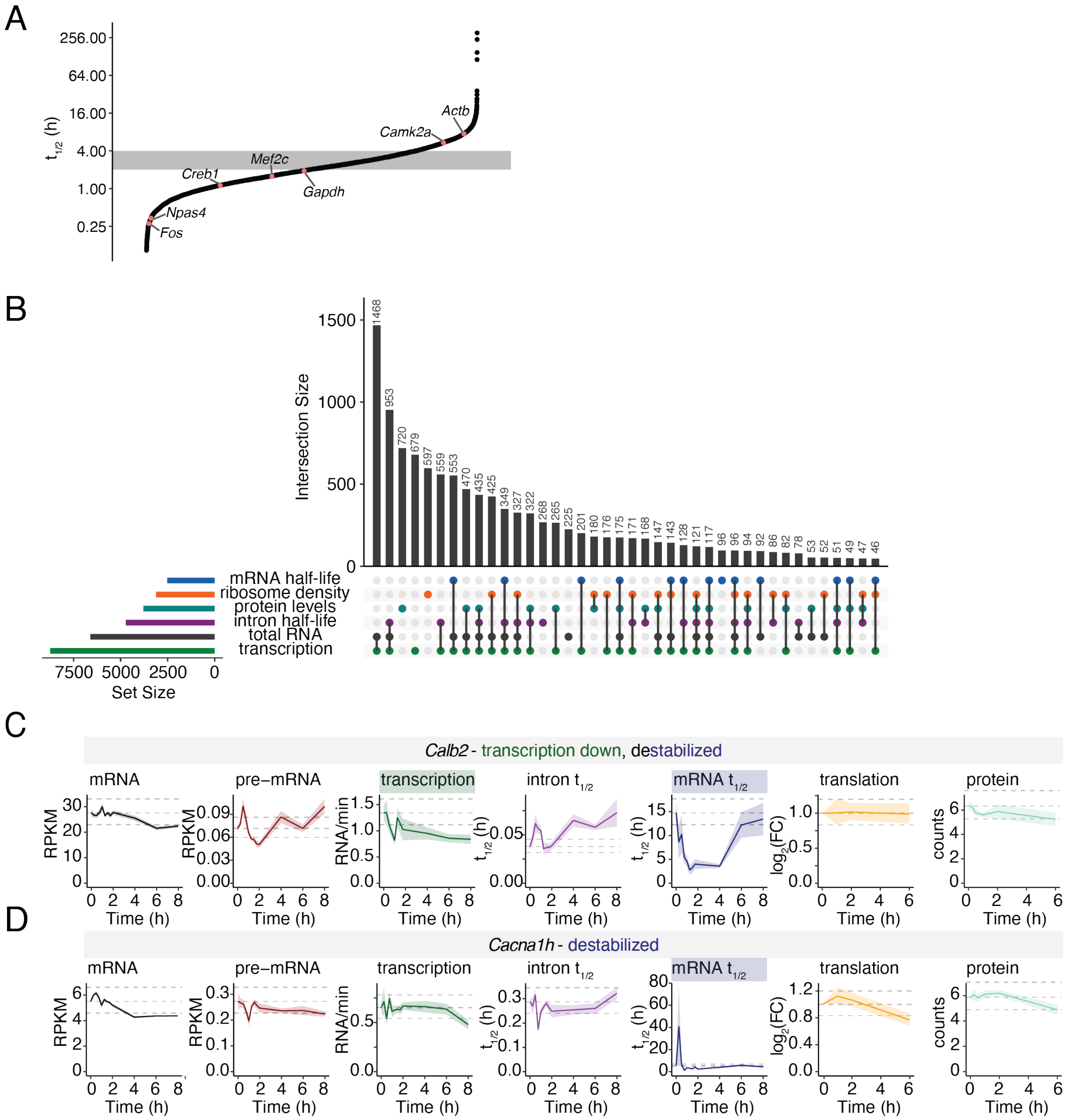


**Supplementary Figure 3: Measuring Activity-Dependent RNA Dynamics, Related to Figure 1.**
(**A**) Distribution of half-lives for all RNAs in unstimulated neurons. ERG transcription factors (*Fos*, *Npas4*) have shorter half-lives compared to constitutively expressed transcription factors (*Srf*, *Mef2c*), while housekeeping genes (*Gapdh*) and genes trafficked to dendrites (*Camk2a*) have longer than average half-lives. Shading indicates average half-life range of 2-4 h. (**B**) Upset plot of numbers of genes exhibiting significant activity-dependent change in total RNA, pre-mRNA, synthesis, processing, degradation, Ribo-seq reads, ribosome density, or protein abundance. (**C&D**) Line plots of synthesis, processing, degradation, and translation, as well as total RNA, pre-mRNA, and protein abundance, for select activity-destabilized genes. Dark line = mean, shading = ± SEM, dashed

lines = mean in unstimulated neurons ±1.2-FC. Notably, gain-of-function mutations within the activity-destabilized gene *Cacna1h* are associated with epilepsy and febrile seizures,[86] suggesting that the activity-dependent decrease in CACNA1H protein levels driven by increased mRNA turnover may be important for regulating spontaneous neuronal firing during postnatal development.

Supplementary Figure 4

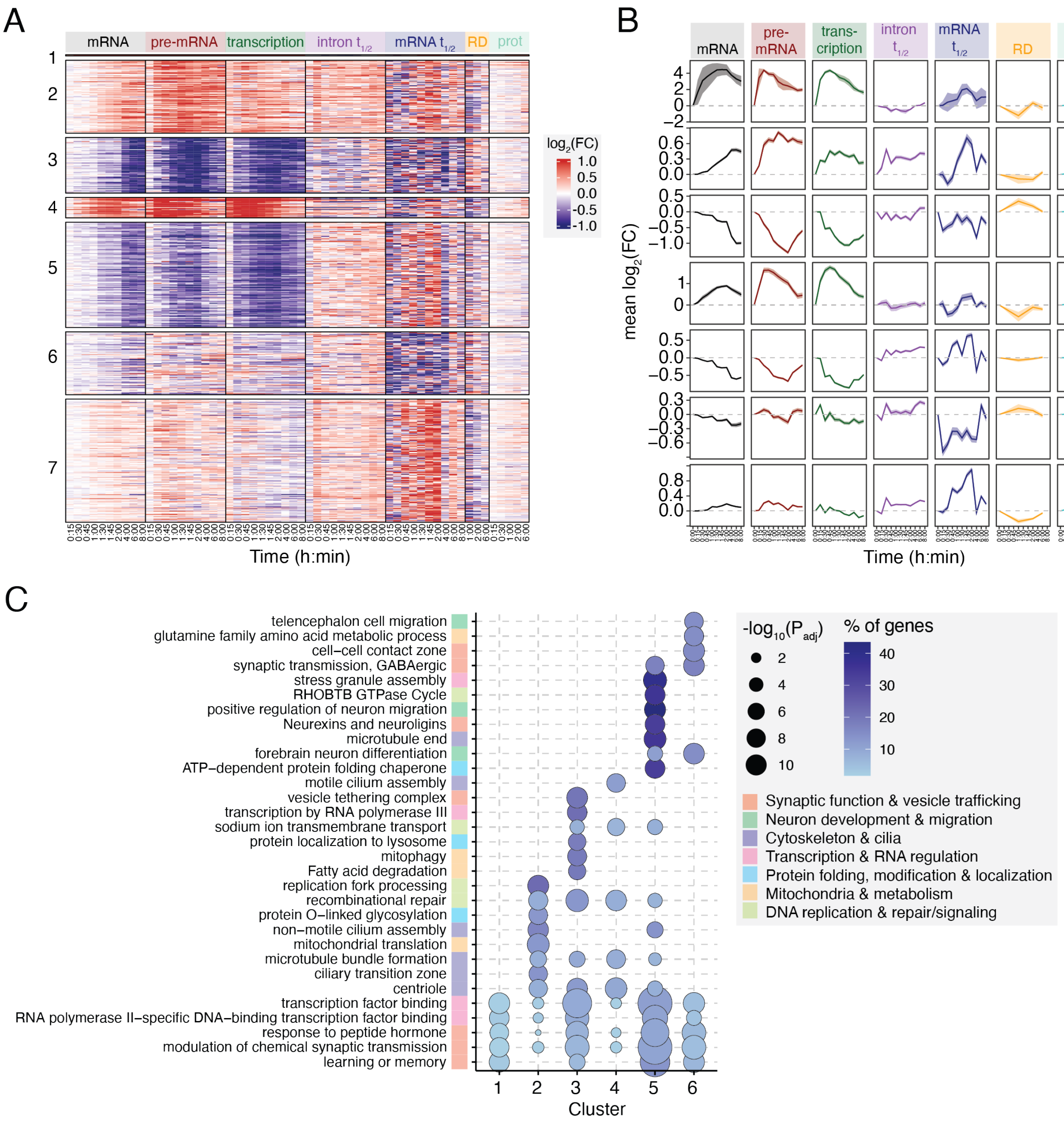


**Supplementary Figure 4: Activity-dependent modulation of RNA stability is widespread and impacts protein abundance, Related to Figure 1.**

(**A**) Heatmap of $\log_2$(FC) in rates of RNA synthesis, processing, and degradation, as well as total RNA, pre-mRNA, ribosome density (RD), and protein abundance, for all activity-regulated genes with detectable protein levels. (**B**) Metaplots of mean fold change in all rates, separated by clusters defined in A. (**C**) Dot plot of select top GO terms (FDR < 0.05) in each cluster defined in Fig. 1E.

### Supplementary Figure 5

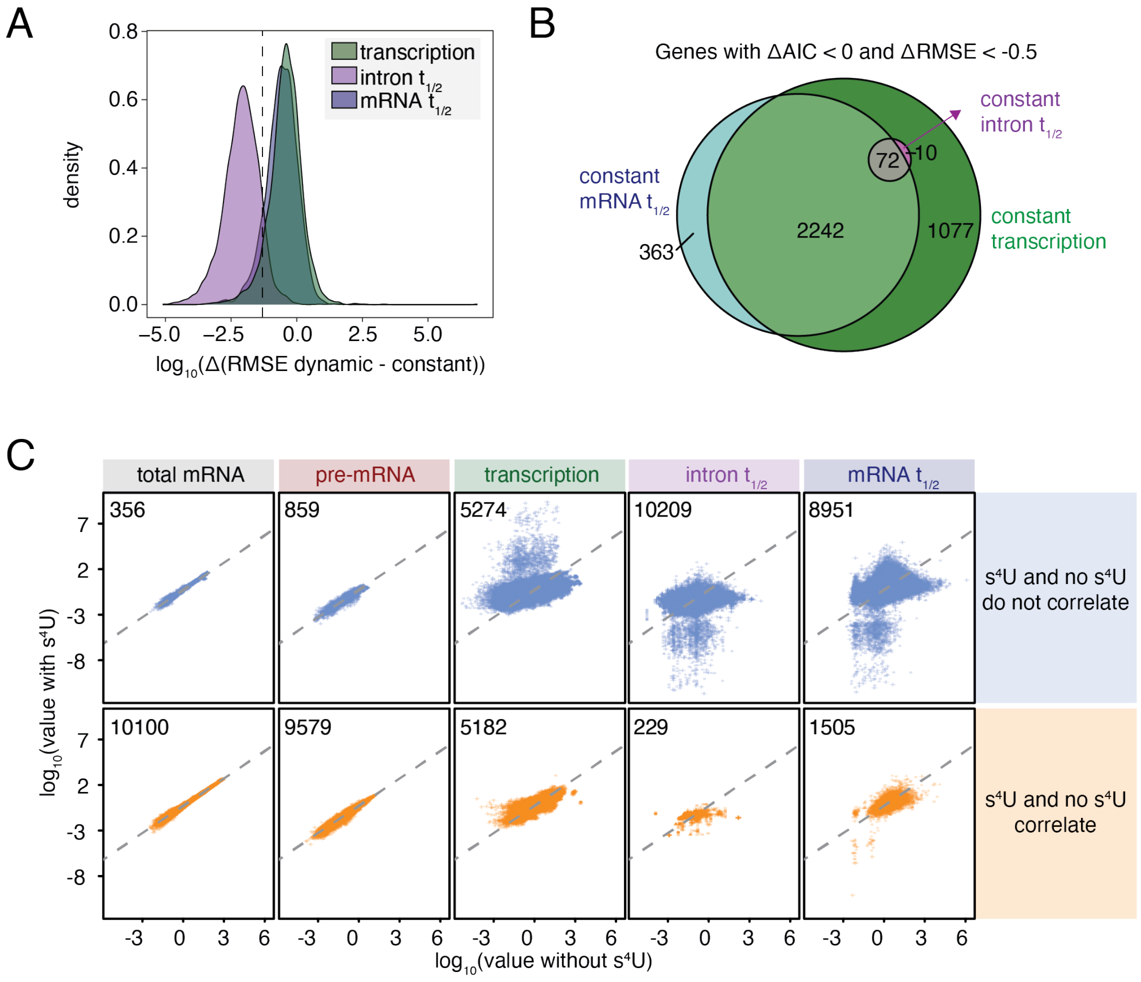


**Supplementary Figure 5: Activity-dependent modulation of RNA stability is widespread and a key determinant of RNA and protein output, Related to Figure 2.**

(**A**) Distribution of ΔRMSE(dynamic - constant) models for each transcript for models of constant transcription (green), intron $t_{1/2}$ (purple) and mRNA $t_{1/2}$ (blue). (**B**) Euler diagram showing overlap of genes requiring dynamic regulation of transcription (green), mRNA half-life (blue), or intron half-life (violet) to explain activity-dependent changes in total RNA levels. Genes were identified using dual criteria: Akaike information criterion (AIC) favoring the dynamic model and ΔRMSE < -0.5 between dynamic and constant-rate models. Numbers indicate gene counts in each category. (**C**) Validation of computational RNA kinetic rate inference against metabolic labeling in primary neurons. Scatterplots compare estimates obtained without metabolic labeling (x-axis; total RNA-seq with INSPEcT-) versus with metabolic labeling (y-axis; $s^4$U-seq with INSPEcT). Values shown are RPKM for total mRNA and pre-mRNA, RNA/min for transcription rate, and half-life ($t_{1/2}$) for intronic and mRNA decay. Top panels: transcripts with poor correlation between the two approaches ($R^2 < 0.6$). Bottom panels: transcripts with strong correlation ($R^2 > 0.6$). Numbers in each panel indicate the number of transcripts plotted.

### Supplementary Figure 6

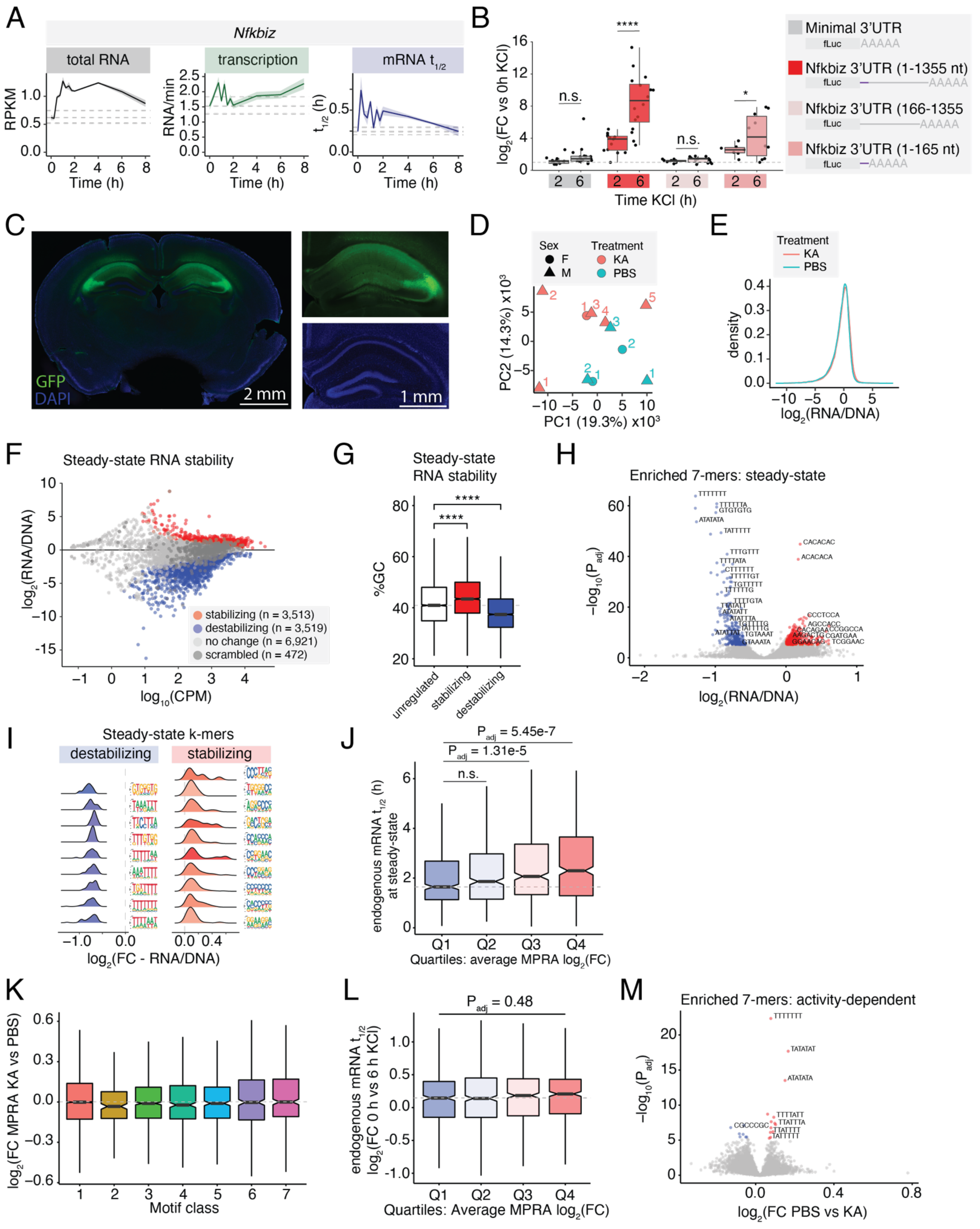

**Supplementary Figure 6: Modular 3'UTR RNA sequences are sufficient to drive activity-dependent changes in RNA stability *in vivo,* Related to Figure 3.**

(**A**) Line plots of synthesis and degradation rates, as well as total RNA abundance, for *Nfkbiz*. Dark line = mean, shading = ±SEM, dashed lines = mean in unstimulated neurons ± 1.2-FC. (**B**) Box plot of $log_2$(FC) in normalized firefly luciferase abundance relative to unstimulated neurons for the luciferase minimal 3'UTR, *Nfkbiz* full 3'UTR (1-1355 nt), a truncated *Nfkbiz* 3'UTR (166-1355 nt), and the SE of the *Nfkbiz* 3'UTR (1-165 nt). *n*=8 independent biological replicates. One-way ANOVA with Tukey's HSD post-hoc test; $P_{adj} < 0.05$ (*), $P_{adj} = 6.8e-14$ (****). (**C**) Immunofluorescence images of virally injected MPRA AAVs expressing EGFP in the mouse hippocampus. Scale bar = 2 mm (left) and 1 mm (right zoom). (**D**) PCA plot of all samples separated by treatment (6 h KA or PBS) and sex. One sample (KA-treated male mouse, replicate 1) was excluded from further analysis because KA treatment did not induce neuronal activity. (**E**) Distribution of $log_2$ RNA abundance (depth-normalized reads) normalized to corresponding viral DNA for each element in the AAV library colored by experimental condition (6 h KA or PBS). (**F**) MA plot of $log_{10}$(counts per million reads (CPM)) versus $log_2$(FC) between RNA abundance and corresponding viral DNA in PBS-injected mice. 3'UTR elements were defined as stabilizing or destabilizing based on DESeq2 $P_{adj} < 0.05$ and $log_2(FC) > 0$ (stabilizing) or < 0 (destabilizing). Dark gray indicates scrambled controls. (**G**) Box plot of %GC content of stabilizing and destabilizing 3'UTR elements in unstimulated neurons. KS test with Dunn's test for multiple hypotheses. **** $P_{adj} < 2.2e-16$. (**H**) Volcano plot of $log_2$(FC) versus -$log_{10}(P_{adj})$ of all 7-mers within stabilizing and destabilizing 3'UTR elements in unstimulated neurons. Red = stabilizing 7-mers, blue = destabilizing 7-mers, light gray = $P_{adj} > 0.05$. (**I**) Distribution plot of mean $log_2$(FC) in reporter expression for MPRA library members containing individual 7-mers, grouped into clusters of similar sequences as defined in Fig. S3I. Each distribution represents the set of 7-mer means within a cluster; sequence logos show the position weight matrix (PWM) for each cluster. (**J**) Boxplots of steady-state RNA half-lives for transcripts binned by average MPRA motif activity at steady-state. MPRA motifs were grouped by their endogenous 3′UTR of origin and averaged to generate a per-transcript motif activity score ($log_2$(RNA/DNA)). Transcripts were then ranked and binned into deciles by this score, with Q1 representing the most destabilizing motif activity and Q4 the most stabilizing. Boxplots represent the median (center line), interquartile range (box), and 1.5× IQR (whiskers); notches indicate a 95% confidence interval around the median. Statistical significance was assessed by Kruskal–Wallis test with Dunn's post hoc pairwise comparisons. (**K**) Boxplots of $log_2$(FC) in MPRA expression, in which motifs were binned by class. Class 1 - Conserved elements (PhyloP ≥ 0.5) from activity-stabilized or -destabilized 3′UTRs (*n*=11,522); Class 2: Fast-evolving elements (PhyloP ≤ -0.5) from activity-regulated 3′UTRs (*n*=244); Class 3: Non-conserved elements (−0.5 < PhyloP < 0.5) from activity-regulated 3′UTRs (*n*=1,234); Class 4: Conserved elements from genes not regulated by activity (*n*=500); Class 5: Non-conserved elements from non-regulated genes (*n*=500); Class 6: Conserved elements from transcription-only genes (*n*=500); Class 7: GC-matched scrambled sequences from class 1 (*n*=500). Importantly, scrambled controls were centered around $log_2(FC) = 0$, suggesting that reporter expression is not globally shifted in response to KA stimulation. (**L**) Boxplots of $log_2$(FC) in endogenous mRNA half-life between 0 and 6 h membrane depolarization for transcripts binned by average MPRA motif $log_2$(FC) in stability in PBS vs 6 h KA. MPRA motifs were grouped by their endogenous 3′UTR of origin and averaged to generate a per-transcript motif activity score ($log_2$(FC RNA KA/PBS)). Transcripts were then ranked and binned into deciles by this score, with Q1 representing the most destabilizing motif activity and Q4 the most stabilizing. Boxplots represent the median (center line), interquartile range (box), and 1.5× IQR (whiskers); notches indicate a 95% confidence interval around the median. Statistical significance was assessed by Kruskal–Wallis test with Dunn's post hoc pairwise comparisons. When we aggregated motif effects across the MPRA by endogenous transcript, these summed stability scores showed only a weak, non-significant correspondence with activity-dependent changes in endogenous transcript half-lives, consistent either with limited sampling or with a more modular mode of regulation in the activity-dependent context. (**M**) Volcano plot of $log_2$(FC) versus -$log_{10}(P_{adj})$ of all 7-mers within activity-stabilizing and -destabilizing 3'UTR elements. Red = activity-stabilizing 7-mers, blue = activity-destabilizing 7-mers, light gray = $P_{adj} > 0.05$.

### Supplementary Figure 7

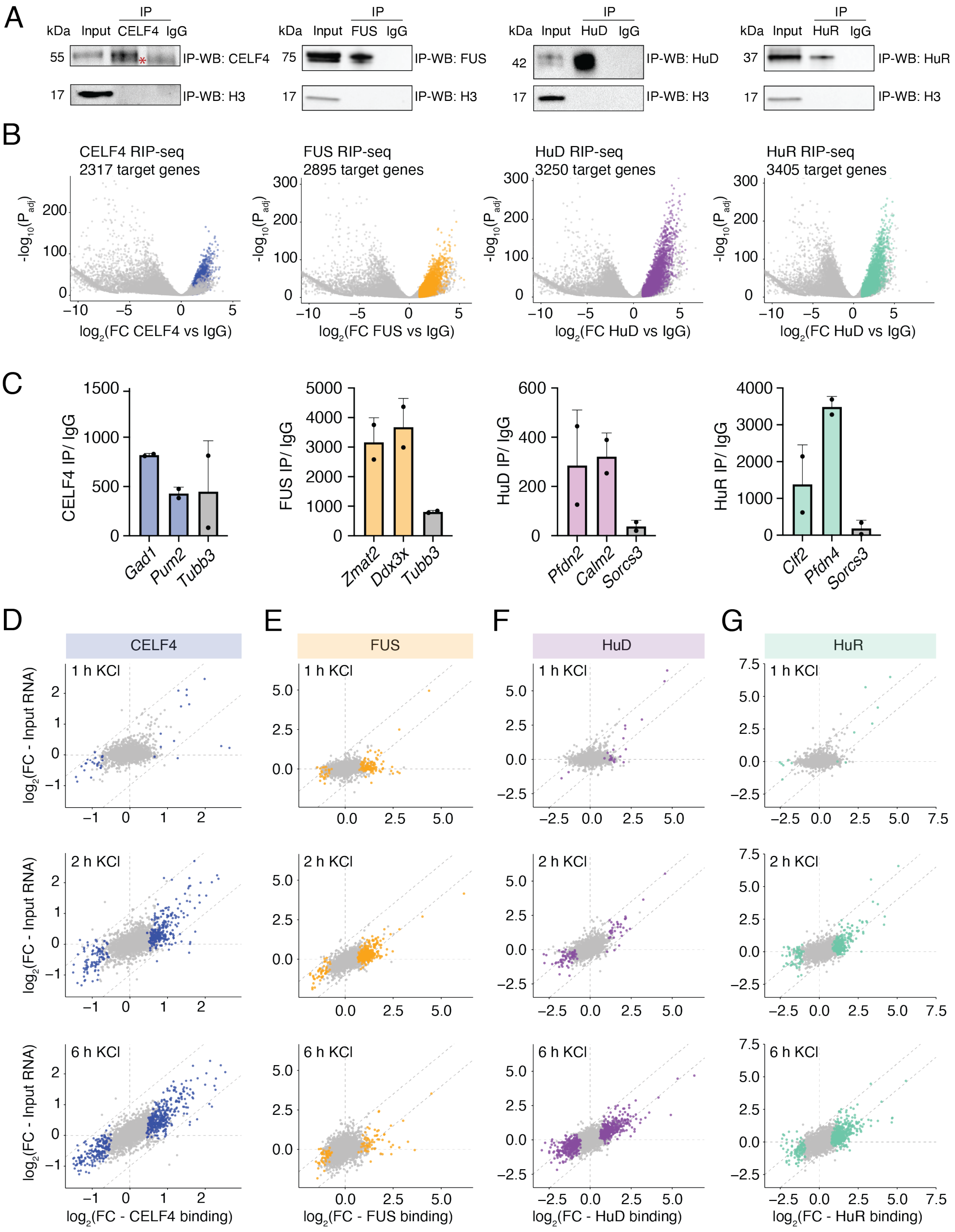

**Supplementary Figure 7: RIP-seq QC, Related to Figure 4.**

(**A**) Representative western blot analysis of immunoprecipitations performed in primary neurons for HuD, HuR, CELF4, and FUS versus an IgG control. (**B**) Volcano plot of $-\log_{10}(P_{adj})$ versus $\log_2(FC)$ of RNAs enriched by each RBP RIP-seq vs total RNA abundance. (**C**) RIP-qPCR validation of RBP targets (CELF4 = blue, FUS = orange, HuD = purple, HuR = teal) vs non-targets (gray), *n*=2 independent biological replicates, bars represent mean ± SEM. (**D-G**) Scatterplot of $\log_2(FC)$ in RBP binding for each RBP (D = CELF4, E = FUS, F = HuD, G = HuR) versus $\log_2(FC)$ in total RNA abundance at 1, 2, and 6 h KCl versus 0 h.

### Supplementary Figure 8

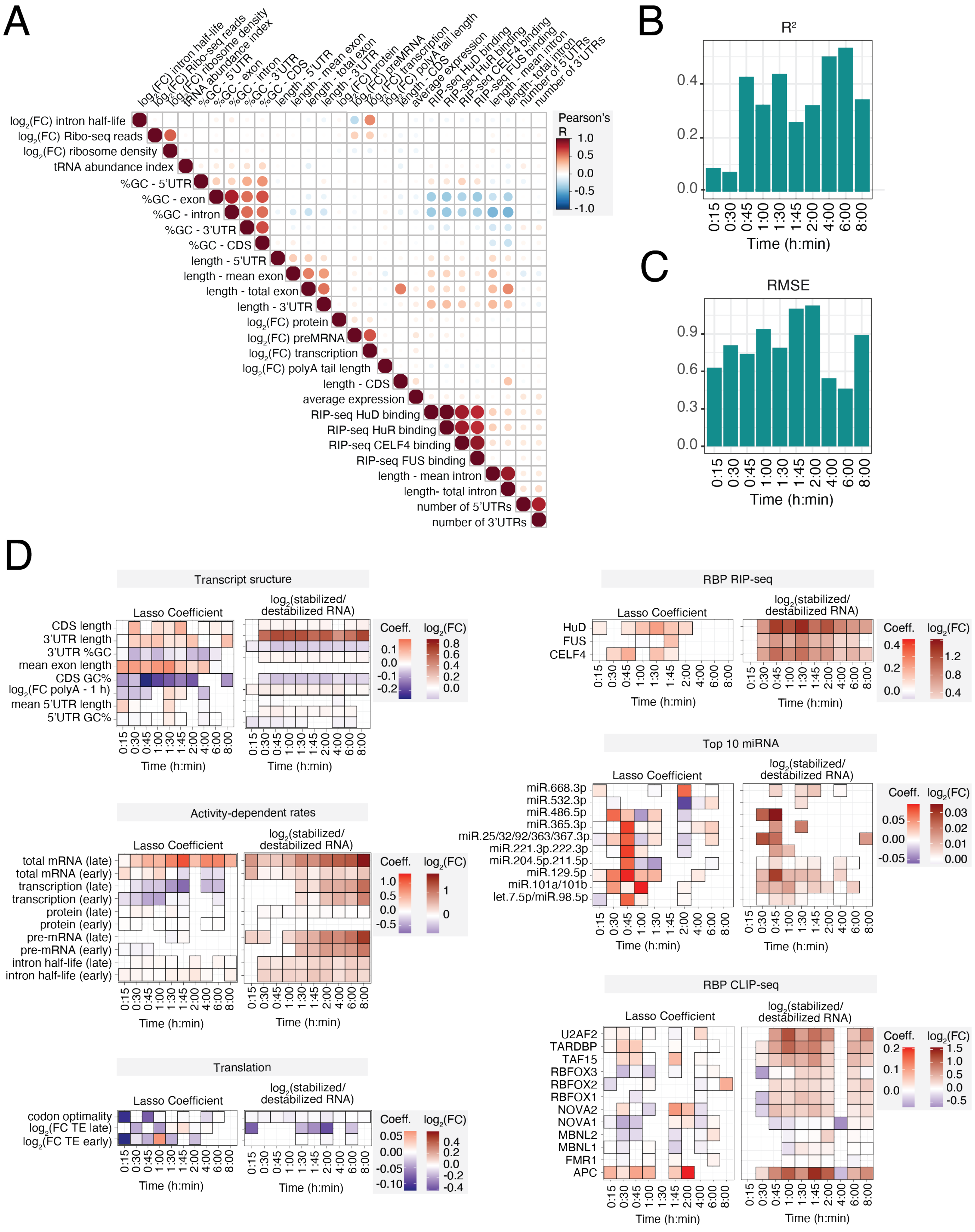

**Supplementary Figure 8: LASSO machine learning QC, Related to Figure 4.**

(**A)** Pearson's R correlation matrix of features included in the LASSO machine learning model. A representative feature from each correlated cluster was selected for input into the model to avoid confounding effects of collinearity. (**B&C**) Bar plot of $R^2$ (C) and root mean square error (RMSE) (D) for LASSO model at each time point of membrane depolarization. (**D**) Heatmap of LASSO coefficient and mean $\log_2$(FC) between activity-stabilized and -destabilized transcripts for each feature in the LASSO model at each time point of membrane depolarization. Box outline indicates significant predictive power (LASSO coefficient) or $P_{adj} < 0.05$ by Kruskal-Wallis test. Color scale represents $\log_2$(FC stabilized/destabilized genes at each time of KCl)).

### Supplementary Figure 9

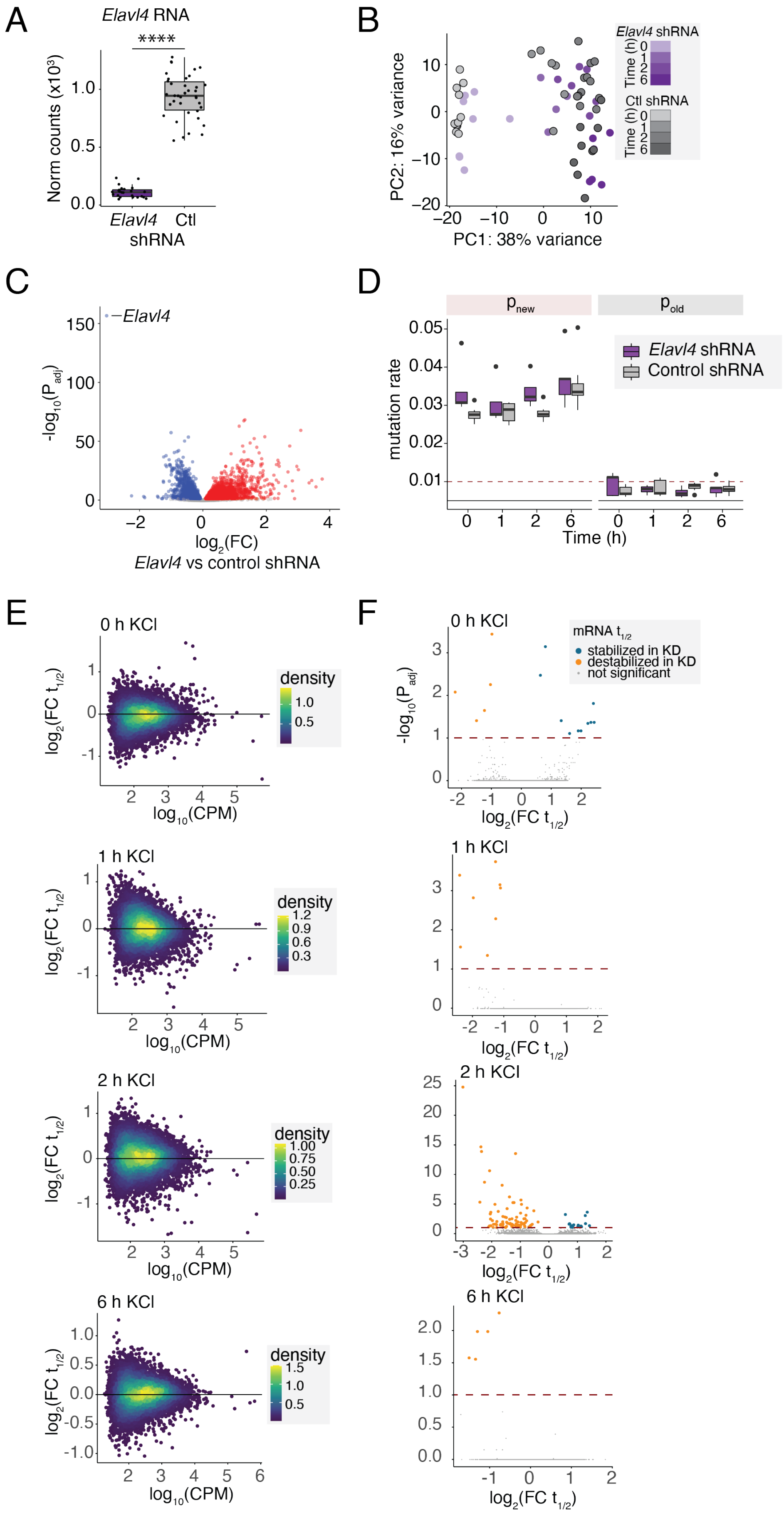

**Supplementary Figure 9: HuD regulates activity-dependent RNA stability, Related to Figure 5.**

(**A**) Boxplot of depth-normalized counts of *Elavl4* (the gene that encodes HuD) in HuD KD in primary cortical neurons with shRNA versus non-targeting control. *n*=4-6 independent biological replicates per shRNA across 4 time points (0, 1, 2, and 6 h membrane depolarization). DESeq2 $P_{adj}$ < 1e-16 (****). (**B**) PCA plot of RNA-seq from HuD KD versus control, colored by time of membrane depolarization. (**C**) Volcano plot of $-\log_{10}(P_{adj})$ versus $\log_2(FC)$ in RNA expression between HuD KD and control across all time points of membrane depolarization. Red = upregulated genes ($P_{adj} < 0.05$, $\log_2(FC) > 0$) (dark red = HuD-bound mRNA, light red = non-target); Blue = downregulated genes ($P_{adj} < 0.05$, $\log_2(FC) < 0$) (dark blue = HuD-bound mRNA, light blue = non-target), Gray = $P_{adj} > 0.05$ (dark gray = HuD-bound mRNA, light gray = non-target). (**D**) Boxplots of average U-to-C mutation rates in TimeLapse-seq data for RNA fractions classified as newly transcribed ($P_{new}$) or pre-existing ($P_{old}$). Primary cultured neurons were metabolically labeled with $s^4U$ at different time points following membrane depolarization, and U-to-C mutations were detected via TimeLapse-seq as a proxy for newly synthesized RNA. Mutation rates were quantified separately for $P_{new}$ and $P_{old}$ fractions using the EZbakR pipeline. Each point represents a biological replicate, grouped by condition (HuD KD or control) and time after stimulation. Gray and red dotted lines indicate expected mutation rate boundaries for $P_{old}$ (low) and $P_{new}$ (high) fractions, respectively. Boxplots show the median (center line), interquartile range (box), and 1.5× IQR (whiskers). (**E**) Scatterplot of $\log_{10}$(Counts Per Million (CPM)) reads versus $\log_2(FC)$ in mRNA half-life ($t_{1/2}$) between HuD KD and control at 0, 1, 2, and 6 h membrane depolarization. (**F**) Volcano plot of $-\log_{10}(P_{adj})$ versus $\log_2(FC)$ in mRNA $t_{1/2}$ between HuD KD and control at 0, 1, 2, and 6 h membrane depolarization. Red line indicates $P_{adj} = 0.1$. Orange = destabilized in HuD KD ($P_{adj} < 0.1$, $\log_2(FC\ t_{1/2}) < 0$); Blue = stabilized in HuD KD ($P_{adj} < 0.1$, $\log_2(FC\ t_{1/2}) > 0$), Gray = $P_{adj} > 0.1$.

### Supplementary Figure 10

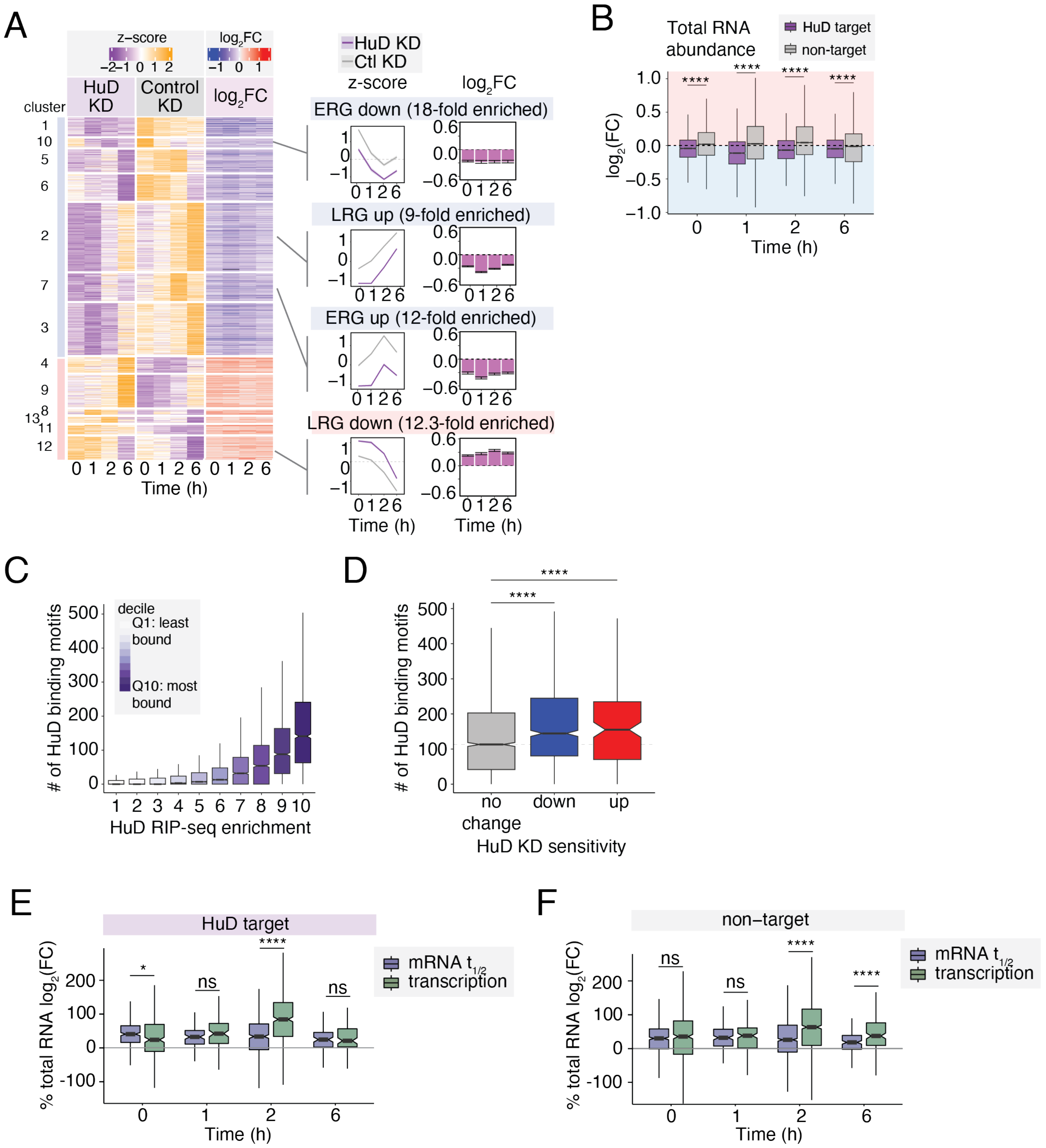


**Supplementary Figure 10: HuD regulates activity-dependent RNA stability, Related to Figure 5.**

(**A**) Left - Heatmap of normalized counts in HuD KD and control neurons, as well as and log2(FC) in mRNA abundance between HuD KD and control, subsetting for HuD-sensitive genes, which are defined as HuD-bound mRNAs that are significantly (LRT Padj <0.05) misregulated upon HuD knockdown. Middle - select metaplots of clusters defined in left panel showing mean ± SEM of normalized counts in HuD KD versus control. Right - select metaplots of clusters defined in left panel showing mean ± SEM of log2(FC) in mRNA abundance in HuD

KD versus control. (**B**) Boxplot of the number of predicted HuD-binding sites per gene within the 3'UTR, where genes were divided into deciles based on HuD enrichment over Input by RIP-seq. (**C**) Boxplot of the number of predicted HuD-binding sites per gene, separated by genes that are upregulated, downregulated, or unchanged in RNA expression between HuD knockdown versus control as defined in Fig. S5.1C. (**D**) Boxplot of $log_2$(FC) in mRNA abundance between HuD KD and control at each time of membrane depolarization, comparing HuD-bound mRNAs versus non-targets. $P_{adj} < 2.2e\text{-}16$ ****, KS test with BH multiple hypothesis correction. Boxplots are shown as median ± IQR (whiskers = 1.5*IQR). (**E**) Box and whisker plot of the relative contribution of transcription and mRNA $t_{1/2}$ to total RNA fold changes in HuD KD versus control for HuD-bound mRNAs. Boxplots are shown as median ± IQR (whiskers = 1.5*IQR). $P_{adj} < 0.001$ (****); $P_{adj} < 0.05$ (*); $P_{adj} = 0.258$ (ns); KS test with BH multiple hypothesis correction. (**F**) Box and whisker plot of the relative contribution of transcription and mRNA $t_{1/2}$ to total RNA fold changes in HuD KD versus control for mRNAs that are not bound by HuD. Boxplots are shown as median ± IQR (whiskers = 1.5*IQR). $P_{adj} < 0.001$ (****); $P_{adj} < 0.05$ (*); $P_{adj} = 0.258$ (ns); KS test with BH multiple hypothesis correction.

Supplementary Figure 11

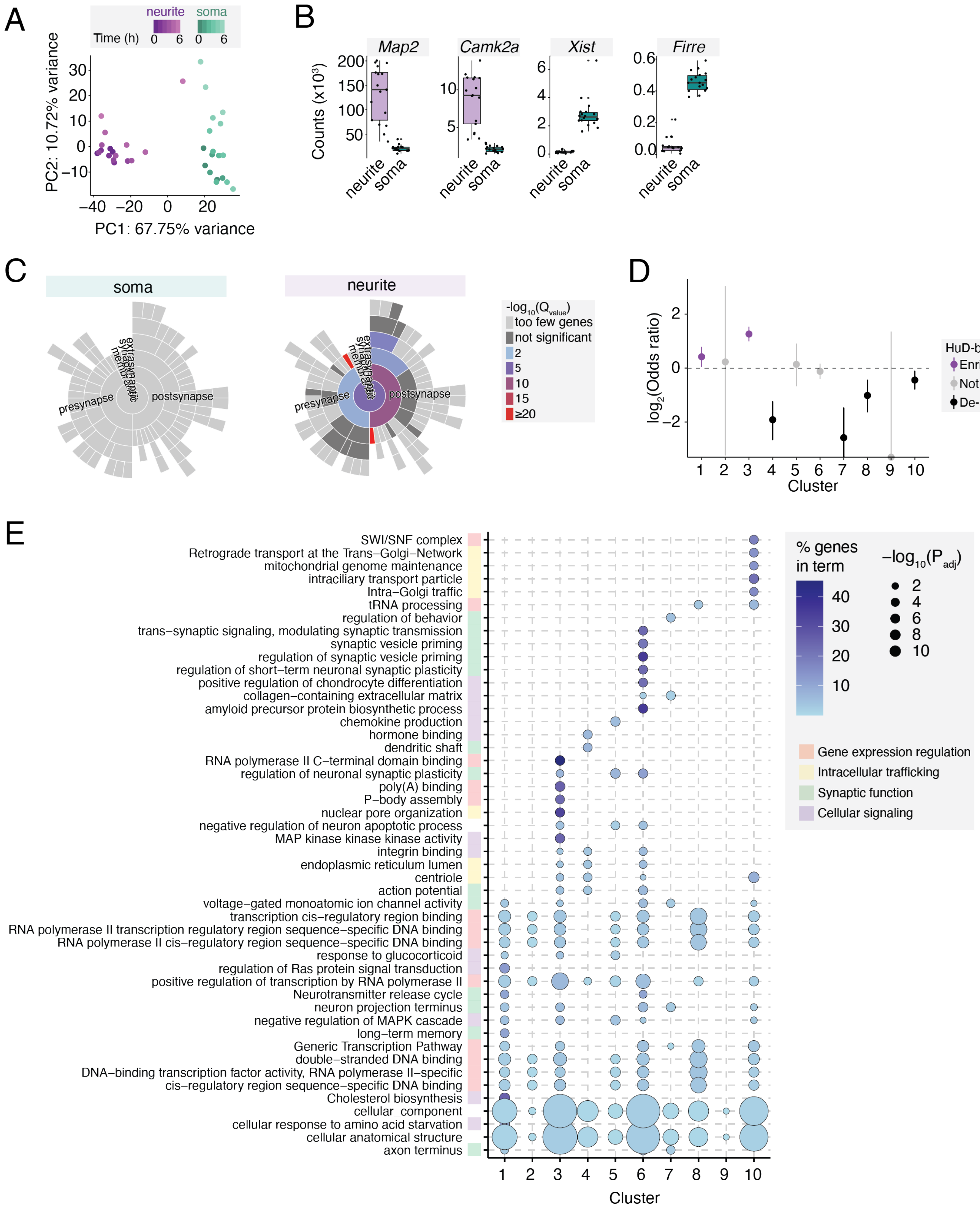


**Supplementary Figure 11: Separation chamber RNA-seq QC, Related to Figure 6.**

(**A**) PCA plot of all samples separated by neurites/soma in separation chambers and colored by time of KCl for the top 2000 most variable genes. (**B**) Boxplot of normalized counts for example soma- and neurite-enriched genes. Boxplots are shown as median ± IQR (whiskers = 1.5*IQR). (**C**) GO analysis of soma vs neurite-enriched transcripts using SynGO. (**D**) Odds ratio of HuD-bound mRNA enrichment across clusters defined in Fig. 6D.

Odds ratios ($\log_2$) with 95% confidence intervals are shown relative to the background gene set (dashed line = no enrichment). Points are colored by FDR significance (purple, $q < 0.05$ $\log_2$(odds ratio) > 0; black, $q < 0.05$ $\log_2$(odds ratio) < 0; gray, $q > 0.05$.). (**E**) Dot plot of the top GO terms (FDR < 0.05) in each cluster defined in Fig. 6D.

### Supplementary Figure 12

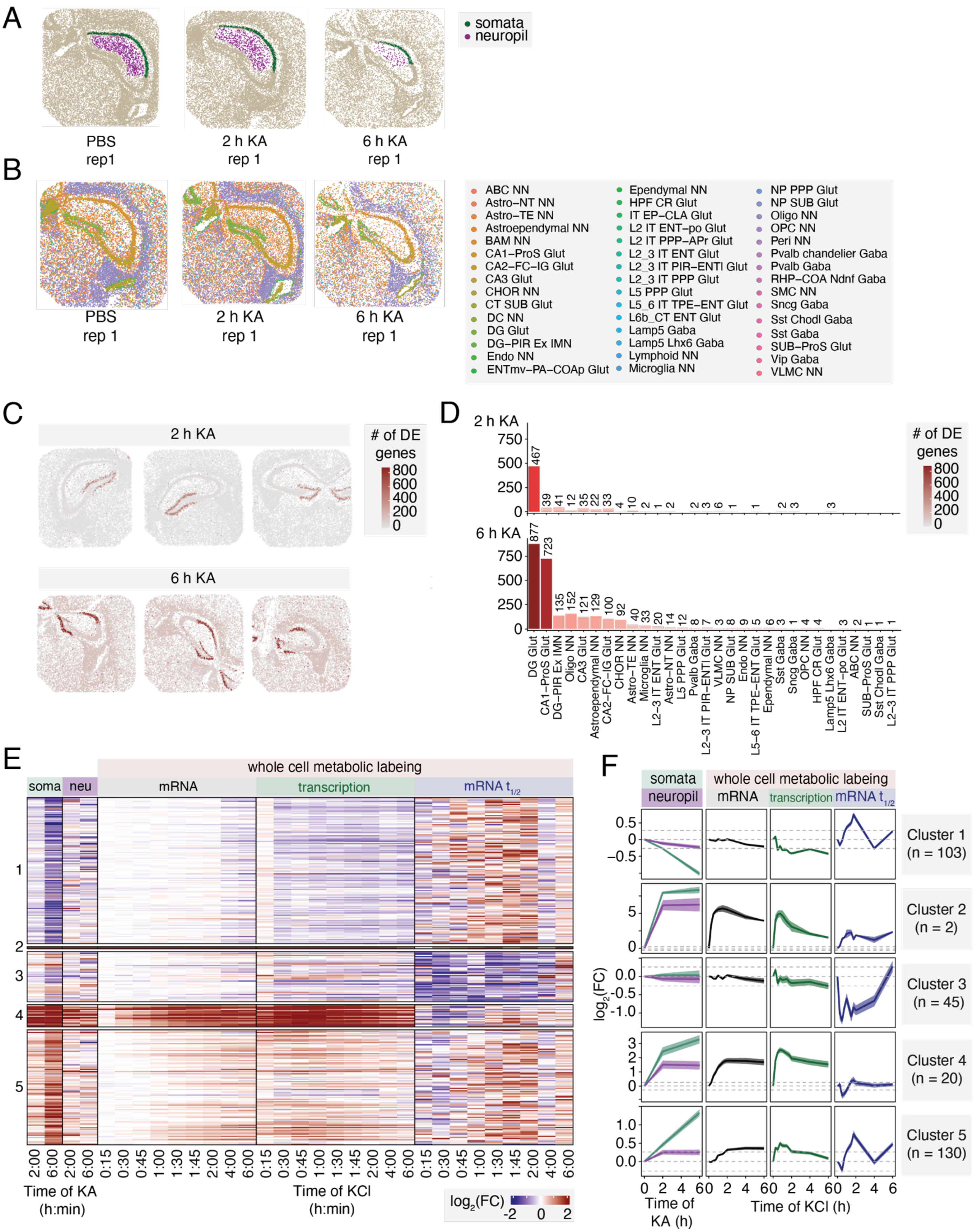

**Supplementary Figure 12: Slide-seqV2 QC, Related to Figure 6.**

(**A**) Spatial plot of beads defined as CA1 soma (green) and neuropil (purple). (**B**) Spatial visualization of Slide-seqV2 slices from PBS-, 2 h KA-, and 6 h KA-injected mice with cell sub-type information overlaid. $n$=143,394 beads, 8 mice. (**C**) Spatial plot of the number of differentially expressed genes by cell type relative to PBS controls in 2 and 6 h KA-injected samples. (**D**) Bar plot of the number of differentially expressed genes by cell type relative to PBS controls in 2 and 6 h KA-injected samples. (**E**) Heatmap of $\log_2$ (FC) in Slide-seqV2 gene expression in soma and neuropil following KA injection versus control PBS injection. (**F**) Metaplots showing activity-dependent changes in mRNA abundance in soma and neurites (top), transcription rates (middle), and mRNA half-life (bottom), as measured by metabolic labeling, for clusters defined in (D). Dark line = mean $\log_2$(FC) of all genes in the cluster, shading = ±SEM, dashed lines = mean in PBS-injected animals (top) or unstimulated neurons (middle, bottom) ± 1.2-FC.

#### Supplementary Figure 13

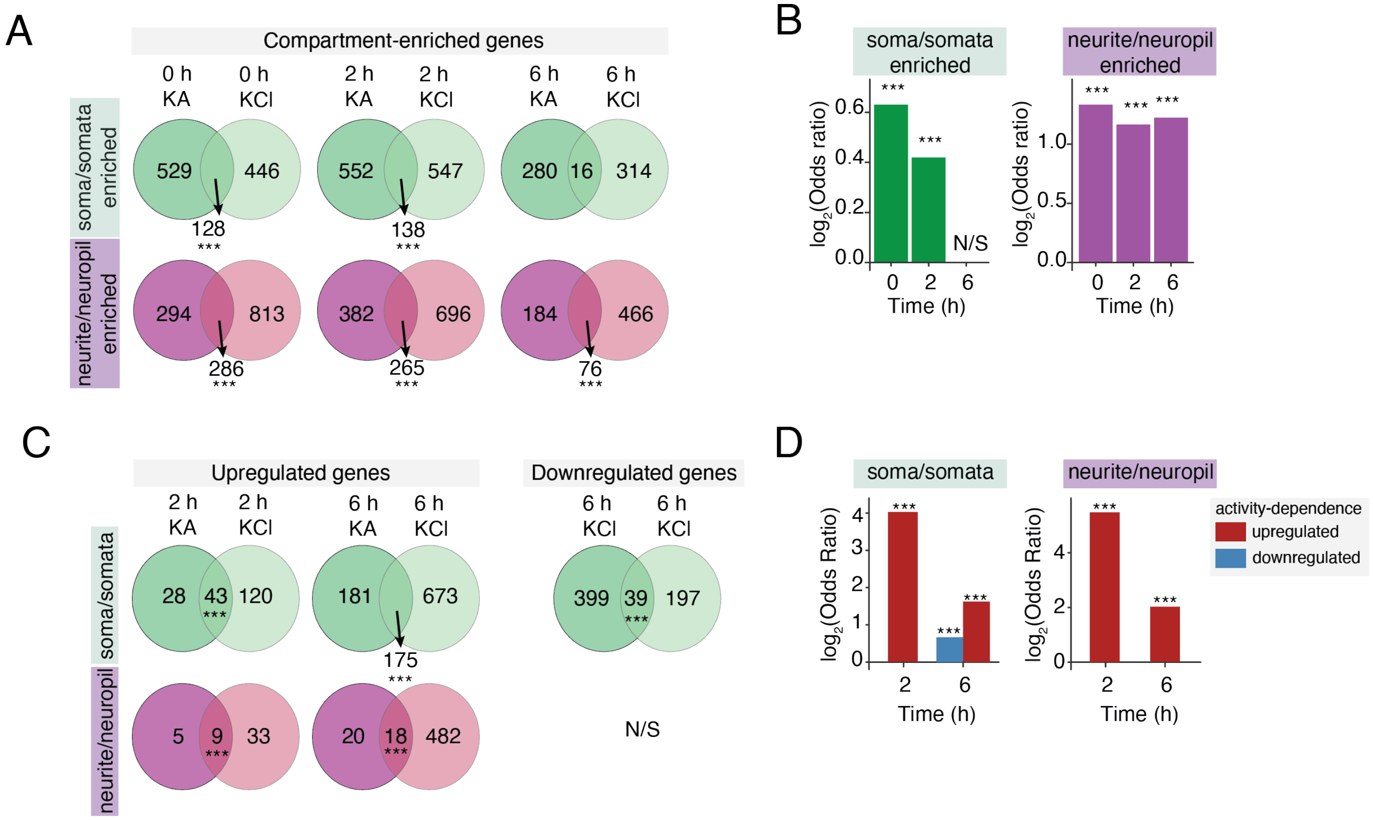


**Supplementary Figure 13: Overlap between Slide-seqV2 and separation chamber RNA-seq, Related to Figure 6.**

(**A**) Venn diagram of overlap between neurons in separation chambers and CA1 hippocampus for genes defined as enriched in neurites/neuropil (top) and soma/somata (bottom). (**B**) Log-odds ratio test of overlap between the two datasets in (A). $P_{adj}$ < 1e-4 (***), Chi-squared overlap test. (**C**) Venn diagram of overlap between activity-regulated genes in KCl-depolarized neurons in separation chambers and KA-stimulated CA1 hippocampus. (**D**) Log-odds ratio test of overlap between the two datasets in (C). $P_{adj} < 0.001$ (***), Chi-squared overlap test.

### Supplementary Figure 14

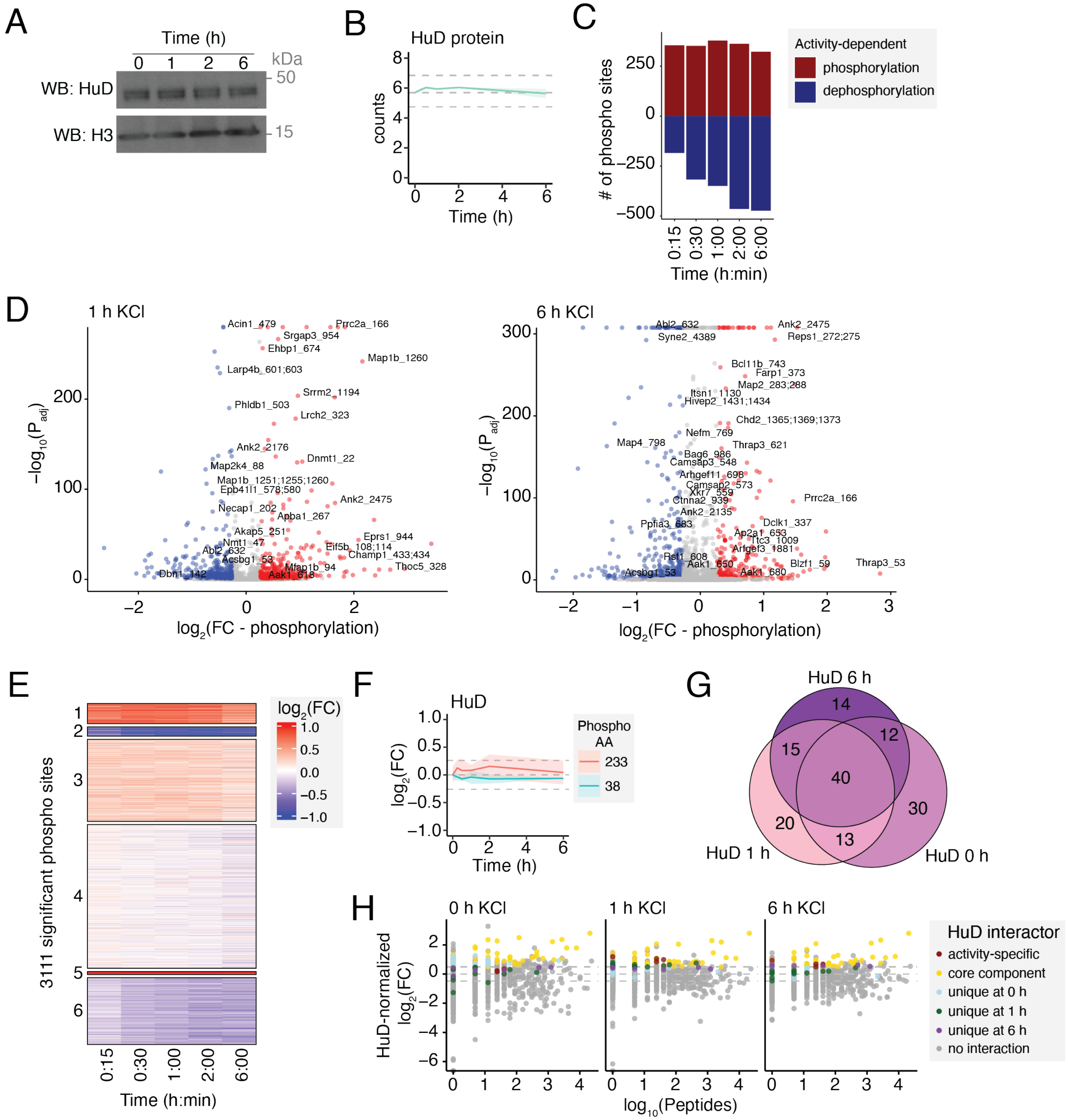


**Supplementary Figure 14: Activity-dependent HuD protein-protein interactions, Related to Figure 7.** (**A**) Representative western blot of HuD protein in primary neurons depolarized with KCl for 0, 1, 2, or 6 h, with histone H3 as a loading control. Experiment was repeated three times with similar results. Unprocessed blots are provided in the source data. (**B**) TMT-MS abundance of HuD protein in primary neurons depolarized with KCl for 0, 1, 2, or 6 h. Dark line = mean, shading = ±SEM, dashed lines = mean in unstimulated neurons ±1.2-FC. $n$=3 independent biological replicates. (**C**) Bar plot of significant (EdgeR Padj $<$ 0.05) differentially phosphorylated residues by TMT-MS phosphoproteomics at various time points following membrane depolarization. Numbers of activity-phosphorylated and -dephosphorylated residues are plotted as positive and negative values on the y-axis, respectively. $n$=3 independent biological replicates. (**D**) Volcano plots of −log10($P$adj) versus log2(FC in phosphorylation) at 1 h (left) and 6 h (right) versus 0 h KCl. (**E**) Heatmap of $\log_2$(FC) in phosphorylation for all residues with a significant activity-dependent change in phosphorylation in at least one time point relative to unstimulated neurons. (**F**) $\text{Log}_2$(FC) in phosphorylation for all detected HuD phosphorylated residues. Dark line = mean, shading = ±SEM, dashed lines = mean in unstimulated neurons ±1.2-FC. (**G**) Venn diagram of overlap between proteins detected as enriched for binding to HuD relative to GFP in primary neurons depolarized with KCl for 0, 1, or 6 h. (**H**) Scatterplot of $\log_{10}$(number of peptides) versus $\log_2$(FC) in HuD versus GFP IP-TMT-MS, color based on gene classification as a HuD-bound mRNA and on whether the corresponding protein represents a core or activity-specific interactor.

### Supplementary Figure 15

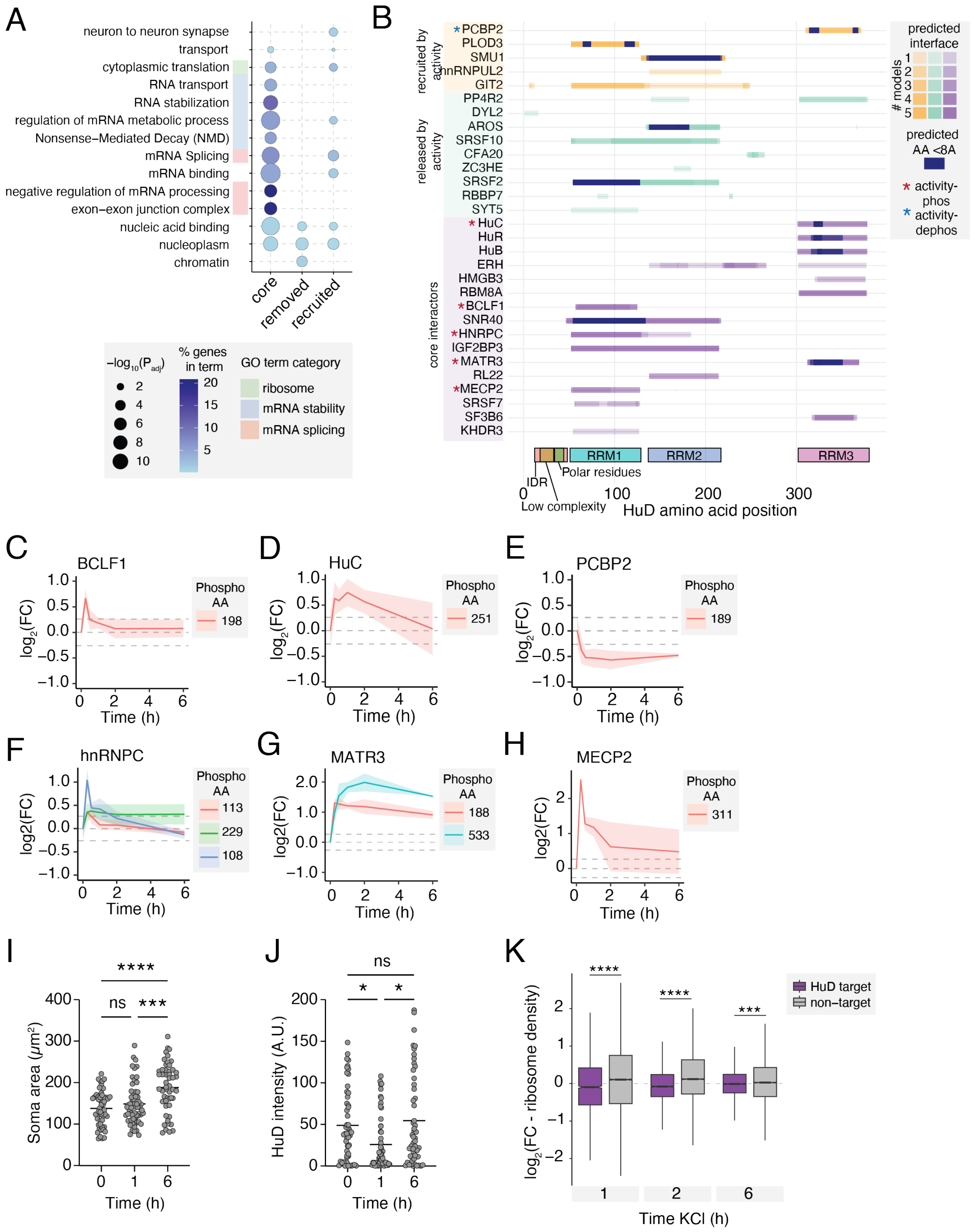

**Supplementary Figure 15: Activity-dependent HuD protein-protein interactions, Related to Figure 7.**
(**A**) Dot plot of the top GO terms (FDR < 0.05), separated based on HuD protein-protein interaction categories defined in Fig. 7B. (**B**) Bar plot of AlphaFold3 predicted protein-protein interaction interfaces with HuD (light blue) and high-confidence residues predicted within 8 Å (dark blue), separated based on interaction categories defined in Fig. 7B. Red stars indicate proteins that are phosphorylated in response to neuronal activity, and blue stars indicate proteins that are dephosphorylated in response to neuronal activity. (**C-H**) $Log_2$(FC) in activity-dependent phosphorylation sites for HuD protein-protein interactors in (B). Dark line = mean, shading = ±SEM, dashed lines = mean in unstimulated neurons ±1.2-FC. (**I**) Soma size per cell, $P_{adj}$ <0.0001 (****), $P_{adj}$ = 0.0006 (***). (**J**) Sum HuD intensity normalized to HuD area per cell, 0 vs 1 h $P_{adj}$ = 0.0131 (*), 0 vs 6 h $P_{adj}$ = 0.0112 (*). Nonparametric Kruskal-Wallis comparison with Dunn's multiple comparison correction, N=42-49 cells, *n*=2 biological replicates for both K and L. (**K**) Boxplot of $log_2$(FC)in ribosome density at 1, 2, or 6 h KCl versus unstimulated neurons, separated by HuD-bound mRNAs versus non-targets. **** = Padj < 0.0001, *** = Padj < 0.001 by Wilcoxon Test with BH correction.
